# Benchmarking methods for detecting differential states between conditions from multi-subject single-cell RNA-seq data

**DOI:** 10.1101/2022.02.16.480662

**Authors:** Sini Junttila, Johannes Smolander, Laura L Elo

**Author notes:** To whom correspondence should be addressed, **Contact information**: Turku Bioscience Centre, University of Turku and Åbo Akademi University, FI-20520 Turku, Finland. these authors contributed equally to this work (Sini Junttila, Johannes Smolander).

## Abstract

Single-cell RNA-sequencing (scRNA-seq) enables researchers to quantify transcriptomes of thousands of cells simultaneously and study transcriptomic changes between cells. scRNA-seq datasets increasingly include multi-subject, multi-condition experiments to investigate cell-type-specific differential states (DS) between conditions. This can be performed by first identifying the cell types in all the subjects and then by performing a DS analysis between the conditions within each cell type. Naïve single-cell DS analysis methods that treat cells statistically independent are subject to false positives in the presence of variation between biological replicates, an issue known as the pseudo-replicate bias. While several methods have already been introduced to carry out the statistical testing in multi-subject scRNA-seq analysis, comparisons that include all these methods are currently lacking. Here, we performed a comprehensive comparison of 18 methods for the identification of DS changes between conditions from multi-subject scRNA-seq data. Our results suggest that the pseudo-bulk methods performed generally best. Both pseudo-bulks and mixed models that model the subjects as a random effect were superior compared with the naive single-cell methods that do not model the subjects in any way. While the naive models achieved higher sensitivity than the pseudo-bulk methods and the mixed models, they were subject to a high number of false positives. In addition, accounting for subjects through latent variable modeling did not improve the performance of the naive methods.

## 1 Introduction

Single-cell RNA-sequencing (scRNA-seq) can be used to quantify transcriptomes of thousands of single cells simultaneously. scRNA-seq experiments comprise multi-subject, multi-condition setups, in which each condition includes samples collected from multiple subjects, cell lines or other biological replicates, and the researchers want to investigate transcriptomic changes between the conditions. Obtaining a large enough number of samples is crucial to ensure that the discovered gene markers are prevalent in the subject groups or treatment conditions, and not only in single subjects or biological replicates.

The analysis workflow of multi-subject, multi-condition scRNA-seq data involves steps that are the same as in any scRNA-seq analysis. Quality control is important to remove poor-quality cells, such as doublets, empty droplets and dead cells [1]. Normalization aims to make the gene expression profiles of different cells more comparable by decreasing the technical bias caused by the library size and other confounding factors [2]. In cell type identification, each cell is given an identity from the cell types that are present in the tissue. Data integration methods can be used to automate the identification of the same cell types across the samples [3,4].

Once the cell types have been confidently identified from all the samples, the next step is to perform differential state (DS) analysis between two or more conditions within each cell type separately. DS changes can be divided into several subtypes [5], including changes in the mean expression, which is commonly known as differential expression (DE). The other DS types model more subtle transcriptomic differences, such as the proportion of highly and lowly expressed cell populations. While virtually all methods have been designed to detect only changes in the average expression, single-cell method developers have recently started to pay attention to the other DS types as well [6,7].

The classical statistical tests for DS testing in scRNA-seq data, such as the Wilcoxon rank-sum test, naively assume the samples are statistically independent. However, this is usually not the case in multi-subject scRNA-seq data, where cells from the same subject often have more similar gene expression profiles, which causes an error in the statistical testing known as the pseudoreplicate bias [8]. To alleviate the pseudoreplicate bias, two approaches currently exist. The first approach is to use mixed models that model subjects as a random effect. The second approach is the pseudo-bulk aggregation, which transforms scRNA-seq data into bulk-like data by aggregating gene counts within each cell type and subject. Both approaches have previously been shown to reduce the number of false positives [6,8–10].

Differential expression analysis in scRNA-seq data was first investigated in papers that did not address the issue of multi-subject setup [11,12]. Since then, a few papers have investigated the issue of multi-subject, multi-condition scRNA-seq differential expression analysis. However, there still remains a lack of consensus regarding the best approaches. The muscat simulator [6] was introduced to enable simulation of multi-subject, multi-condition data based on reference data, and it also allows to simulate other DS types with more subtle differences in addition to DE. The muscat R package also provides functions for several pseudo-bulk methods and mixed models.

A more recent paper by Zimmerman et al. [8] compared several off-the-shelf mixed models, pseudo-bulk methods and naïve methods that do not model the subjects in any way using a limited simulation setup. The simulation was based on plate-based data with dropouts and not droplet data, such as Chromium [13], which is currently the most popular scRNA-seq protocol and generally not considered zero-inflated [14]. The authors recommended a mixed model based on the MAST statistical test [15] (MAST_RE) that accounts for the subjects as a random effect and claimed it was superior compared with the pseudo-bulk methods. Another recent paper by Squair et al. [10] compared naïve methods, pseudo-bulk methods and one mixed model method (muscat_MM). Their comparison was not based on a simulation but a comparison between paired scRNA-seq and bulk RNA-seq data. The ground truth for the bulk data was defined using two of the bulk differential expression tests, which could cause significant bias to the results. The comparison did not consider the recently introduced MAST model (MAST_RE) [8] or NEBULA, which is another recently introduced mixed model specifically designed for the DS analysis of multi-subject scRNA-seq data [16].

To address the need for better understanding the relative performance of various naïve, pseudo-bulk, and mixed model methods, we compared 18 different methods for DS analysis of multi-subject scRNA-seq data. Our comparison included three mixed models (MAST_RE [8], muscat_MM [6] and NEBULA-LN [16]) that model subjects as a random effect, six pseudo-bulk methods (edgeR [17] and DESeq2 [18] with sum aggregation, Limma [19] and ROTS [20] with sum and mean aggregation), and five naïve methods (the popular Wilcoxon rank-sum test and four other methods from the Seurat R package [3]). Additionally, we tested four latent variable methods from the Seurat R package that can be used to account for variables such as batch effects in DS analysis. To compare the DS analysis methods, we first carried out a comprehensive simulation analysis based on two different simulation models. The performance was assessed using several gold standard performance metrics: area under the receiver operating characteristic curve (AUROC), sensitivity, specificity, and precision. Finally, we estimated the proportion of false positives by performing a mock comparison between random groups using real data.

## 2 Materials and methods

### 2.1 Methods for detecting differential states

In total, we considered 18 DS analysis methods in our comparison (**Table 1**). These methods belong to two broad categories: pseudo-bulk methods and single-cell methods. The pseudo-bulk methods aggregate count values from each sample and cell type (cluster) to create data that can be analyzed using the same methods as bulk RNA-seq data, maintaining the same number of genes but reducing the number of cells to the number of samples in the gene expression matrix. Single-cell methods assume that the data have been normalized at the single-cell level, and the DS analysis is carried out using the normalized data directly. The single-cell methods can be further divided into two sub-categories: mixed models and naive methods. The mixed models model the subjects as a random effect, whereas the naive models assume that all the cells are statistically independent and do not model the subjects in any way. In addition, we considered a third type of single-cell methods from the Seurat R package, the latent variable models, that test whether the difference in gene expression between the groups can be explained by the difference in one or multiple latent variables. These methods were designed to account for batch effects or other confounders in the data.

**Table 1.**
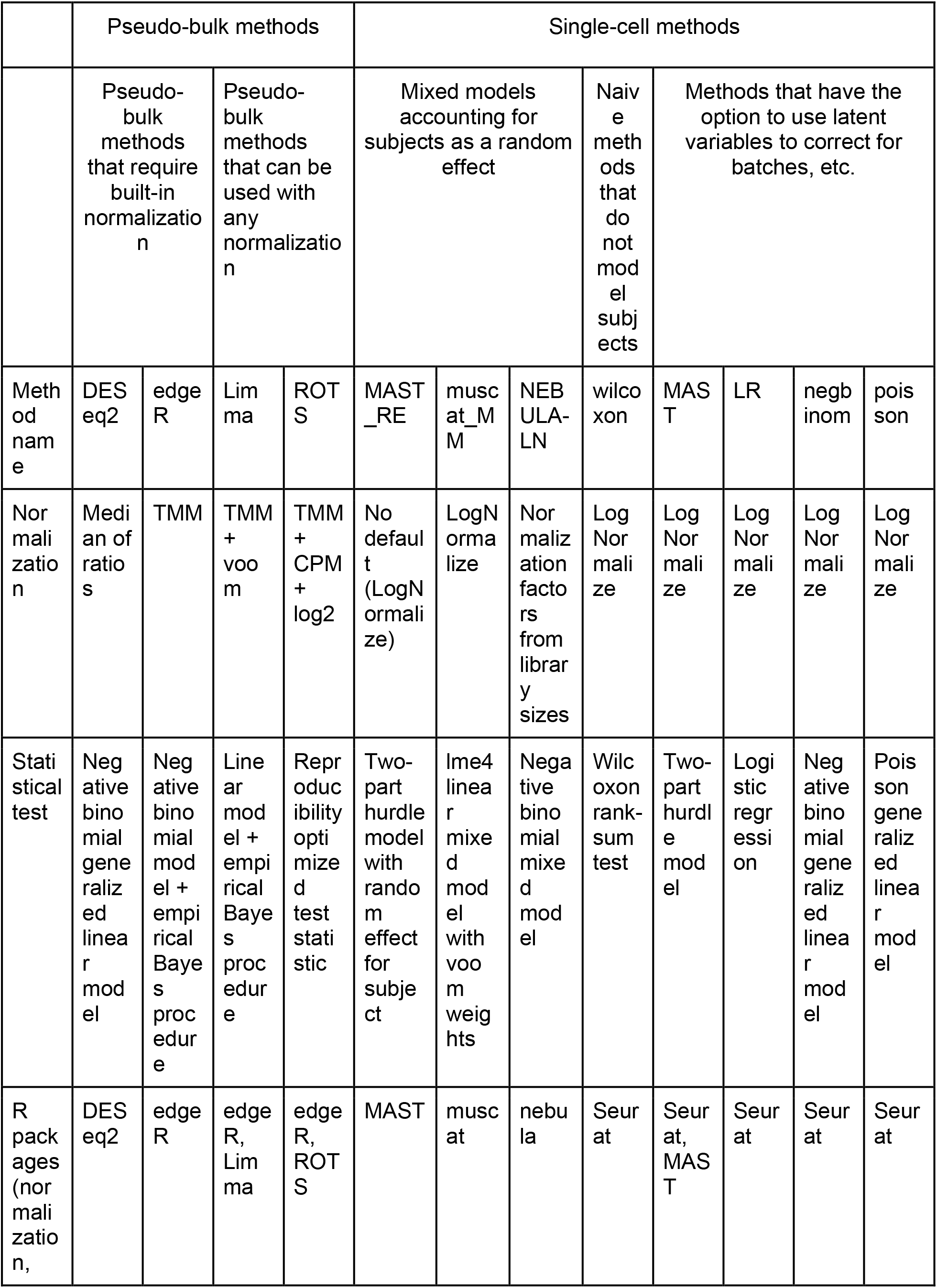
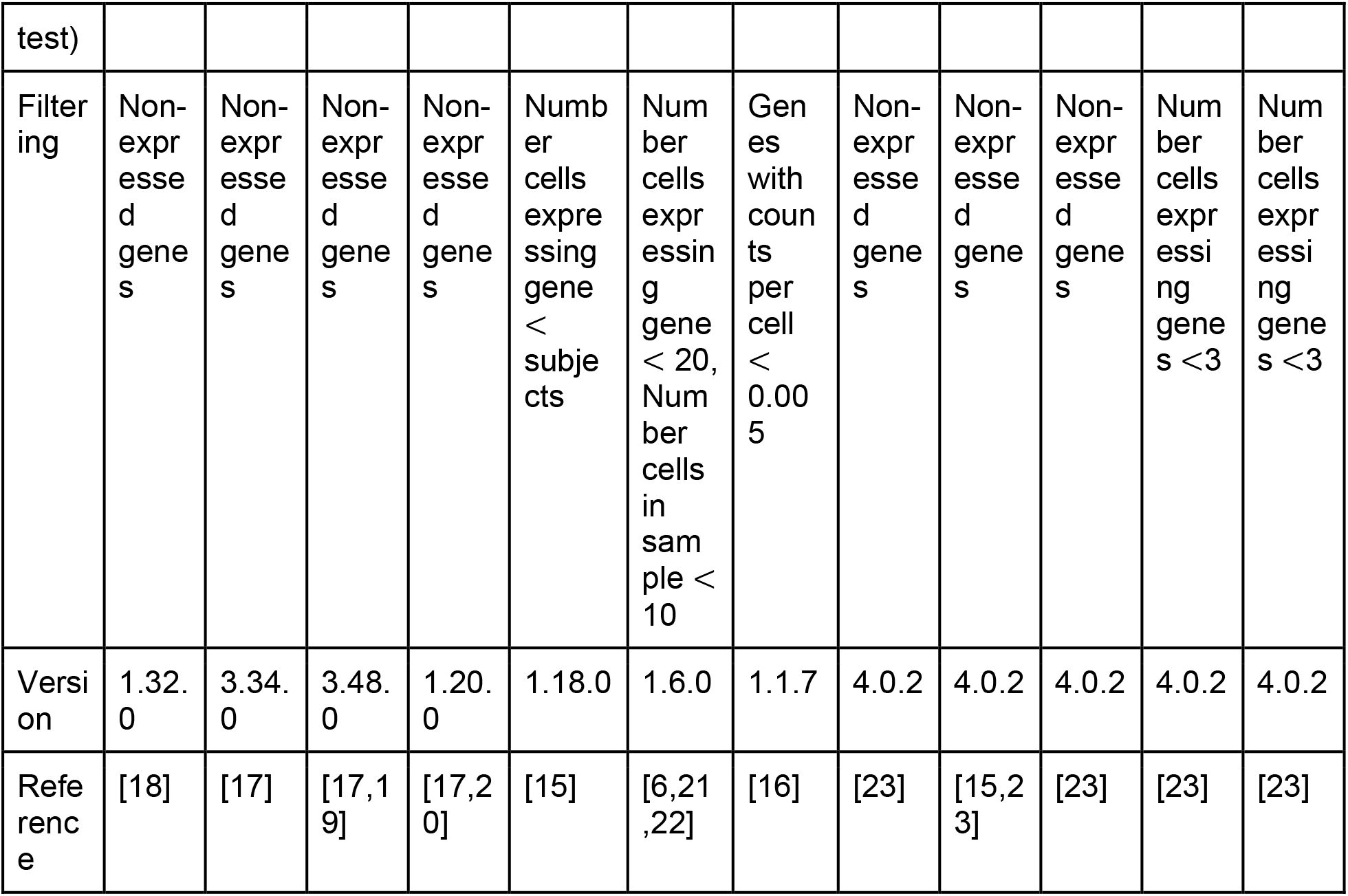
Details of the methods for the differential state analysis of scRNA-seq data compared in this study.

The aggregation of the count values for pseudo-bulk methods can be performed using two approaches: cumulative summing of raw count values (sum) or averaging single cell normalized count values (mean). The sum aggregation is followed by bulk normalization, and it has achieved better performances in earlier studies than the mean aggregation [6]. A recent study by Thurman et al. [9] recommended the sum aggregation with DEseq2 for multi-subject DS analysis, which is a popular statistical test for bulk RNA-seq DE analysis [18]. We selected DEseq2 and three other statistical tests, Limma, edgeR and ROTS [17,19,20] as a representation of the pseudo-bulk methods. In addition to performing the pseudo-bulk aggregation for all four statistical tests by the sum aggregation, we also tested the mean aggregation for two of the statistical tests (ROTS and Limma) that can be used with any normalization method. The sum and mean aggregated pseudo-bulk methods are denoted with _sum and _mean suffixes in the results, respectively.

Mixed models that account for the subjects as a random effect are gathering increasing interest. We included three mixed models in our comparison: MAST_RE, muscat_MM, and NEBULA-LN. A recent paper by Zimmerman et al. [8] recommended for multi-subject DS analysis a MAST model (MAST_RE) [15] that models the subjects as a random effect. The muscat R package includes a mixed model (muscat_MM), which uses the lme4 linear mixed model with voom weights [6,21,22]. NEBULA-LN is a recently introduced negative binomial mixed model designed for fast, multi-subject DS analysis and estimation of co-expression between genes [16].

Seurat is a popular R package for scRNA-seq data analysis, including a wide array of statistical tests for DS analysis [3,23]. These include naive methods that do not model the subject in any way, such as the Wilcoxon rank-sum test, as well as models that can be used with “latent variables” to account for different confounding factors during the statistical testing. The way in which the latent variable modeling is performed varies depending on the statistical test. The batch effect is the only confounder that is mentioned in the documentation, but the user can include an arbitrary number of latent variables in the *FindVariables* function. The four statistical tests of Seurat that support the use of latent variables are MAST, logistic regression, negative binomial generalized linear model (negbinom), and poisson generalized linear model (poisson). We included these four tests and their naive versions in our comparison. In addition, we included the Wilcoxon rank-sum test, which is the default method for DS analysis in Seurat. Other approaches for performing multi-subject DS analysis, such as mixed models with random effects or pseudo-bulk methods, are not currently available in Seurat (version 4.1)

### 2.2 Simulation of scRNA-seq data

Since it is in practice very difficult to ascertain which genes are differentially expressed between conditions in real scRNA-seq data, simulation is necessary to obtain an accurate benchmark. To simulate scRNA-seq data, we used two different approaches. The first approach is based on a reference-free negative binomial generative model presented in the original study of one of the benchmarked tools (NEBULA). This approach can simulate DE and non-DE genes by controlling the average fold-change between the groups but not other DS types. The second approach uses muscat, which is a recently introduced R package based on a reference-based negative binomial generative model that enables simulating multi-subject, multi-condition scRNA-seq data using real data as reference [6]. It can simulate genes of four different DS types and two non-DS types: changes in the mean expression (DE), the proportions of low and high expression components (DP), differential modality (DM), both proportions and modality (DB), equivalent expression (EE) and expression at low and high components by an equal proportion (EP).

#### 2.2.1 Simulation using a reference-free negative binomial generative model

We performed a reference-free negative binomial generative model simulation using the approach from the original paper of one of the benchmarked tools (NEBULA-LN) [16]. This simulation allowed tuning the model parameters, including two overdispersion parameters (cell and sample) that create random variation into the gene expression levels between cells and samples, and the average number of cells per sample.

To generate gene expression data that included both non-DE and DE genes, we made changes to the original NEBULA simulation. As in the original simulation, we simulated non-DE genes by setting logFC=0. Additionally, we simulated DE genes with logFC between 0.5 and 2.0. In total, our simulation included 1280 datasets, each containing 100 DE genes and 1900 non-DE genes. We simulated the 1280 datasets by adjusting five different parameters: the number of samples (6,8,10,12,14,16,18,20,30,40), the average number of cells per sample (100,500,1000,2000), the distribution for sampling the average number of cells (Poisson, negative binomial), cell overdispersion (0.05, 0.10, 0.20, 0.50), and sample overdispersion (0.1, 1, 10, 100). The average expression term in the generative model ranged from −4 to 2.

#### 2.2.2 Simulation using a reference-based negative binomial generative model

In our simulation with muscat, we considered reference data from four studies (Kang [24], Kallionpää [25], Thurman [9] and Liu [26]), which are summarized in **Table 2**. The Kang dataset comprises peripheral blood mononuclear cells (PBMC) from lupus patients before and after treatment with interferon-β. The Kallionpää dataset includes PBMC cells from children that developed type I diabetes at a young age along with paired control samples. The Thurman dataset includes cells segregated from large and small airway surface epithelium of newborn cystic fibrosis (CF) and non-CF pigs. The Liu dataset includes PBMC cells from COVID-19 patients, patients with tropical infectious diseases, and healthy subjects.

**Table 2.** Details of the reference datasets used in the simulation using a reference-based negative binomial generative model.

|  | Kang | Kallionpää | Liu | Thurman |
| --- | --- | --- | --- | --- |
| Number of replicates in simulation | 8, 16 | 12 | 12, 16, 20 | 8 |
| Tissue type | PBMC | PBMC | PBMC | airway surface epithelium |
| Conditions | IFN-β-treated vs. non-treated | T1D cases vs. matched controls | COVID-19 vs. healthy controls | CF vs. non-CF |
| Organism | human | human | human | pig |
| Number of control samples in original dataset | 8 | 4 | 13 | 4 |
| Reference | [24] | [25] | [26] | [9] |

For each simulated dataset, we simulated three clusters with varying magnitudes of differences. 10% of the genes in each cluster were assigned a differential distribution (2.5% for each of the four differential distributions DE, DP, DM and DB). The relative log-fold-change (logFC) values were set to 0.5, 1 and 1.25 for clusters 1, 2 and 3, respectively. Using Kang data as reference, four datasets were simulated: 20,000 cells and four replicates per condition, 20,000 cells and eight replicates per condition, 5,000 cells and four replicates per condition, 5,000 cells and eight replicates per condition. One dataset was simulated using Kallionpää data as reference: 7,500 cells and four replicates per condition. Using Liu data as reference, three datasets were simulated: 12,000 cells and six replicates per condition, 16,000 cells and eight replicates per condition, 20,000 cells and ten replicates per condition. One dataset was simulated using Thurman data as reference: 20,000 cells and four replicates per condition. Additionally, to investigate the impact of the number of cells and the number of samples on the performance, we extended the muscat simulation for the Liu dataset so that it included more variation in the number of cells per sample (500, 1000, 2000, 4000) and the number of subjects (8, 12, 16, 20, 24, 28, 32, 36, 40).

For Kang, Liu and Thurman reference data, we used the cell type annotation that was provided by the authors of the original studies in the muscat simulation. For Kallionpää data, we performed Seurat integration (v 4.0.3) with the default parameter values and used the resulting clustering in the muscat simulation.

#### 2.2.3 Simulation of imbalanced distribution of cells across the samples

In both simulations, the datasets contained an almost even distribution for the number of cells between subjects. However, this assumption is not valid in many real situations, and a recent paper by Zimmerman et al. [8] suggested that especially the performance of pseudo-bulk methods deteriorates when a dataset has an uneven distribution for the number of cells. Therefore, we also simulated clusters that had large variation in the number of cells between the samples. In the reference-free simulation, the random sampling of the number of cells was performed using two statistical distributions: Poisson for balanced distribution and negative binomial for imbalanced distribution. In the reference-based simulation, we randomly subsampled cells for all simulated datasets without replacement so that the proportion of remaining cells in the samples varied with even intervals from 0.20 to 1. The subsampling was performed for each of the clusters separately and the proportions of remaining cells were chosen randomly for the samples.

### 2.3 Performance evaluation

We performed Receiver Operating Characteristic (ROC) curve analysis on the simulation results using pROC R package [27]. As the predictor we used the p-values and as the response the ground truth provided by the simulation on which genes had differential states. Since the methods had different gene filtering strategies (see **Table 1**), we only considered genes that were included by all methods.

While the AUROC is useful for assessing the performance so that the evaluation is not constrained to a specific p-value threshold, and it can be interpreted as measuring the accuracy of ranking positive genes higher than negatives, it is possible to achieve a perfect AUROC score with statistically insignificant p-values. To assess the ability of the methods to provide well-calibrated p-values, we also calculated the sensitivity, specificity and precision of the methods using the false discovery rate (FDR) of 0.05 as a cut-off. Before adjusting the p-values for multiple comparisons, we excluded the genes that were not included by all the methods (see **Table 1**).

### 2.4 Mock comparison using real data to estimate the proportion of false positives

To estimate the proportion of false positives, we performed a mock analysis using a real scRNA-seq dataset that includes PBMCs from healthy subjects, patients with flu or COVID-19 [26]. We took the 13 healthy control samples and used the metadata stored in the publicly available Seurat object (GEO accession GSE161918) to extract B cells that were labeled as singlets and had at maximum 10% mitochondrial reads. We randomly assigned one of the two mock groups for each sample and performed statistical testing between the mock groups using each of the 18 methods to determine the DS genes. A gene was considered significant if FDR ≤ 0.05. Before adjusting the p-values for multiple comparisons, we excluded the genes that were not included by all the methods (see **Table 1**). We performed the random mock group assignment 30 times using different random seeds.

## 3 Results

### 3.1 Simulation based on a reference-free negative binomial generative model

We simulated data based on a reference-free negative binomial generative model from the original paper of the NEBULA method (see **Section 2.2.2**). To benchmark the methods in a way that is not limited to a single p-value cutoff, we calculated the AUROC for each method and cluster (see **Section 2.3**). The AUROC values were, on average, highest for the pseudo-bulk methods, followed by the naïve and latent variable methods (**Fig. 1a**). The number of cells and samples did not have a noticeable impact on the superiority of the method types. (**Supplementary Fig. 1**).

**Figure 1.**
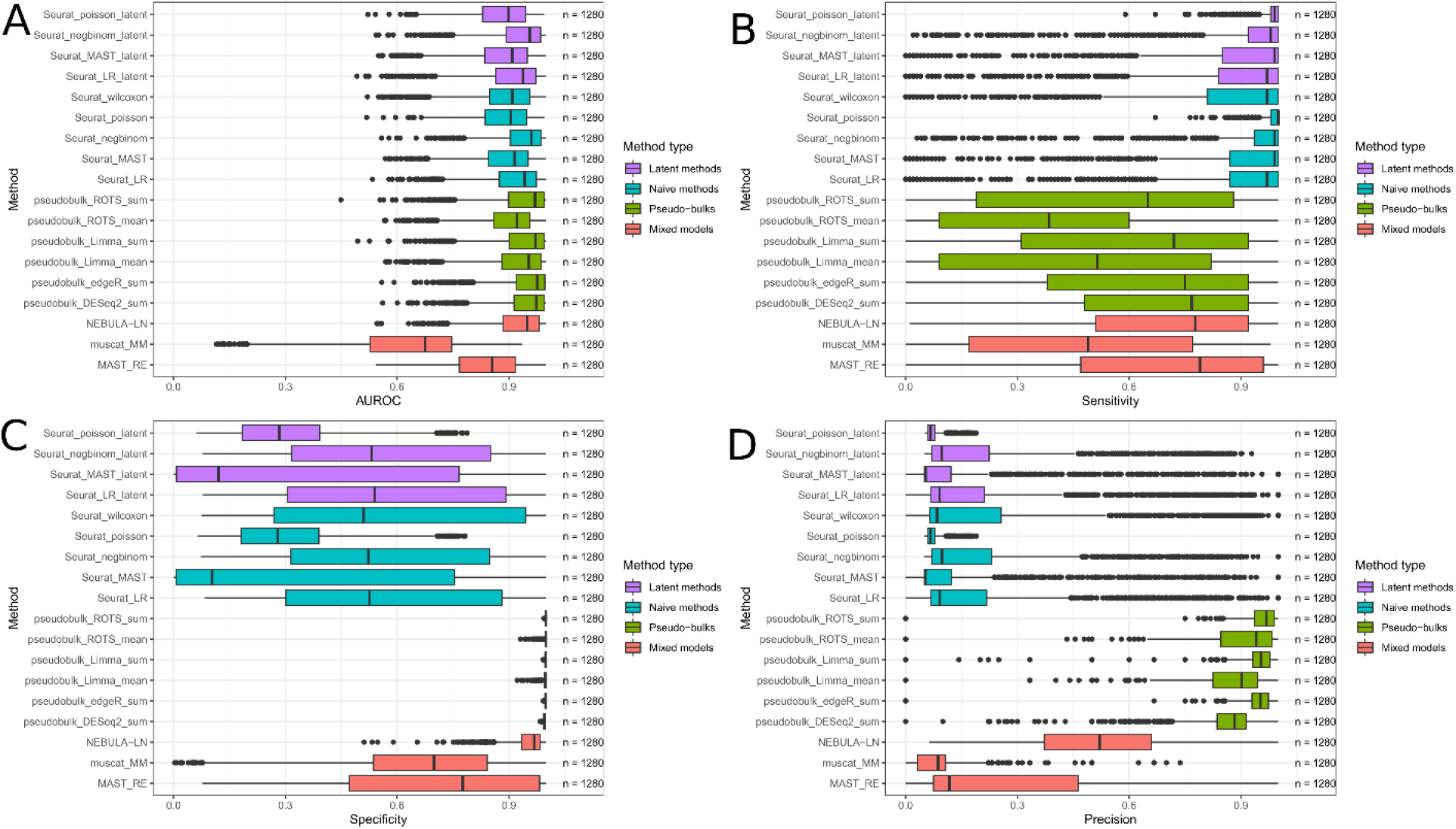
Results of the simulation based on a reference-free negative binomial generative model. Each boxplot shows values for 1280 simulated datasets with varying data properties.

In addition to the AUROC, we calculated the sensitivity, specificity and precision using FDR of 0.05 as a cutoff to define the positives and negatives. Overall, the sensitivity was higher for the naive methods and the latent methods compared to the pseudo-bulk methods and the mixed models (**Fig. 1b**), and it increased when the number of samples increased with all the methods, as expected (**Supplementary Fig. 2**). However, the pseudo-bulk methods generally provided significantly better precision and specificity compared to all other method types (**Fig. 1c-d, Supplementary Fig. 3-4**). With Limma and ROTS we also tested the effect of the aggregation method on the results, suggesting systematically better performance of the sum over the mean aggregation (**Fig. 1**, **Supplementary Fig. 1-4**). **Supplementary Figures 5-22** show the results of **Supplementary Figures 1-4** for each method separately.

Finally, we investigated how the imbalance in the number of cells between the samples affected the performance, which indicated that in these data the differences were relatively small for all methods (**Supplementary Fig. 23**).

### 3.2 Simulation based on a reference-based negative binomial generative model

We used muscat R package to simulate scRNA-seq data using data from four different studies (Kang, Kallionpää, Thurman, Liu; see **Section 2.2.2**), to study the effects of different DS types, including changes in the mean expression (DE), changes in the proportions of low and high expression-state components (DP), changes in modality (DM), and changes in both proportions and modality (DB). In total, 54 cell populations (clusters) were used in the benchmarking.

We first calculated the AUROC for each method and cluster and grouped the results by the DS type (**Fig. 2a**). These results indicate that the DS type did not have a notable impact on the ranking of the methods. Unsurprisingly, the performance scores for the DE type were consistently higher than for the three other DS types, which contained more subtle transcriptomic differences between the groups than the DE genes. The pseudo-bulk methods and the naive methods achieved higher performance than the latent models and the mixed models. The latent models were clearly the weakest-performing models. The mixed models had considerable variation between their performances: MAST_RE achieved slightly better overall performance compared with NEBULA-LN, whereas muscat_MM was inferior with all four data types. Again, the pseudo-bulk aggregation by summing performed generally better than the mean aggregation.

**Figure 2.**
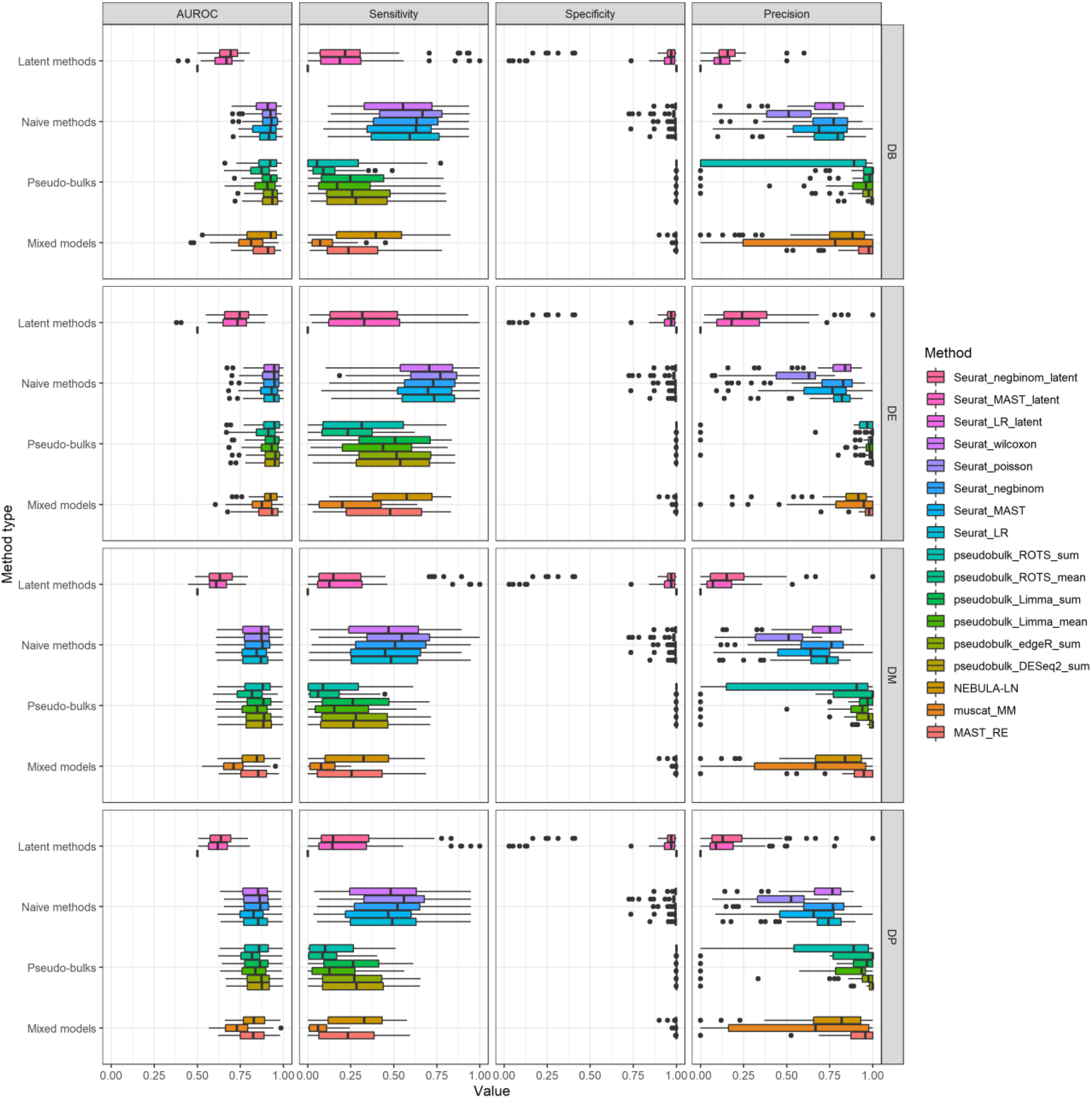
Results of the simulation based on a reference-based negative binomial generative model. Each box plot includes performance values for 54 cell populations (clusters). The rows signify the four different differential states: changes in both proportions and modality (DB), changes in the mean expression (DE), differential modality (DM), and changes in the proportions of low and high expression-state components (DP). The columns group the results by the four different performance metrics: Area Under the Receiver Operating Characteristic (AUROC) curve, sensitivity, specificity, and precision. Seurat_poisson_latent was left out from the results due to its high failure rate for the simulation.

In addition to the AUROC, we calculated the sensitivity, specificity and precision using FDR of 0.05 as a cutoff to define the positives and negatives (**Fig. 2b**). The results suggested that the naive methods provided the best sensitivity among the method types, followed by the latent variable models. However, their specificity and precision were worse compared to the pseudo-bulk methods and the mixed models (**Fig. 2c-d**). In other words, the pseudo-bulk methods and the mixed models were able to effectively minimize the number of false positives, whereas a significant proportion of the findings, generally 25%, found by the naive methods and the latent variable models were false. Overall, the precision and specificity were higher for the pseudo-bulk methods than the mixed models. The overall results remained similar when investigating the impact of the number of cells and samples on the performance (**Supplementary Figures 24-56**).

To take a closer look at the two best-performing mixed models (NEBULA-LN and MAST_RE) and the four pseudo-bulk methods that use the sum aggregation, we studied the genes that behaved differently between the six methods (**Fig. 3**). We defined a metric called overlap, which was calculated by counting in each group how many pseudo-bulk normalized data points (samples) were within the range of the values of the other group after which we divided the counts by the number of samples in one group, and then took their average. When investigating the overlaps of the gene-wise distributions between the sample groups, the false positives of the pseudo-bulk methods had a small overlap (from 0 to 35%) compared to true negatives (on average above 50%), suggesting that the false positives of the pseudo-bulk methods occurred due to errors in normalization or simulation (**Fig. 3a**). The false positives of the mixed models did not show such a trend, but their overlap values were similar to those of the true negatives (**Fig. 3a**). The fold-changes indicated that the false positives of the pseudo-bulk methods also had a higher fold-change than the mixed models or the true negatives and they were comparable to the true positives (**Fig. 3b**). However, the false positives of the pseudo-bulk methods had generally lower average expression than the mixed models or the true positives (**Fig. 3c-d**).

**Figure 3.**
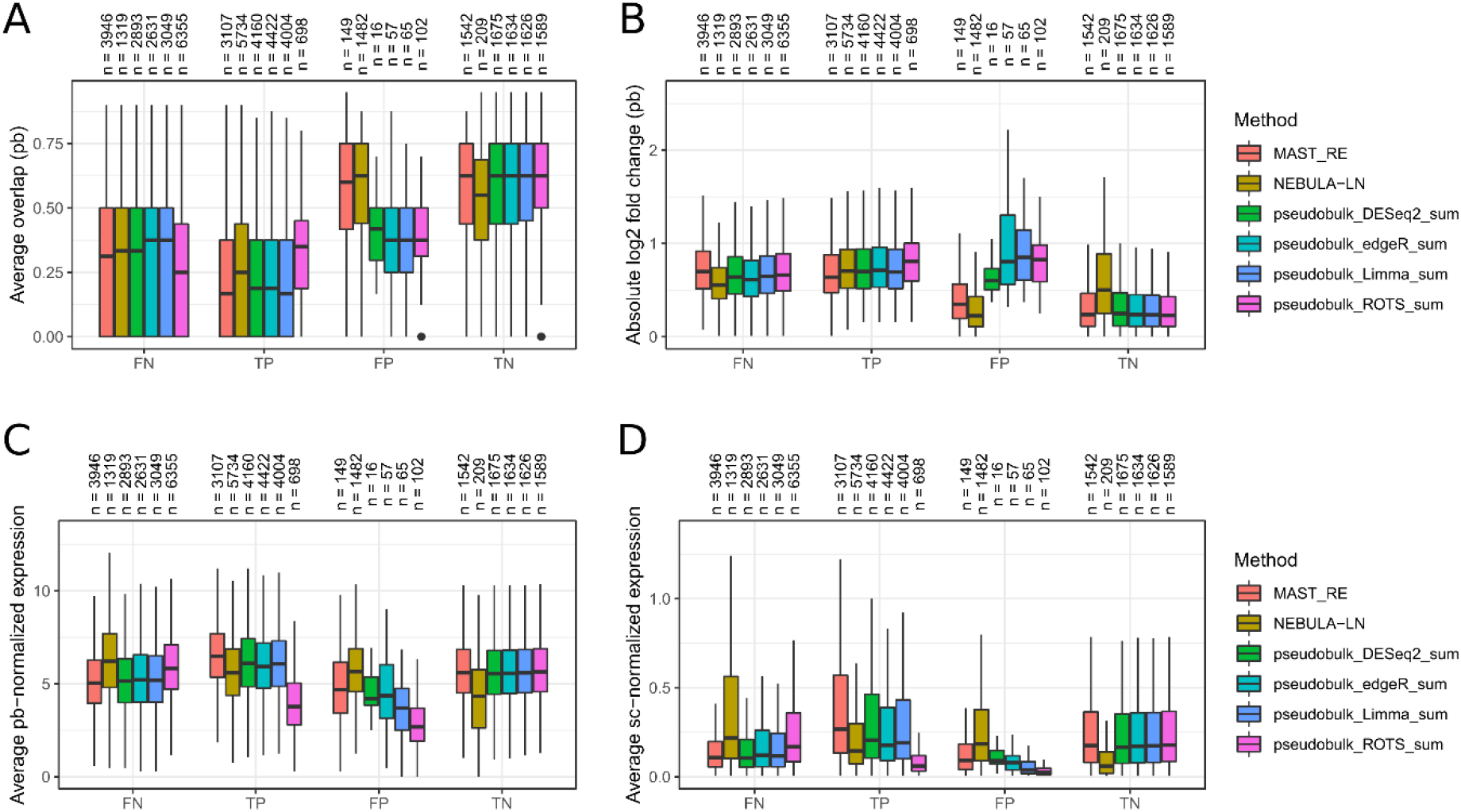
Analysis of the genes that were differently detected between the pseudo-bulk methods and two top-performing mixed models in the negative binomial simulation. **(a)** Average overlap was calculated by counting in each group how many pseudo-bulk (pb) normalized data points (samples) were within the range of the values of the other group, divided by the number of samples in the group, and then taking their average. **(b)** Absolute value of the log2-transformed fold-change calculated between the pseudo-bulk normalized gene expression values of the two groups. We added a pseudo-count value of 1 to each mean expression when calculating their fold change. **(c)** Average pseudo-bulk normalized gene expression. The pseudo-bulk normalization was performed using the normalization that was used for Limma and ROTS (see **Table 1**). **(d)** Average single-cell (sc) normalized gene expression. The singlecell normalization was performed using the normalization method that muscat simulator uses. To make the boxplots more readable, we removed the outliers for **(b-d)**.

Finally, we investigated how the imbalance of the number of cells affected the performance (**Supplementary Fig. 57**). In general, the AUROC values of the methods were slightly lower in the imbalanced datasets, but especially the sensitivity of the methods decreased in the imbalanced datasets.

### 3.3 Mock comparison with real data

To further estimate the proportion of false positives for each method in a real experimental setting, we carried out a mock comparison by randomly dividing 13 healthy subjects from a COVID-19 study into two mock groups. The results suggest that the naive methods that did not account for subjects in any way and the latent methods were subject to a high number of false positives (**Fig. 4**). By contrast, the pseudo-bulk methods and the mixed models generally produced small numbers of false positives. This is in accordance with the simulation results in **Section 3.1**. The logistic regression model of Seurat, which models the subjects as a latent variable, did not find any false positives, but this is likely due to the method’s inability to produce any findings for some data types, like the negative binomial simulation (see **Section 3.1**).

**Figure 4.**
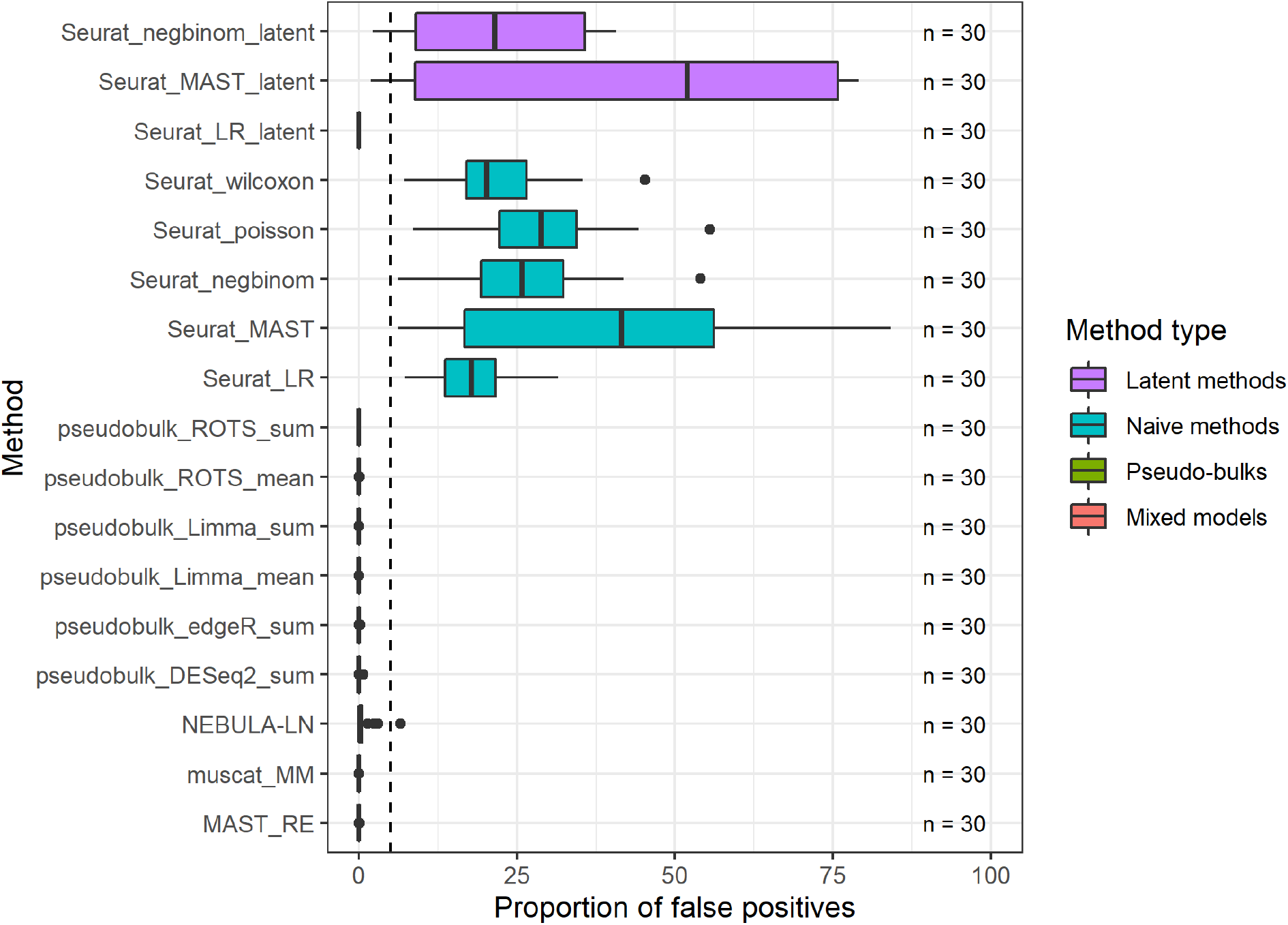
Mock analysis using real data to estimate the proportion of false positives. The mock analysis was performed by segregating good-quality B cells from a COVID-19 dataset [26] that consists of 13 healthy control subjects and by randomly assigning one of the two mock groups for each subject. The assumption is that no genes with differential states should be found between the random mock groups. The random sampling was repeated 30 times. Seurat_poisson_latent was left out from the results due to its high failure rate (29/30) for the mock data. The dashed vertical line at 5% denotes the expected maximum proportion of false positives with an FDR threshold of 0.05.

## 4 Discussion

Finding differential states between conditions from scRNA-seq data involves performing statistical testing between two or more groups of cells for each cell type separately. scRNA-seq experiments increasingly include multiple subjects or biological replicates to confirm that the transcriptomic changes are prevalent in groups and not only in single subjects. This requires specialized tools due to the hierarchical structure of the data. Cells from the same subject often have more similar gene expression profiles, which violates the statistical independence assumption of the basic statistical tests.

This issue has been already addressed in recently published papers that have proposed new, improved methods for the DS analysis of multi-subject scRNA-seq data [6,8,9,16]. The two approaches that currently seem most promising are the pseudo-bulk methods that aggregate counts from each cluster and sample, and the mixed models that model the subjects as a random effect. They have both been demonstrated to decrease the number of false positives compared to naïve single-cell DS analysis methods [8,10]. However, no attempts have yet been made to compare these tools in the same work.

In this paper, we compared 18 tools for the DS analysis of multi-subject scRNA-seq data. These methods included both pseudo-bulk methods and mixed models, but also naive single-cell methods that do not model the subjects in any way, as well as methods that model the subjects as a latent variable. Our benchmarking framework included both simulated and real data. For the simulation, we considered both a reference-free negative binomial generative model and a reference-based negative binomial generative model. The reference-free generative model simulated genes with changes in the mean expression (DE). The reference-based negative binomial generative model simulation was performed using the muscat R package, which is currently the only simulator that can simulate multi-subject scRNA-seq data with four different DS types. Finally, we performed a mock comparison using 13 healthy control subjects from a large COVID-19 dataset [26], which enabled us to estimate the proportion of false positives for each method.

The results indicated that the naive methods were indeed subject to a higher rate of false positives than the pseudo-bulk methods and the mixed models. This conclusion is supported by all our analyses. While the naive models generally provided higher sensitivity, this benefit was negated by the lower precision and specificity. In other words, the naive models reported a lot of findings, but a large proportion of these were false positives. Although the AUROC results suggested that the p-values of the naive methods accurately ranked the positives before the negatives, the main issue was that the p-values were poorly calibrated. With FDR of 0.05 as a cutoff to define the positives and negatives, the methods can be expected to find at most 5% false positives from all positives. The naive methods found high proportions of false positives, up to 40% in the mock comparison and the simulations. In the simulation, the precision values of the pseudo-bulk methods were in better accordance with the FDR cut-off than the precision values of the mixed models.

We observed notable variation in the performances of the mixed models. Of the three mixed models that we considered in our comparison, MAST_RE and NEBULA-LN achieved considerably better overall performance than muscat_MM. However, the performances of the pseudo-bulk methods were mostly similar. Our results suggested that the pseudo-bulk aggregation by calculating the mean of single-cell normalized data provided inferior performance compared to the sum approach that cumulatively sums the count values and then uses bulk RNA-seq normalization. This is in line with at least one previous study that found that the sum aggregation outperformed the mean aggregation [6].

We investigated how the number of cells and samples affected the performance of the methods. An earlier study [8] found that the pseudo-bulks performed worse than the mixed models when the number of samples was small. The same paper also suggested that the pseudo-bulks would perform worse than the best mixed model (MAST_RE) when the samples have an uneven distribution for the number of cells. However, we were unable to validate these findings.

The popular Seurat pipeline includes four statistical tests that allows to incorporate several latent variables in the models. The latent models test if the observed differential expression change between the conditions can be explained by the difference in one or several variables. According to the package manual, this is recommended if the data contains batch effects in the DS analysis, but no instructions are provided for any other variables. As of Seurat v4.1, using latent variables is currently still the only way to account for the subjects in the DS analysis with Seurat. Our results strongly suggest against their use in the DS analysis when the subject is included as a latent variable. The latent models performed generally even worse than their naive counterparts. A recent study came to the same conclusion when they used ComBat to correct the data for the subject effect prior to the DS analysis [8,28]. However, the latent models might still be appropriate when modeling batch effects or other variables as latent variables.

To conclude, we performed a comprehensive comparison to benchmark 18 methods for DS analysis of multi-subject scRNA-seq data. Our results suggest that the pseudo-bulk methods and the mixed models that model subjects as a random effect were superior compared to the naive single-cell methods that do not model the subjects in any way. We also recommend not to perform DS analysis using Seurat’s statistical tests so that the subjects are modeled as a latent variable. Overall, the pseudo-bulk methods outperformed the mixed models. If the user wants to achieve high specificity and precision at the risk of losing some true positives, we recommend the pseudo-bulk ROTS with the sum aggregation. If sensitivity is more important than the false positive results, we recommend the pseudo-bulk methods Limma, DESeq2, or edgeR combined with the sum aggregation. We recommend that scRNA-seq analysis pipeline developers should begin to include pseudo-bulk methods and mixed models in their pipelines. To facilitate DS analysis of multi-subject scRNA-seq data, the codes that implement all the methods in this paper are freely available online (https://github.com/elolab/multisubjectDSanalysis).

## Key Points

- Naive single-cell DS analysis methods are subject to high proportions of false positives when analysing data with multiple biological replicates
- Latent variable models are not effective in reducing false discoveries when accounting for biological replicates
- Pseudo-bulk methods and mixed models that account for biological replicates as a random effect are effective in reducing false discoveries
- Overall, our results suggest that pseudo-bulk methods outperform mixed models

## Acknowledgments

The authors thank the Elo lab for fruitful discussions and comments on the manuscript.

**Sini Junttila** is a postdoc at the Medical Bioinformatics Center at Turku Bioscience Centre. Her expertise is in analysis of scRNA-seq and epigenomics data.

**Johannes Smolander** is a PhD student at the University of Turku specializing on single-cell RNA-seq data analysis. He works at the Medical Bioinformatics Center at Turku Bioscience Centre.

**Laura L. Elo** is Professor of Computational Medicine and Head of Medical Bioinformatics Centre, University of Turku, Finland. Her main research interests include computational biomedicine and bioinformatics.

## Contributions

LLE and SJ conceived the study. LLE, SJ and JS designed the study. JS and SJ implemented the study. JS wrote the manuscript. LLE and SJ commented the manuscript. LLE supervised the study. SJ and JS share the first authorship.

## Availability of data and materials

Codes for the 18 DS analysis methods are available at https://github.com/elolab/multisubjectDSanalysis,

**Supplementary data**

## References

1. Ilicic T, Kim JK, Kolodziejczyk AA, et al. Classification of low quality cells from single-cell RNA-seq data. Genome Biol. 2016; 17:29

2. Cole MB, Risso D, Wagner A, et al. Performance Assessment and Selection of Normalization Procedures for Single-Cell RNA-Seq. Cell Syst. 2019; 8:315–328.e8

3. Butler A, Hoffman P, Smibert P, et al. Integrating single-cell transcriptomic data across different conditions, technologies, and species. Nat. Biotechnol. 2018; 36:411–420

4. Luecken MD, Büttner M, Chaichoompu K, et al. Benchmarking atlas-level data integration in single-cell genomics. Nat. Methods 2022; 19:41–50

5. Korthauer KD, Chu L-F, Newton MA, et al. A statistical approach for identifying differential distributions in single-cell RNA-seq experiments. Genome Biol. 2016; 17:222

6. Crowell HL, Soneson C, Germain P-L, et al. muscat detects subpopulation-specific state transitions from multi-sample multi-condition single-cell transcriptomics data. Nat. Commun. 2020; 11:6077

7. Tiberi S, Crowell HL, Weber LM, et al. distinct: a novel approach to differential distribution analyses. 2021; 2020.11.24.394213

8. Zimmerman KD, Espeland MA, Langefeld CD. A practical solution to pseudoreplication bias in single-cell studies. Nat. Commun. 2021; 12:738

9. Thurman AL, Ratcliff JA, Chimenti MS, et al. Differential gene expression analysis for multi-subject single-cell RNA-sequencing studies with aggregateBioVar. Bioinformatics 2021;

10. Squair JW, Gautier M, Kathe C, et al. Confronting false discoveries in single-cell differential expression. Nat. Commun. 2021; 12:5692

11. Jaakkola MK, Seyednasrollah F, Mehmood A, et al. Comparison of methods to detect differentially expressed genes between single-cell populations. Brief. Bioinform. 2017; 18:735–743

12. Soneson C, Robinson MD. Bias, robustness and scalability in single-cell differential expression analysis. Nat. Methods 2018; 15:255–261

13. Zheng GXY, Terry JM, Belgrader P, et al. Massively parallel digital transcriptional profiling of single cells. Nat. Commun. 2017; 8:14049

14. Svensson V. Droplet scRNA-seq is not zero-inflated. Nat. Biotechnol. 2020; 38:147–150

15. Finak G, McDavid A, Yajima M, et al. MAST: a flexible statistical framework for assessing transcriptional changes and characterizing heterogeneity in single-cell RNA sequencing data. Genome Biol. 2015; 16:278

16. He L, Davila-Velderrain J, Sumida TS, et al. NEBULA is a fast negative binomial mixed model for differential or co-expression analysis of large-scale multi-subject single-cell data. Commun. Biol. 2021; 4:1–17

17. Robinson MD, McCarthy DJ, Smyth GK. edgeR: a Bioconductor package for differential expression analysis of digital gene expression data. Bioinformatics 2010; 26:139–140

18. Love MI, Huber W, Anders S. Moderated estimation of fold change and dispersion for RNA-seq data with DESeq2. Genome Biol. 2014; 15:550

19. Ritchie ME, Phipson B, Wu D, et al. limma powers differential expression analyses for RNA-sequencing and microarray studies. Nucleic Acids Res. 2015; 43:e47–e47

20. Suomi T, Seyednasrollah F, Jaakkola MK, et al. ROTS: An R package for reproducibility-optimized statistical testing. PLOS Comput. Biol. 2017; 13:e1005562

21. Bates D, Mächler M, Bolker B, et al. Fitting Linear Mixed-Effects Models Using lme4. J. Stat. Softw. 2015; 67:1–48

22. Law CW, Chen Y, Shi W, et al. voom: precision weights unlock linear model analysis tools for RNA-seq read counts. Genome Biol. 2014; 15:R29

23. Hao Y, Hao S, Andersen-Nissen E, et al. Integrated analysis of multimodal single-cell data. Cell 2021; 0:

24. Kang HM, Subramaniam M, Targ S, et al. Multiplexed droplet single-cell RNA-sequencing using natural genetic variation. Nat. Biotechnol. 2018; 36:89–94

25. Kallionpää H, Somani J, Tuomela S, et al. Early Detection of Peripheral Blood Cell Signature in Children Developing β-Cell Autoimmunity at a Young Age. Diabetes 2019; 68:2024–2034

26. Liu C, Martins AJ, Lau WW, et al. Time-resolved systems immunology reveals a late juncture linked to fatal COVID-19. Cell 2021; 184:1836–1857.e22

27. Robin X, Turck N, Hainard A, et al. pROC: an open-source package for R and S+ to analyze and compare ROC curves. BMC Bioinformatics 2011; 12:77

28. Johnson WE, Li C, Rabinovic A. Adjusting batch effects in microarray expression data using empirical Bayes methods. Biostatistics 2007; 8:118–127

