## Supplementary File for "Benchmarking methods for detecting differential states between conditions from multi-subject single-cell RNA-seq data"

### Supplementary Figure 1

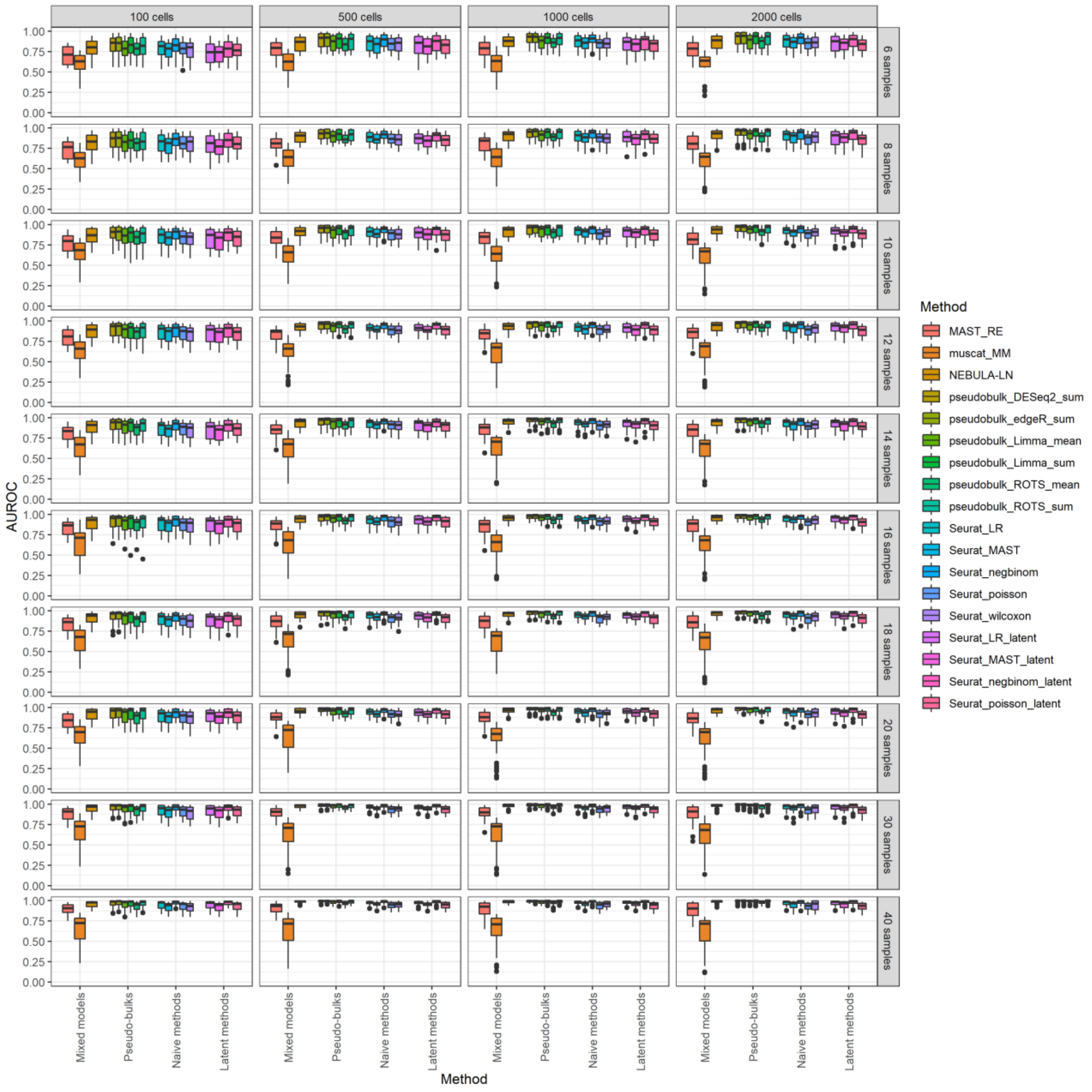


**Supplementary Figure 1. Area Under Receiver Operating Characteristic (AUROC) values for the reference-free negative binomial generative simulation**. Each boxplot includes results of 32 data sets simulated using four cell overdispersion values (0.1,1,10,100), four sample overdispersion values (0.05, 0.1, 0.2, 0.5) and two cell number distributions (imbalanced=negative binomial distribution, balanced=poisson distribution). The results are grouped by the average number of cells per subject (columns) and the number of subjects per dataset (e.g. 6 means 3 vs. 3). Each dataset includes 2000 genes, of which 100 are defined as positives by having $2 \geq \text{logFC} \geq 0.5$ between the two conditions, and the rest 1900 have logFC=0. The simulation is based on the simulation in the original paper of the NEBULA method. To estimate AUROC, we used the uncorrected p-values as the predicted values. We considered only those genes that were not filtered out by any of the methods.

### Supplementary Figure 2

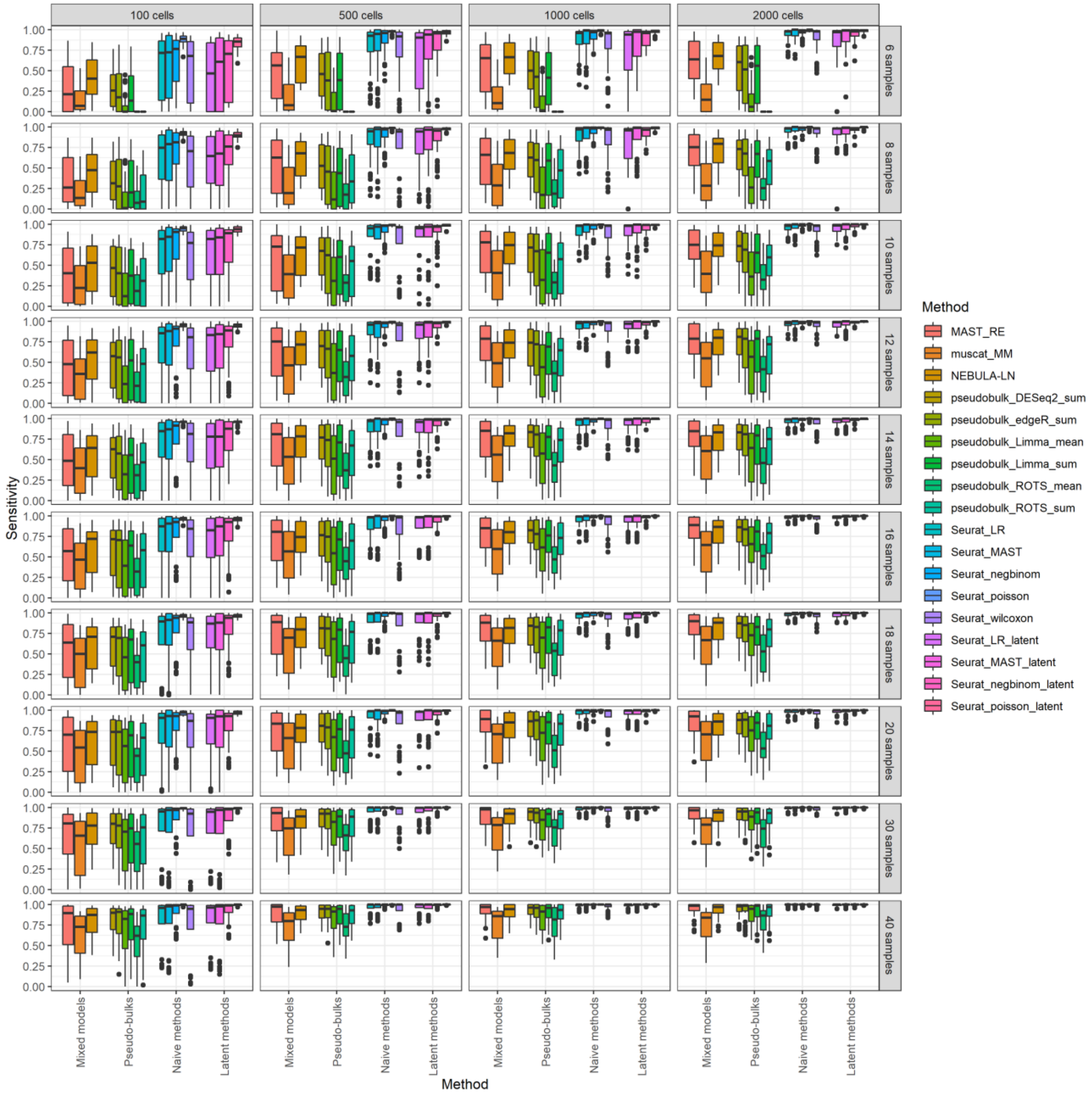


**Supplementary Figure 2. Sensitivity values for the reference-free negative binomial generative simulation**. Each boxplot includes results of 32 data sets simulated using four cell overdispersion values (0.1,1,10,100), four sample overdispersion values (0.05, 0.1, 0.2, 0.5) and two cell number distributions (imbalanced=negative binomial distribution, balanced=poisson distribution). The results are grouped by the average number of cells per subject (columns) and the number of subjects per dataset (e.g. 6 means 3 vs. 3). Each dataset includes 2000 genes, of which 100 are defined as positives by having 2 ≥ logFC ≥ 0.5 between the two conditions, and the rest 1900 have logFC=0. The simulation is based on the simulation in the original paper of the NEBULA method. To define the positive and negative DE genes for each method, we adjusted the p-values using the Benjamini-Hochberg method (also known as the FDR method) and used FDR=0.05 as the threshold. We considered only those genes that were not filtered out by any of the methods.

### Supplementary Figure 3

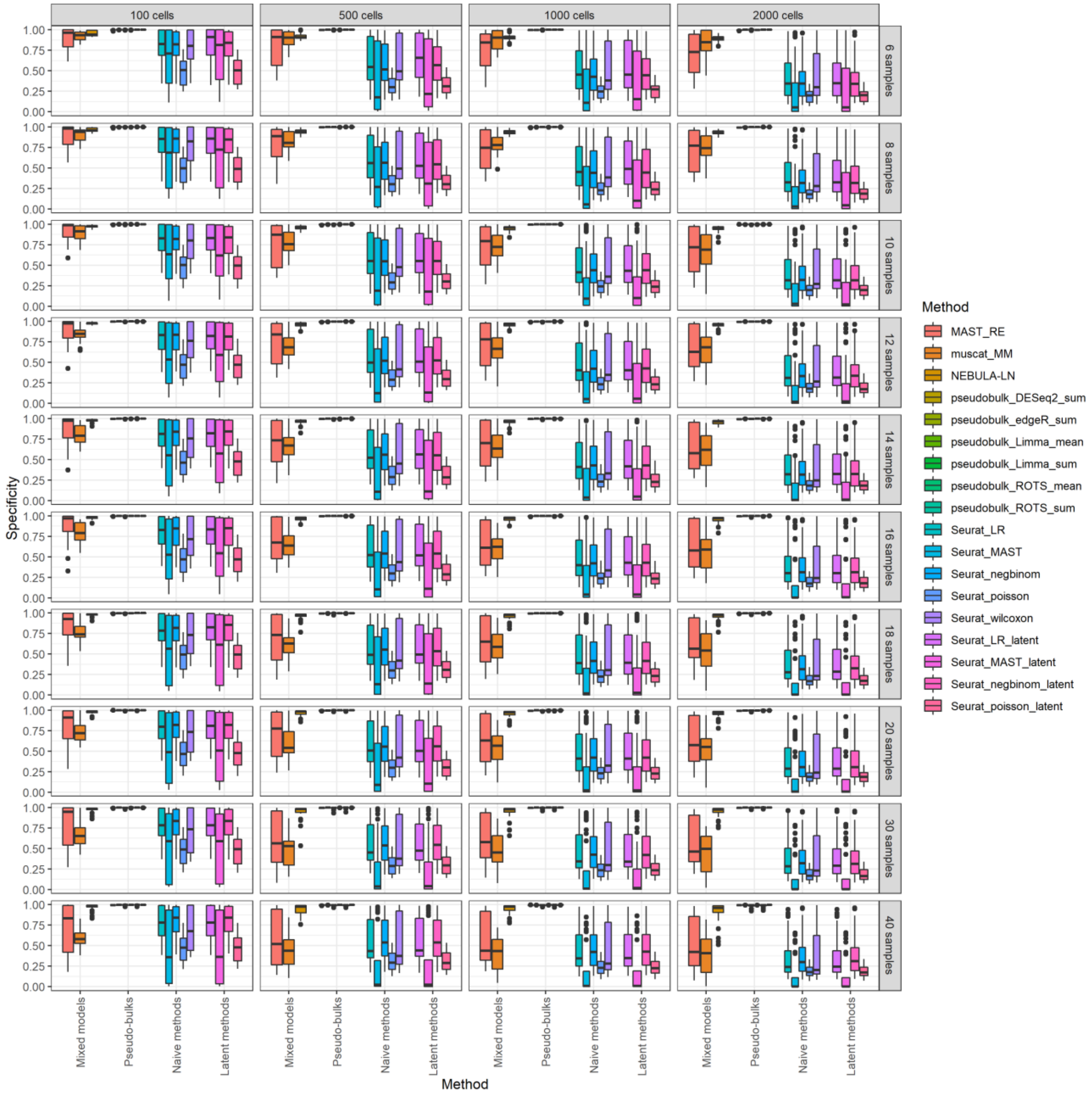


**Supplementary Figure 3. Specificity values for the reference-free negative binomial generative simulation**. Each boxplot includes results of 32 data sets simulated using four cell overdispersion values (0.1,1,10,100), four sample overdispersion values (0.05, 0.1, 0.2, 0.5) and two cell number distributions (imbalanced=negative binomial distribution, balanced=poisson distribution). The results are grouped by the average number of cells per subject (columns) and the number of subjects per dataset (e.g. 6 means 3 vs. 3). Each dataset includes 2000 genes, of which 100 are defined as positives by having $2 \geq \text{logFC} \geq 0.5$ between the two conditions, and the rest 1900 have logFC=0. The simulation is based on the simulation in the original paper of the NEBULA method. To define the positive and negative DE genes for each method, we adjusted the p-values using the Benjamini-Hochberg method (also known as the FDR method) and used FDR=0.05 as the threshold. We considered only those genes that were not filtered out by any of the methods.

### Supplementary Figure 4

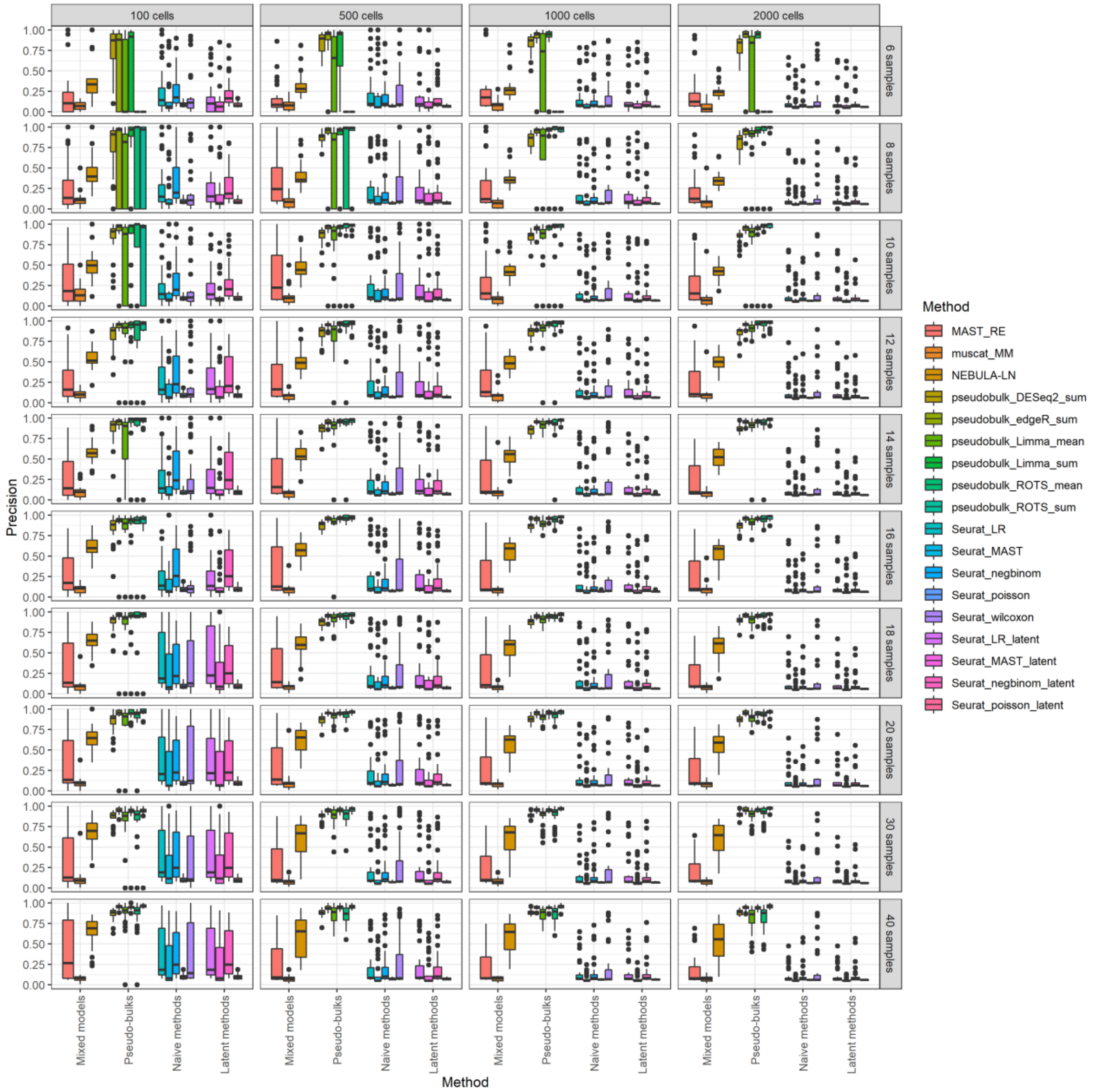


**Supplementary Figure 4. Precision values for the reference-free negative binomial generative simulation**. Each boxplot includes results of 32 data sets simulated using four cell overdispersion values (0.1,1,10,100), four sample overdispersion values (0.05, 0.1, 0.2, 0.5) and two cell number distributions (imbalanced=negative binomial distribution, balanced=poisson distribution). The results are grouped by the average number of cells per subject (columns) and the number of subjects per dataset (e.g. 6 means 3 vs. 3). Each dataset includes 2000 genes, of which 100 are defined as positives by having 2 ≥ logFC ≥ 0.5 between the two conditions, and the rest 1900 have logFC=0. The simulation is based on the simulation in the original paper of the NEBULA method. To define the positive and negative DE genes for each method, we adjusted the p-values using the Benjamini-Hochberg method (also known as the FDR method) and used FDR=0.05 as the threshold. We considered only those genes that were not filtered out by any of the methods. If the precision could not be calculated because there were no positive genes in the results, we set the precision to zero.

### Supplementary Figure 5

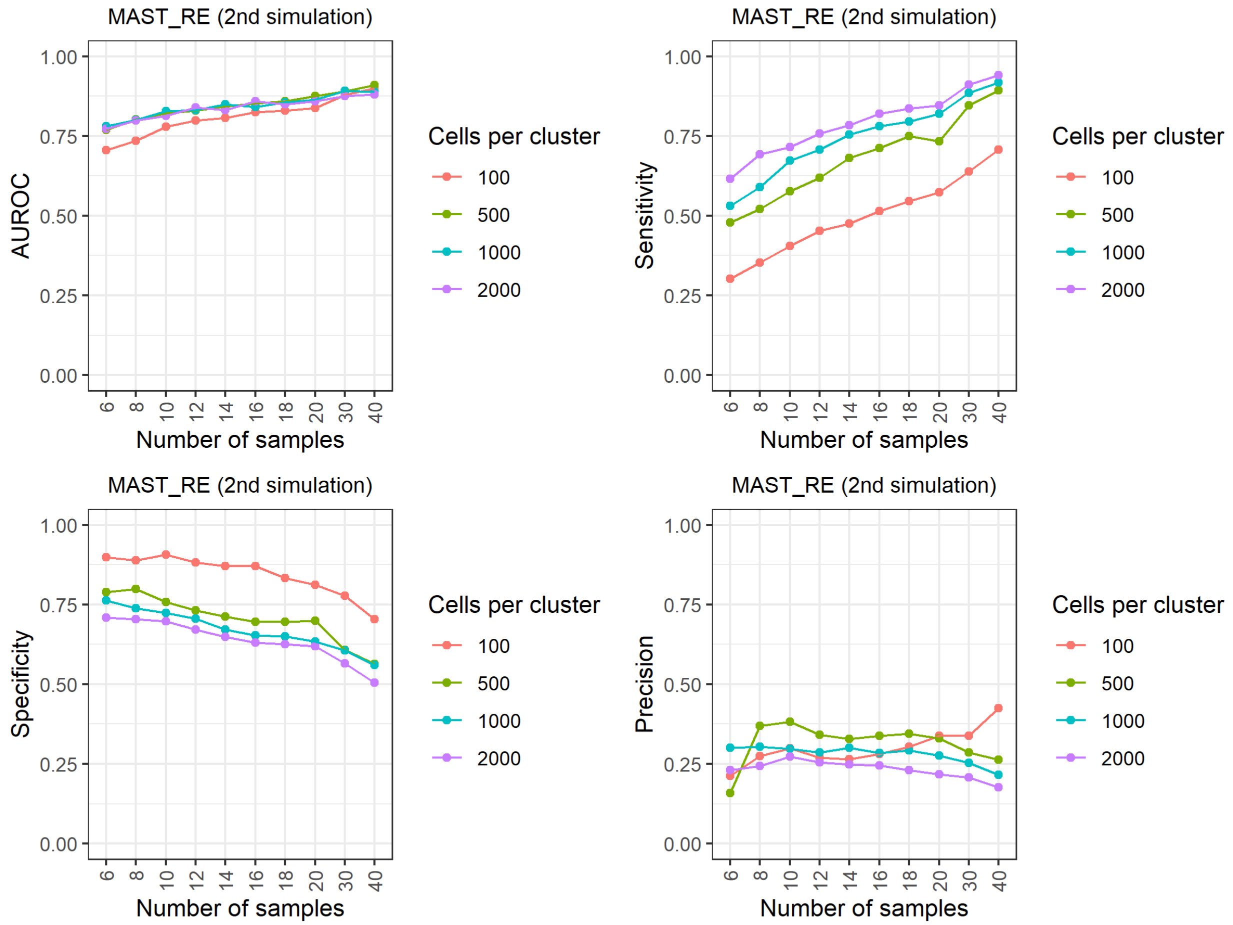


**Supplementary Figure 5. Results of the reference-free negative binomial generative simulation for MAST_RE**.

### Supplementary Figure 6

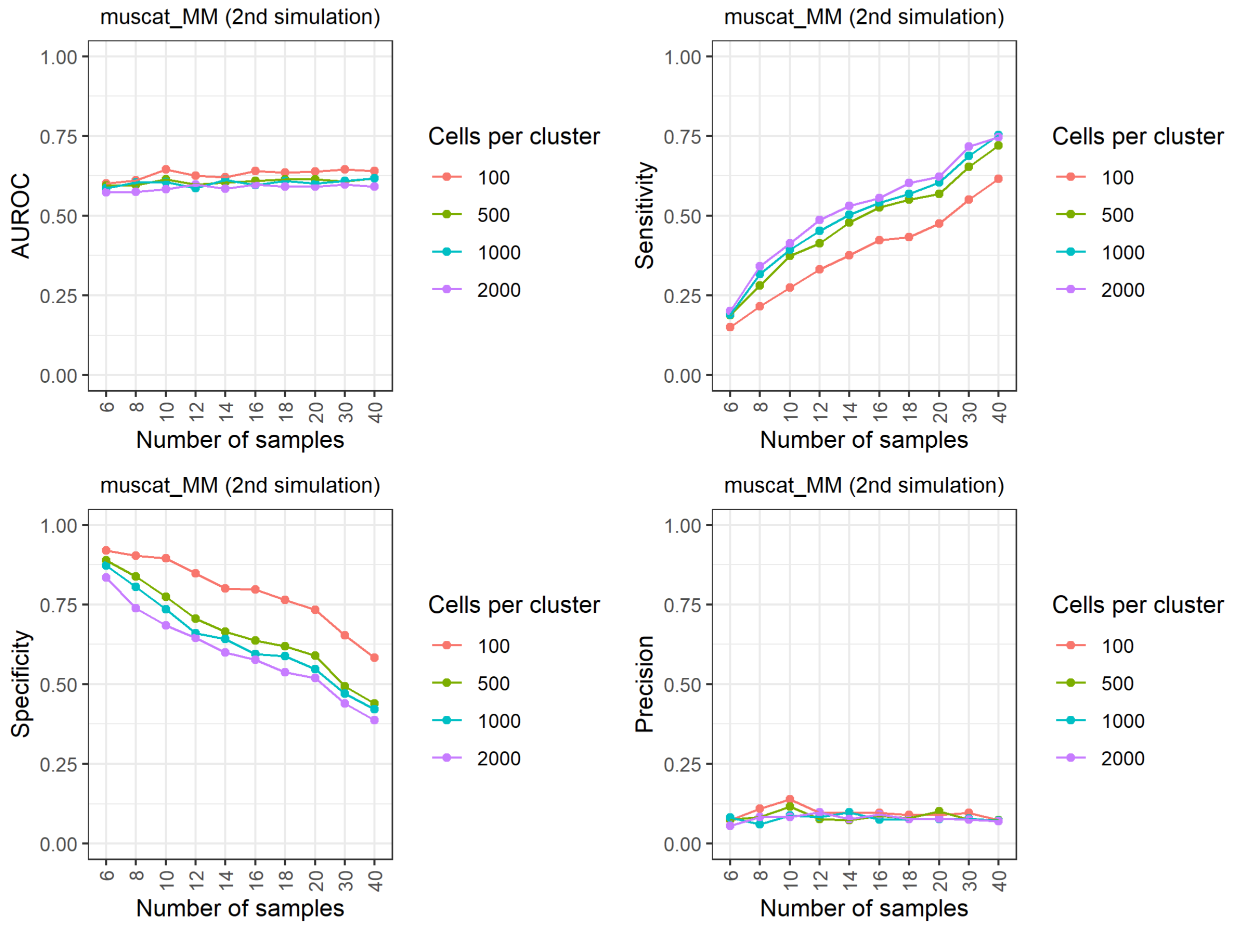


**Supplementary Figure 6. Results of the reference-free negative binomial generative simulation for muscat_MM**.

### Supplementary Figure 7

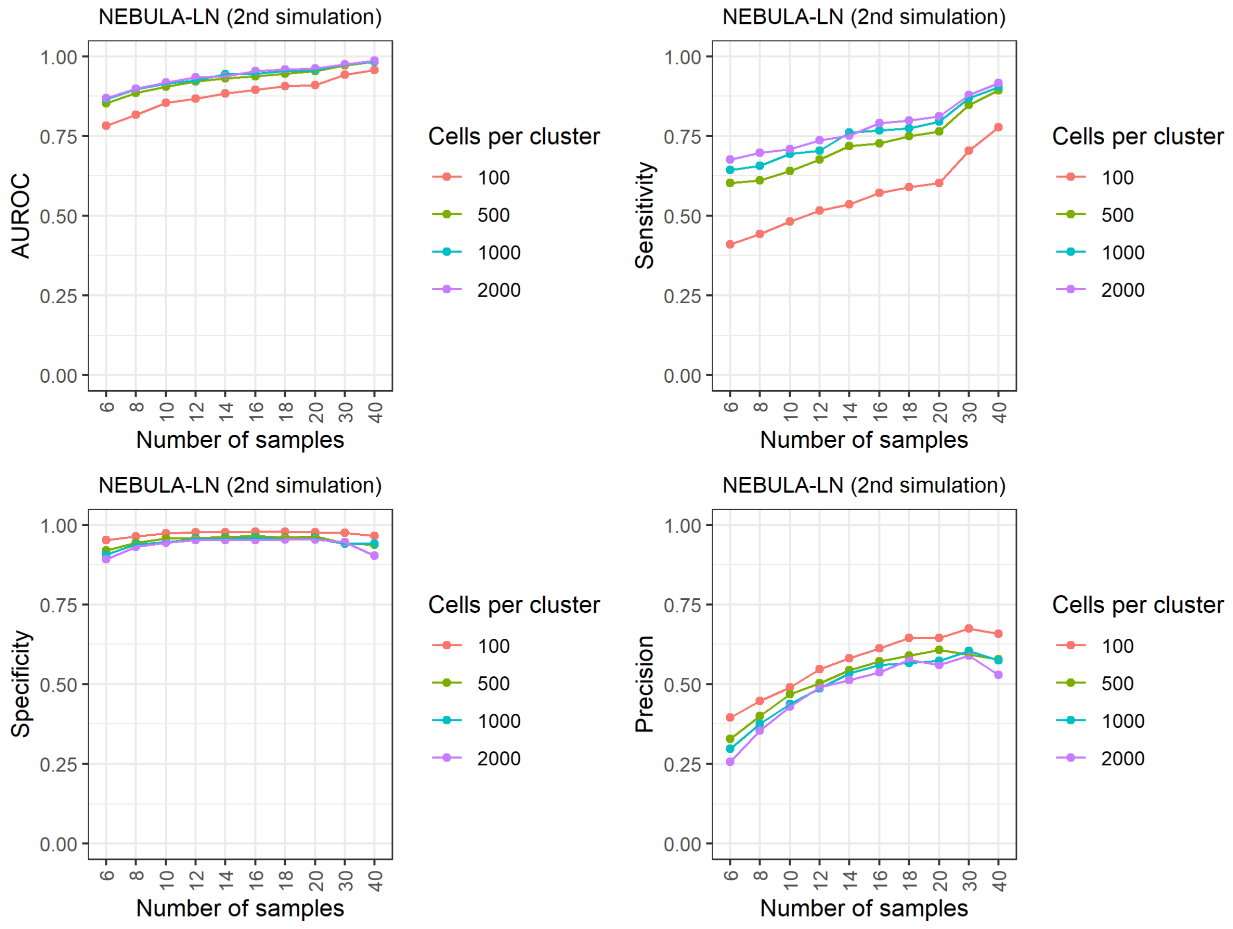


**Supplementary Figure 7. Results of the reference-free negative binomial generative simulation for NEBULA-LN**.

### Supplementary Figure 8

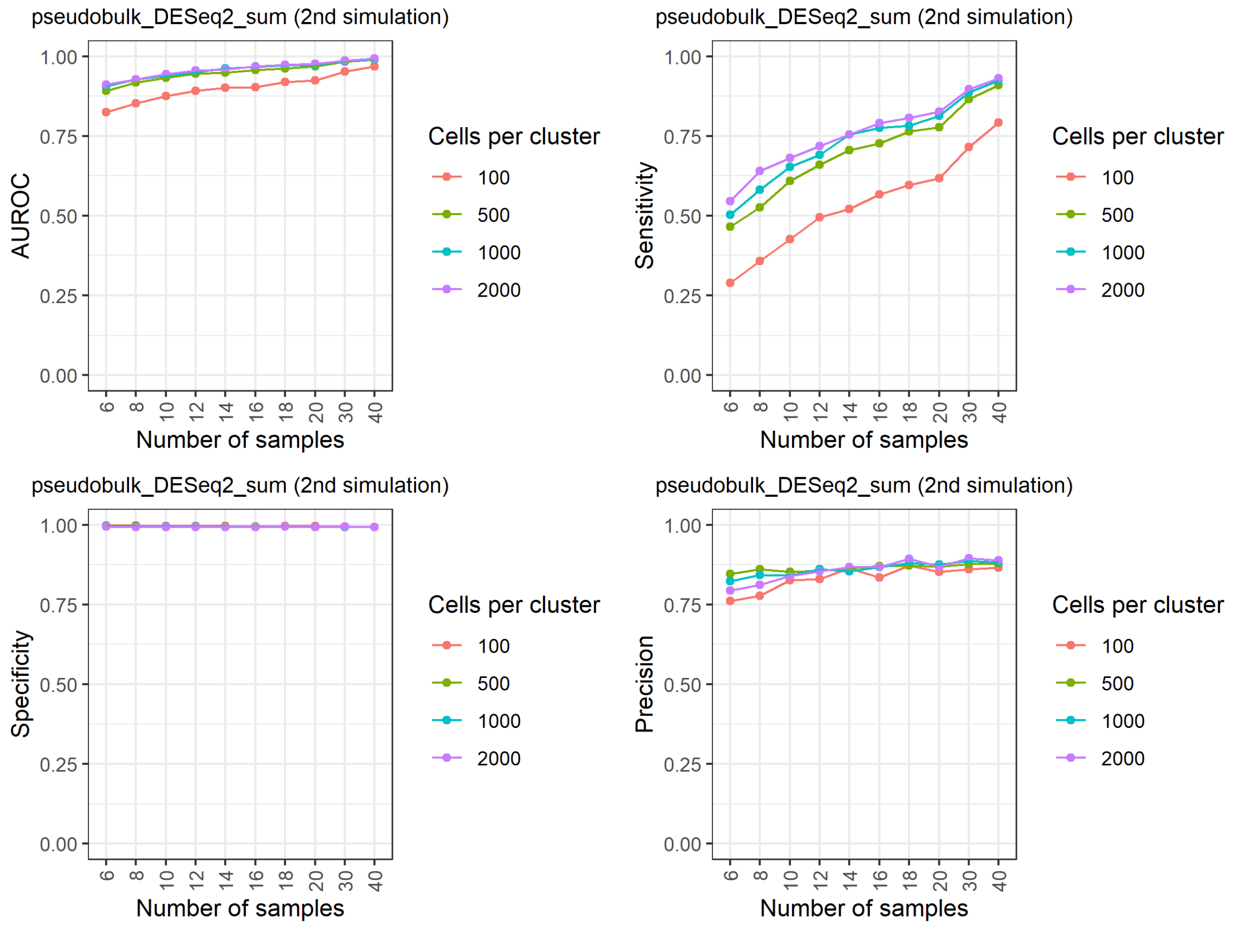


**Supplementary Figure 8. Results of the reference-free negative binomial generative simulation for pseudobulk_DESeq2_sum**.

### Supplementary Figure 9

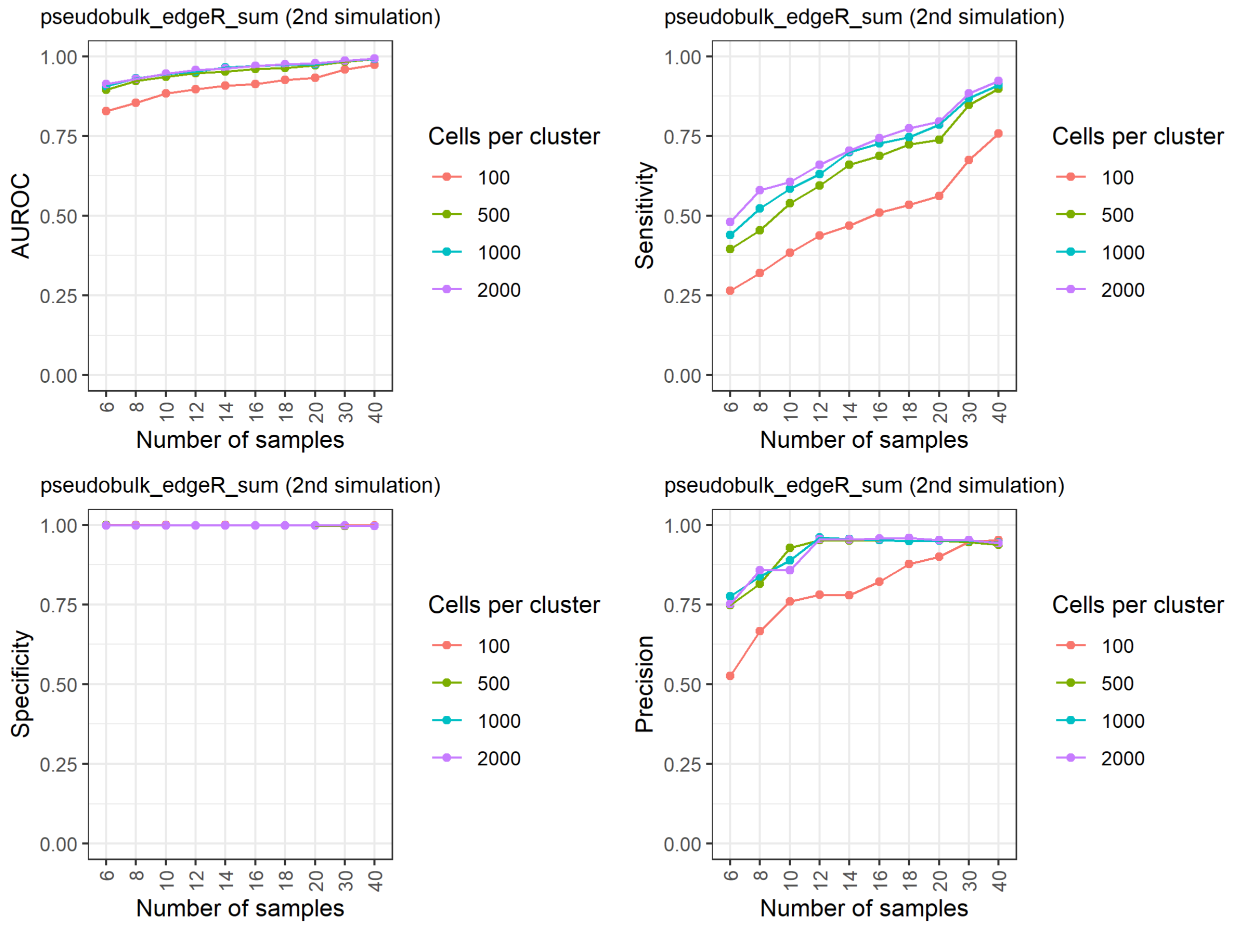


**Supplementary Figure 9. Results of the reference-free negative binomial generative simulation for pseudobulk_edgeR_sum**.

### Supplementary Figure 10

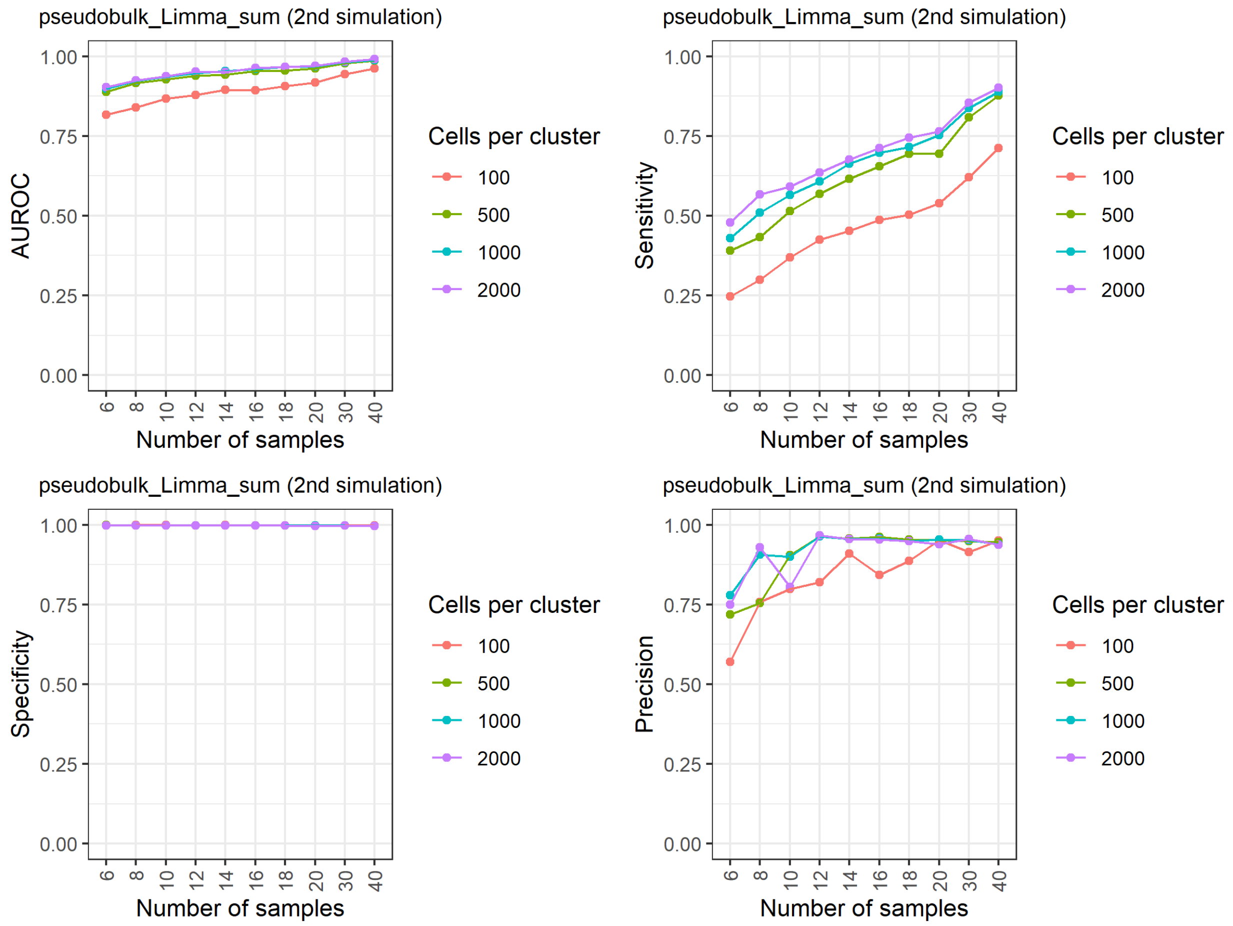


**Supplementary Figure 10. Results of the reference-free negative binomial generative simulation for pseudobulk_Limma_sum**.

### Supplementary Figure 11

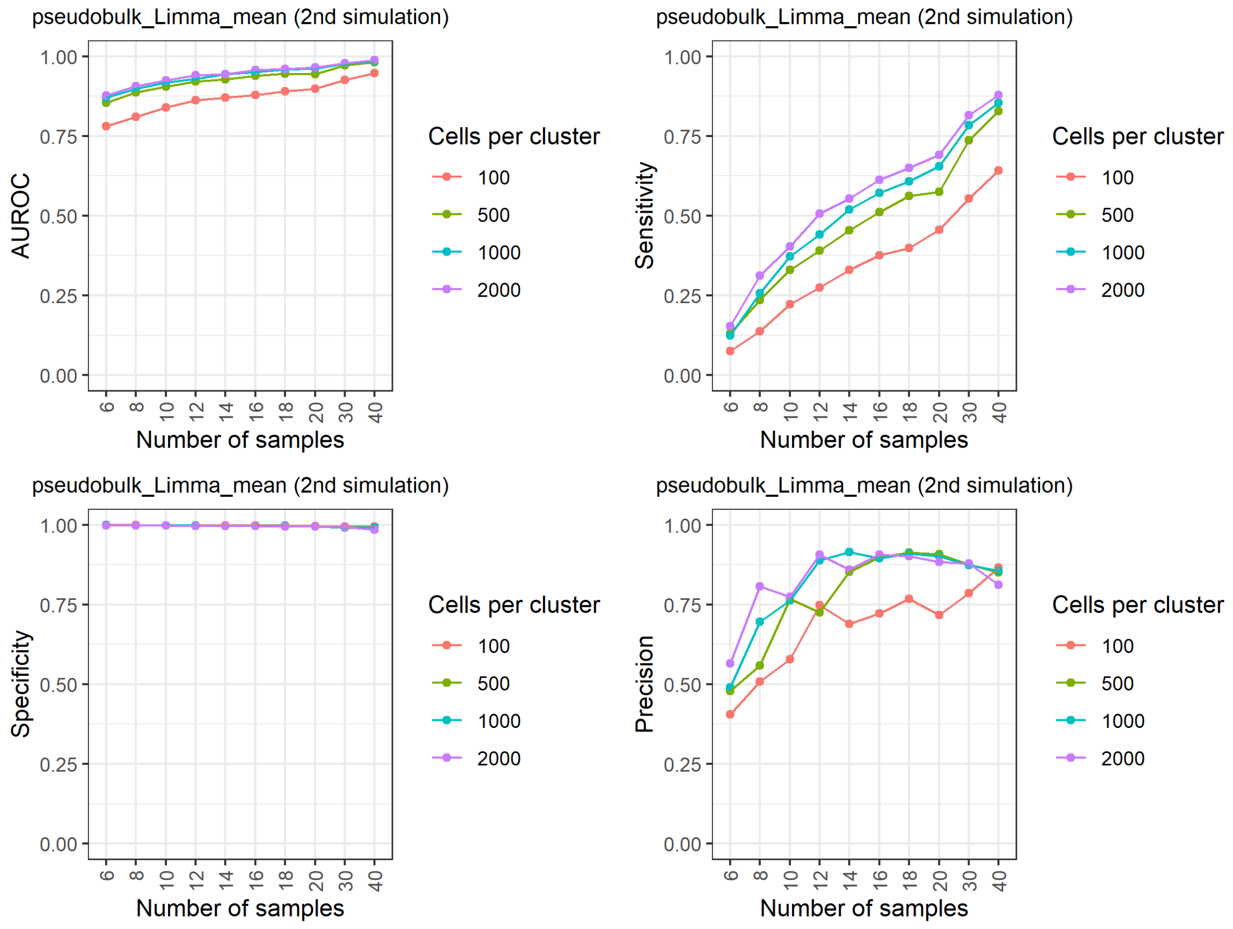


**Supplementary Figure 11. Results of the reference-free negative binomial generative simulation for pseudobulk_Limma_mean**.

### Supplementary Figure 12

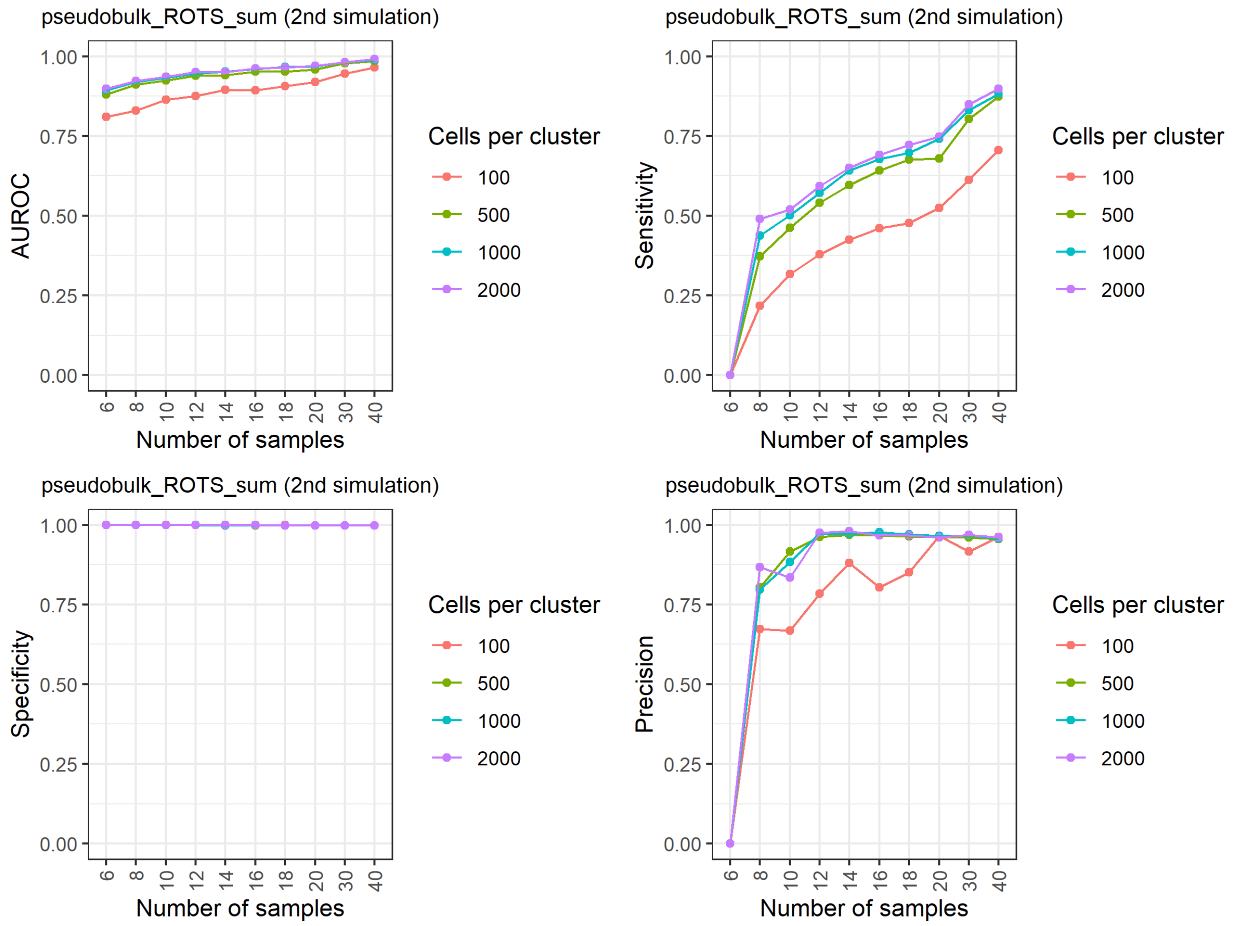


**Supplementary Figure 12. Results of the reference-free negative binomial generative simulation for pseudobulk_ROTS_sum**.

### Supplementary Figure 13

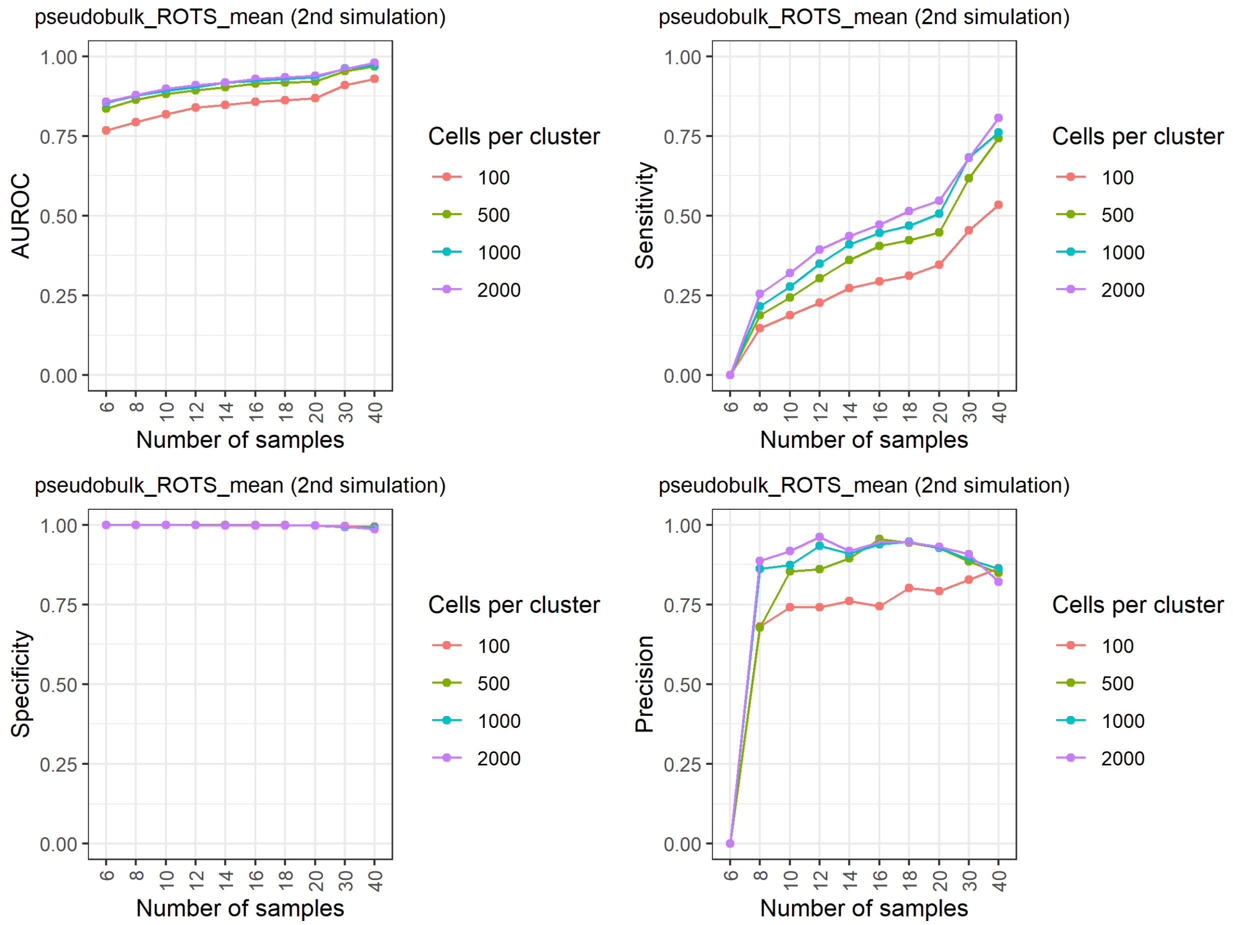


**Supplementary Figure 13. Results of the reference-free negative binomial generative simulation for pseudobulk_ROTS_mean**.

### Supplementary Figure 14

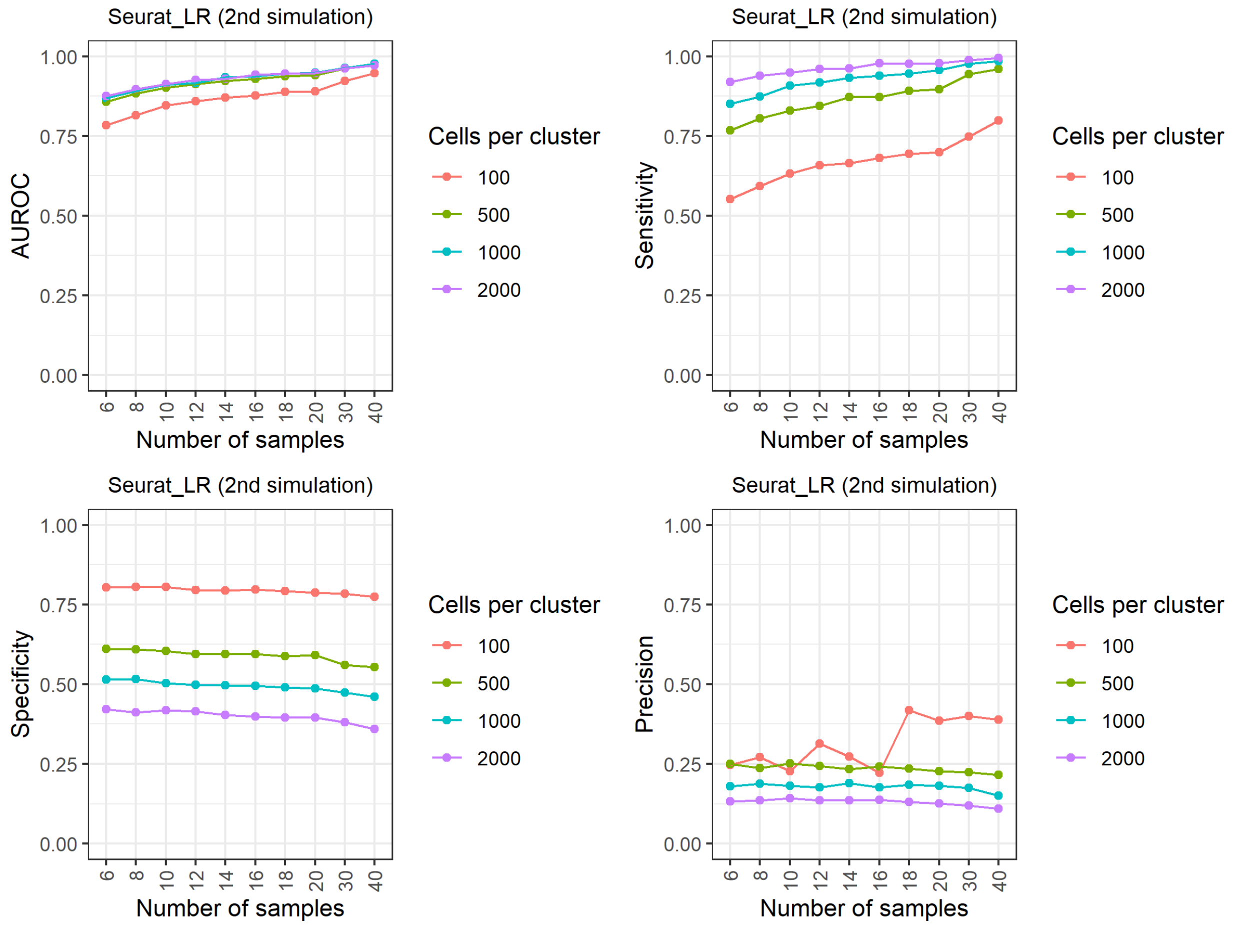


**Supplementary Figure 14. Results of the reference-free negative binomial generative simulation for Seurat_LR**.

### Supplementary Figure 15

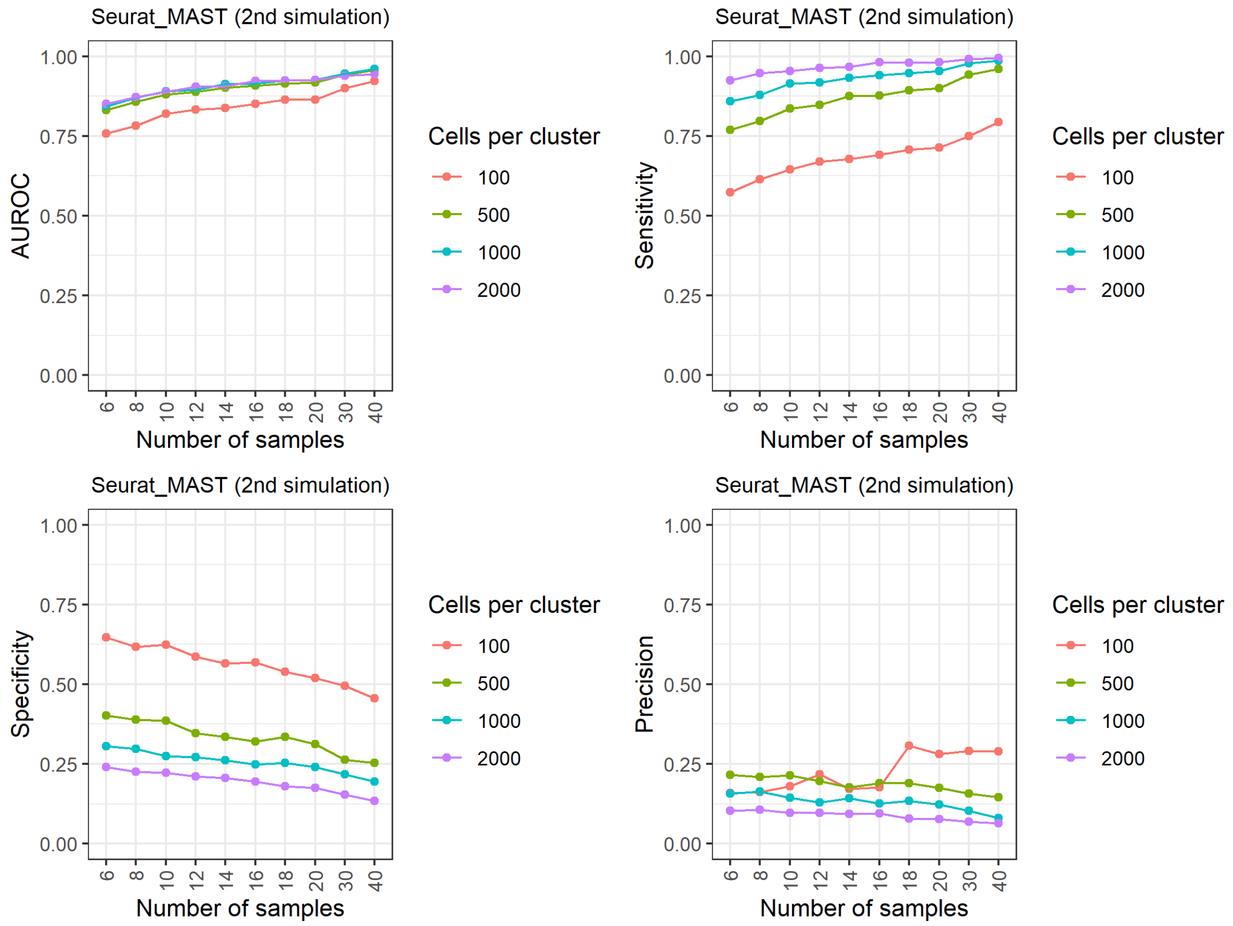


**Supplementary Figure 15. Results of the reference-free negative binomial generative simulation for Seurat_MAST**.

### Supplementary Figure 16

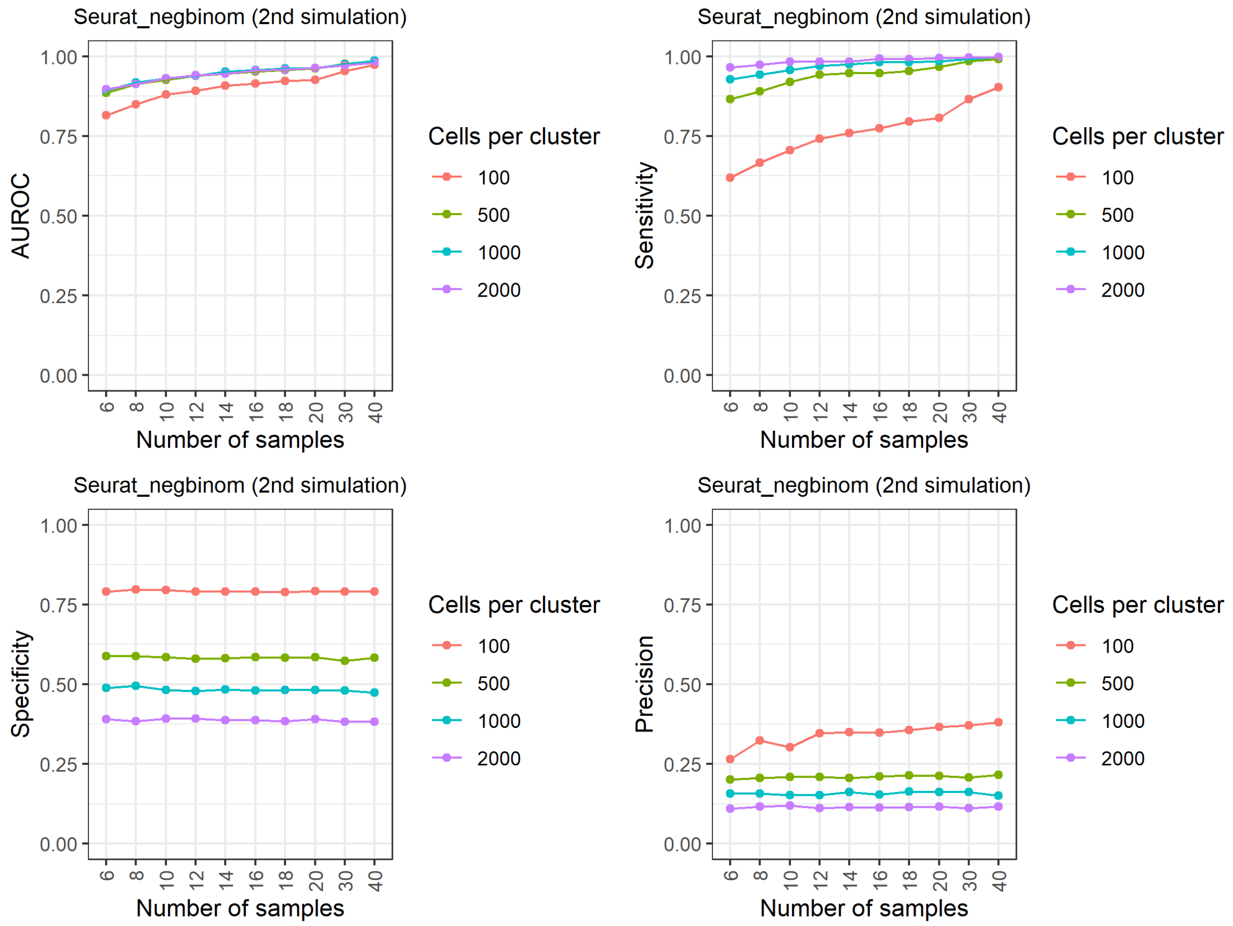


**Supplementary Figure 16. Results of the reference-free negative binomial generative simulation for Seurat_negbinom**.

### Supplementary Figure 17

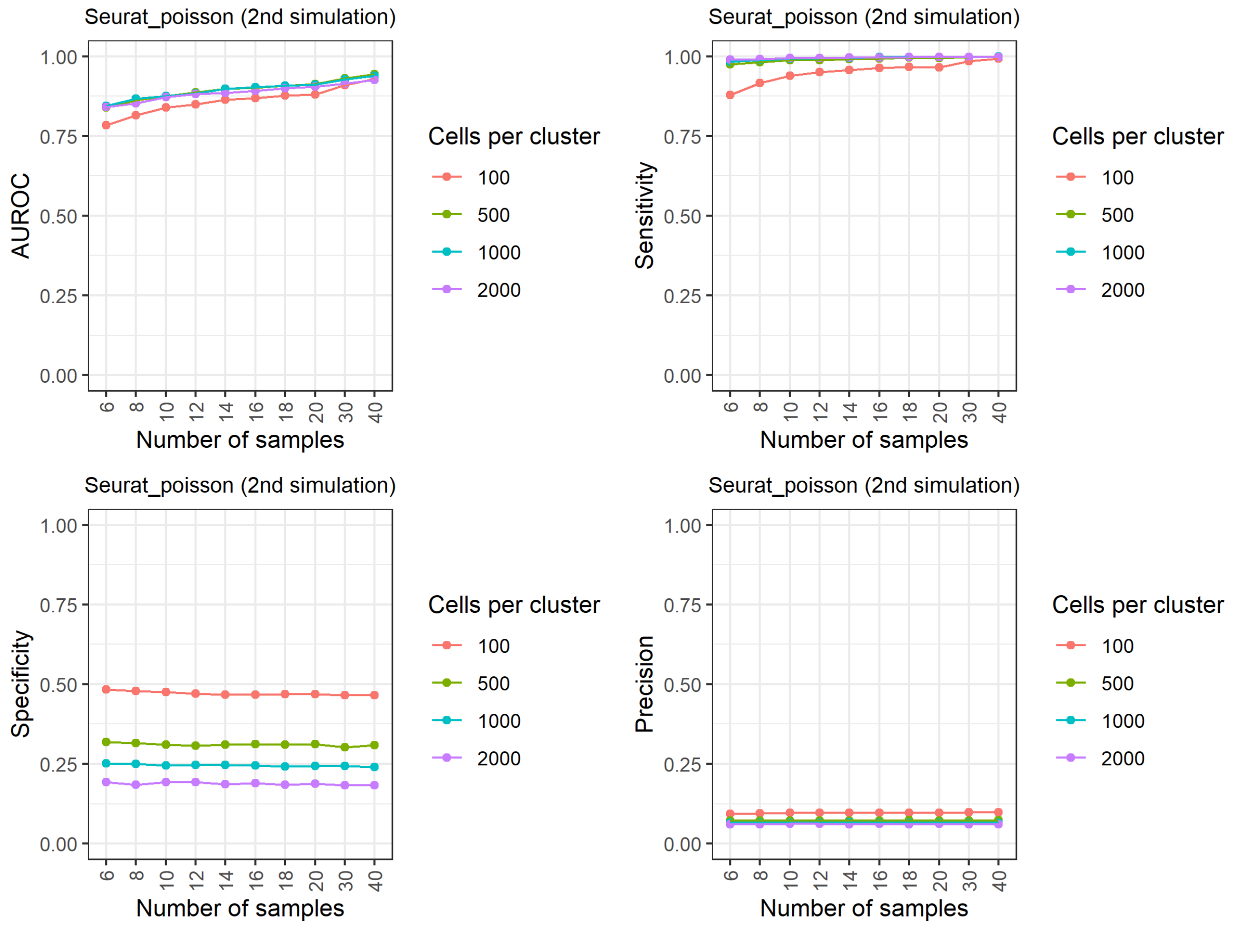


**Supplementary Figure 17. Results of the reference-free negative binomial generative simulation for Seurat_poisson**.

### Supplementary Figure 18

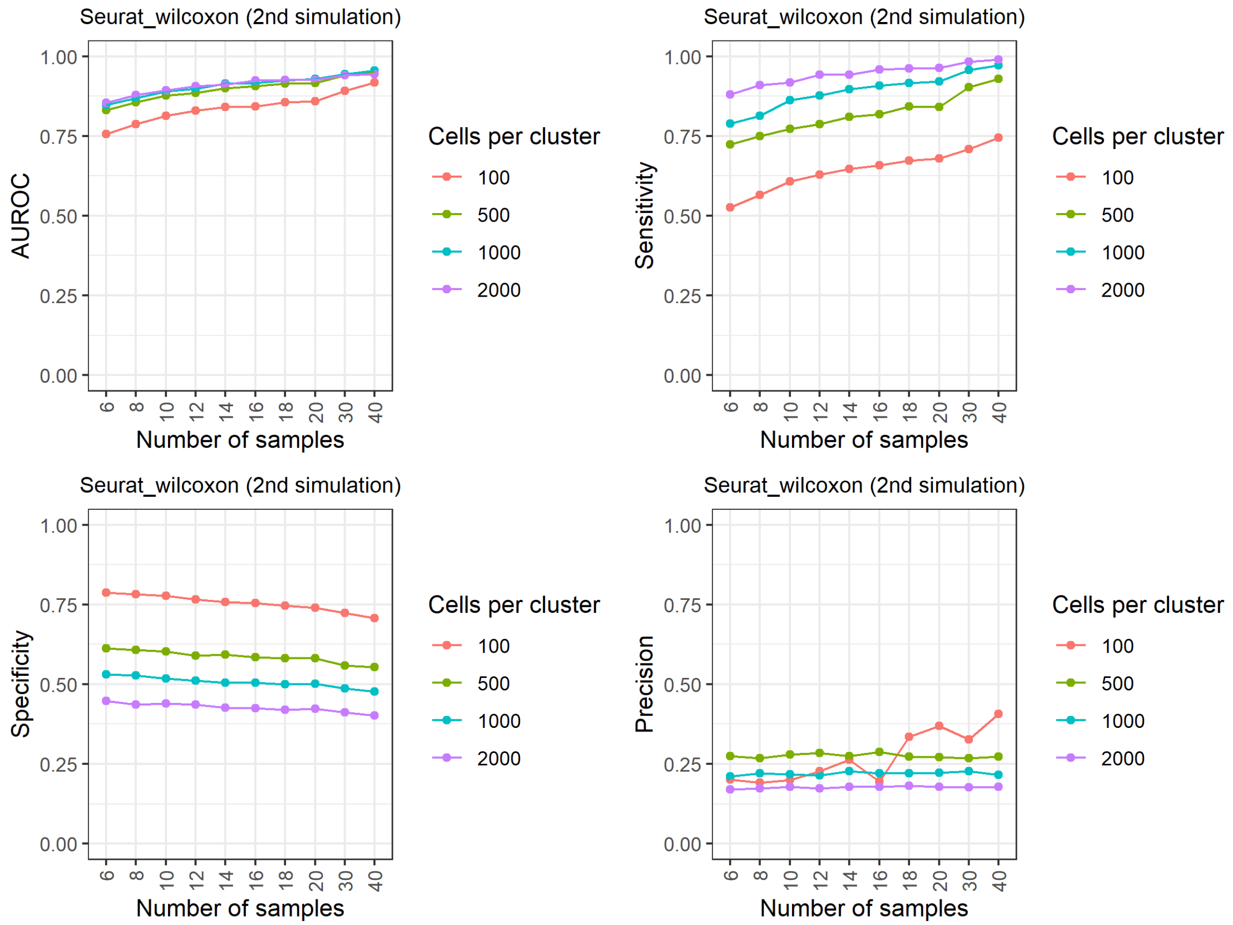


**Supplementary Figure 18. Results of the reference-free negative binomial generative simulation for Seurat_wilcoxon**.

### Supplementary Figure 19

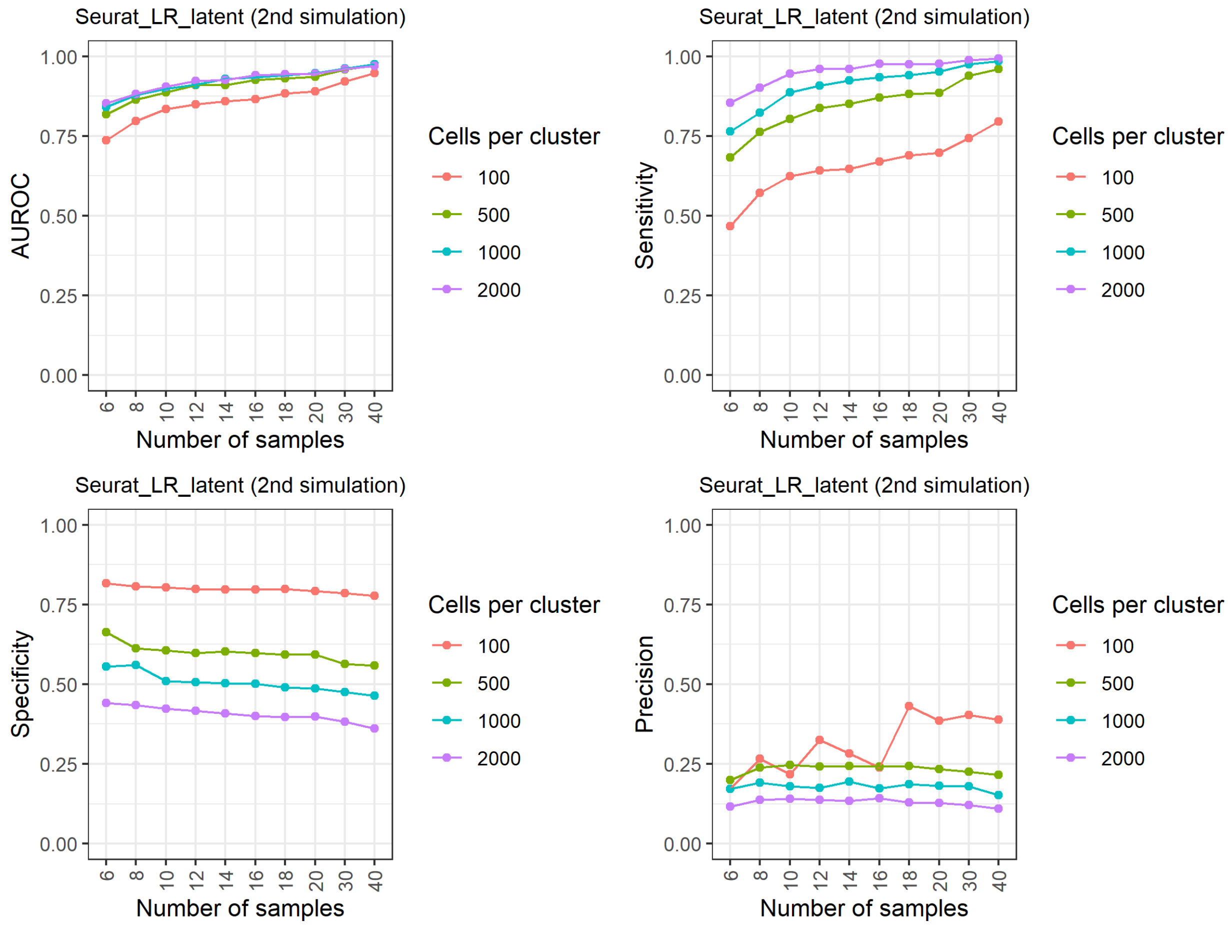


**Supplementary Figure 19. Results of the reference-free negative binomial generative simulation for Seurat_LR_latent**.

### Supplementary Figure 20

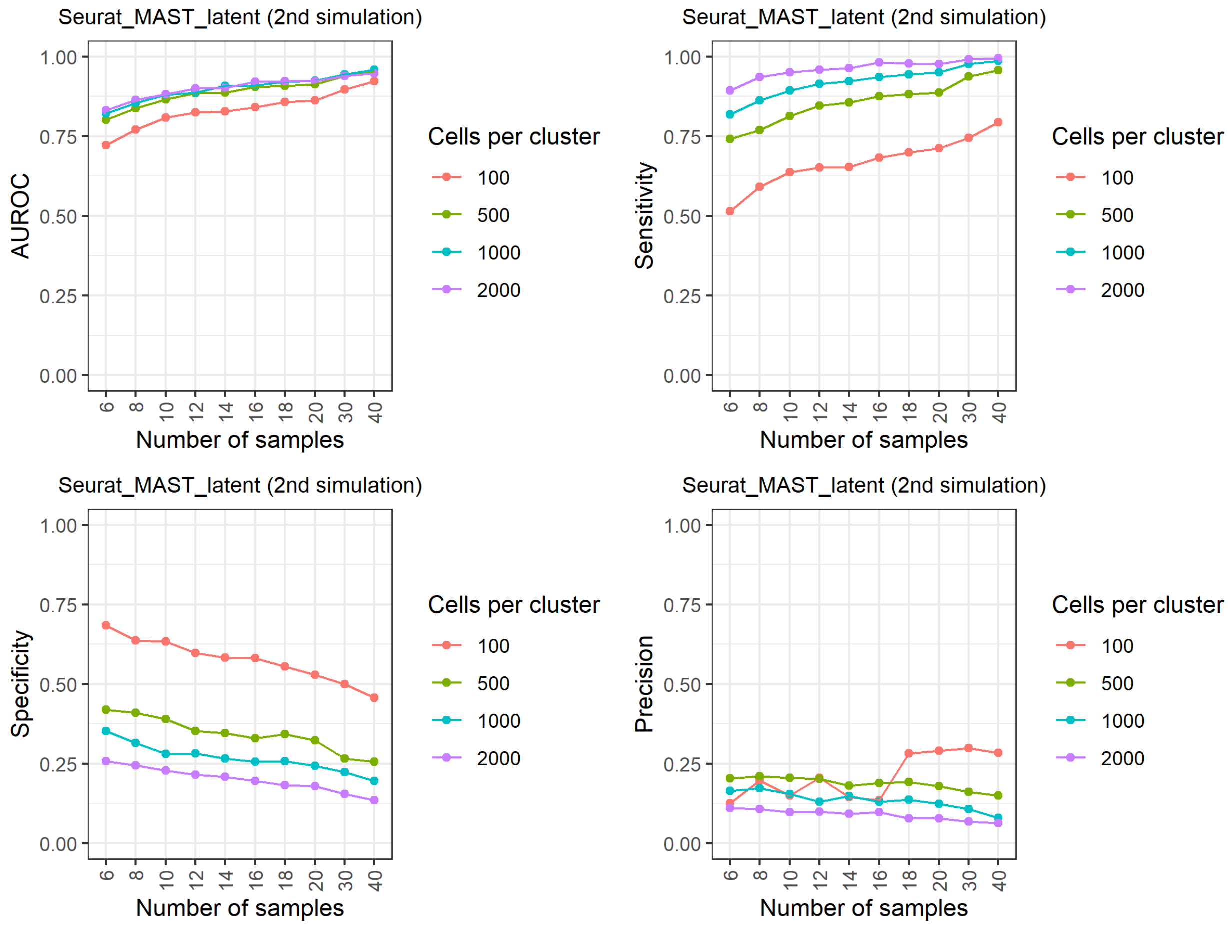


**Supplementary Figure 20. Results of the reference-free negative binomial generative simulation for Seurat_MAST_latent**.

### Supplementary Figure 21

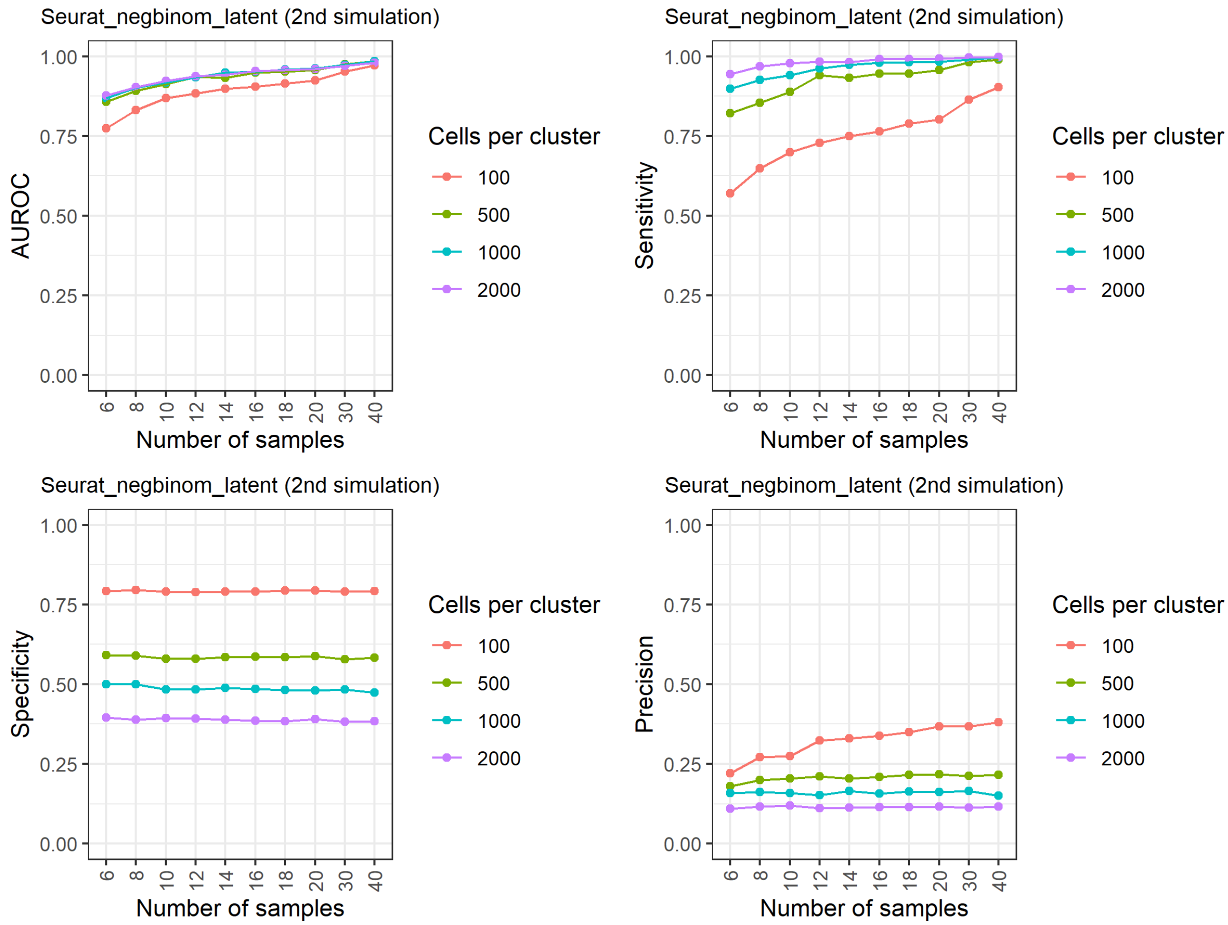


**Supplementary Figure 21. Results of the reference-free negative binomial generative simulation for Seurat_negbinom_latent**.

### Supplementary Figure 22

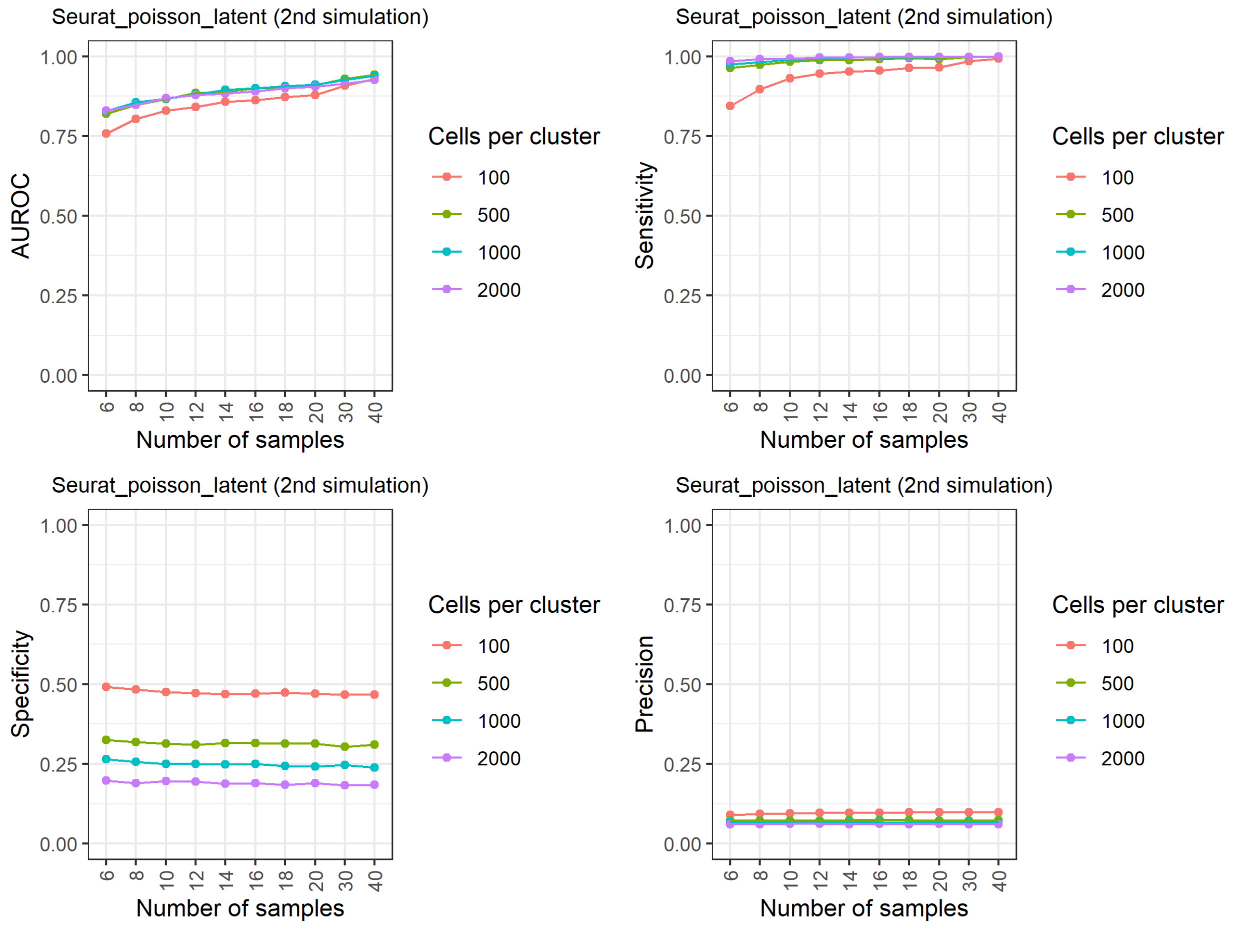


**Supplementary Figure 22. Results of the reference-free negative binomial generative simulation for Seurat_poisson_latent**.

### Supplementary Figure 23

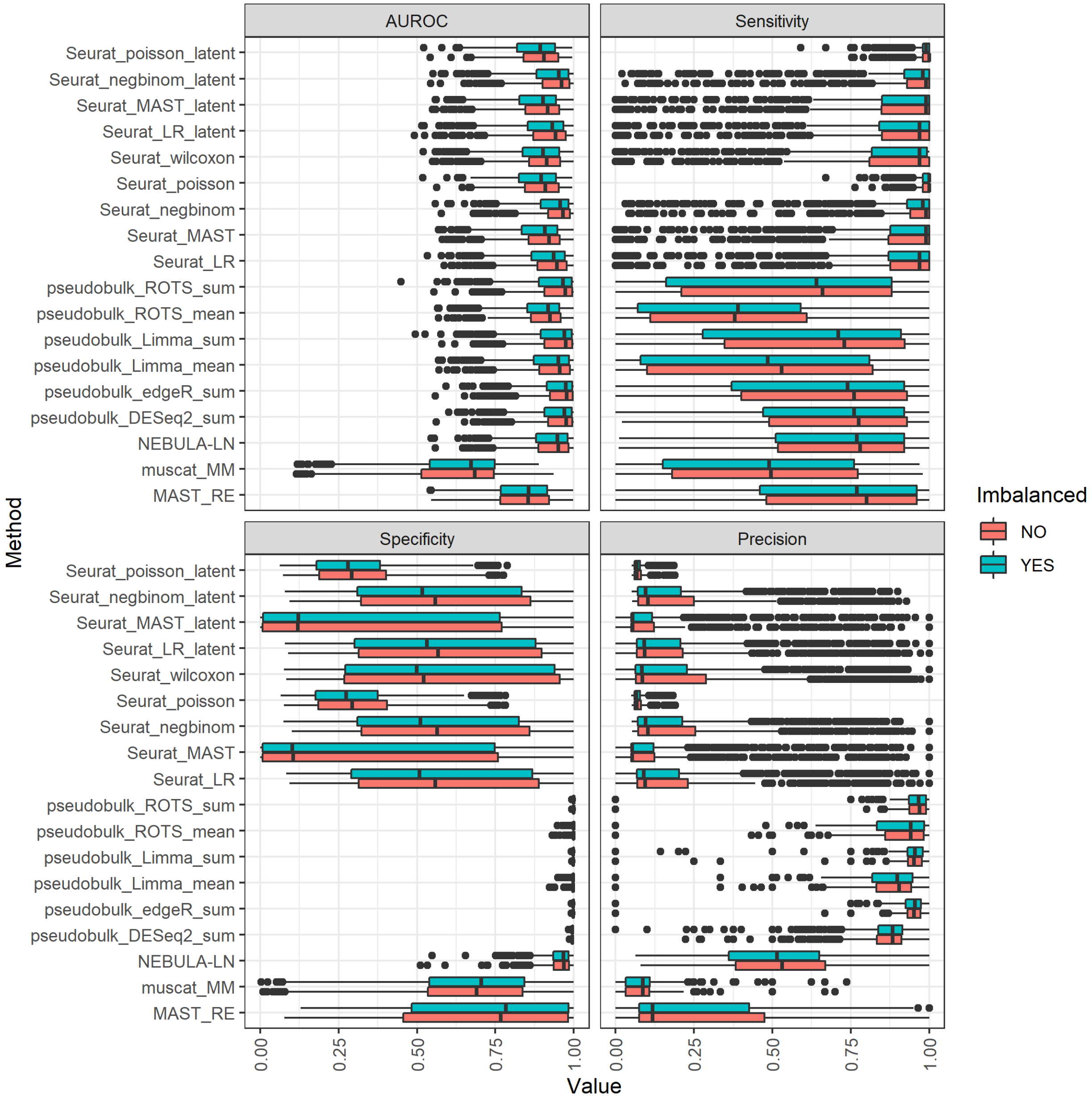


**Supplementary Figure 23. Results of the reference-free negative binomial generative simulation grouped by whether the data sets were downsampled to generate an imbalance distribution of cells across the samples or not.**

### Supplementary Figure 24

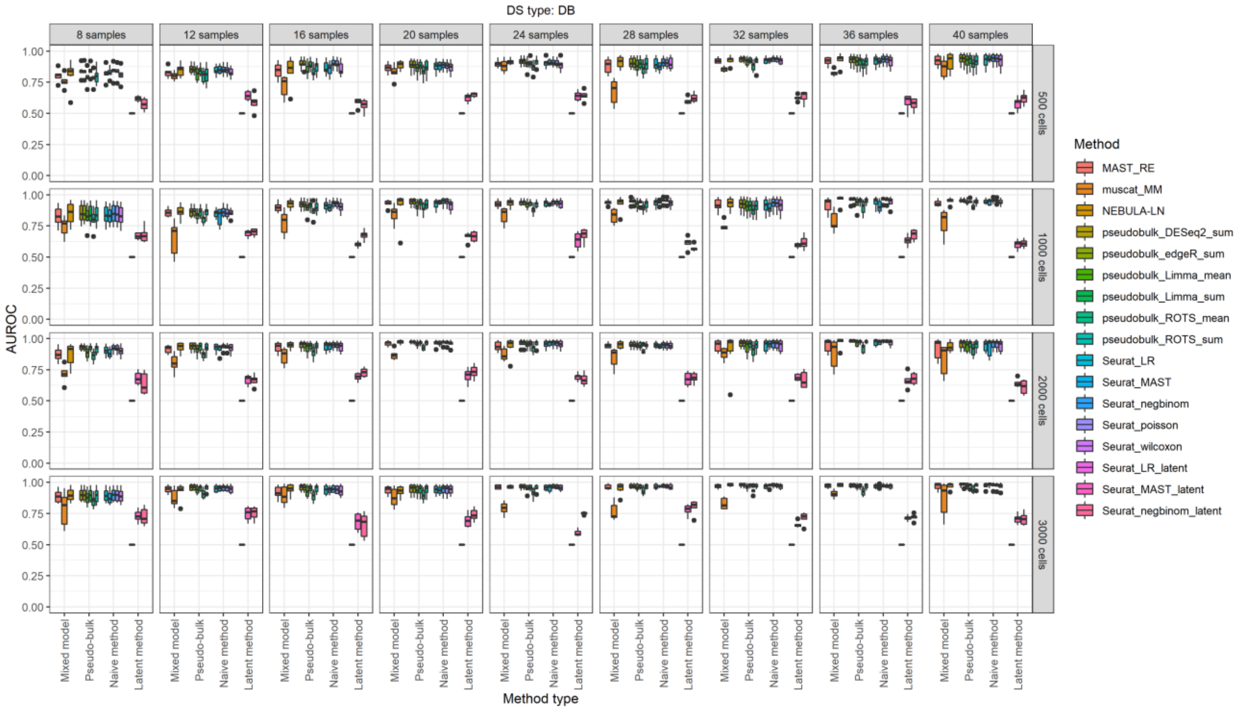


**Supplementary Figure 24. Area Under Receiver Operating Characteristic (AUROC) values for the reference-based negative binomial generative simulation (muscat).** The results are grouped in columns by the number of samples in the comparison and in the rows by the number of cells per subject. In this simulation, each sample includes three clusters. These results are for the differential state (DS) type, which includes changes in both modality and proportions (DB). See more details in **Section 2.2.1** of the manuscript.

### Supplementary Figure 25

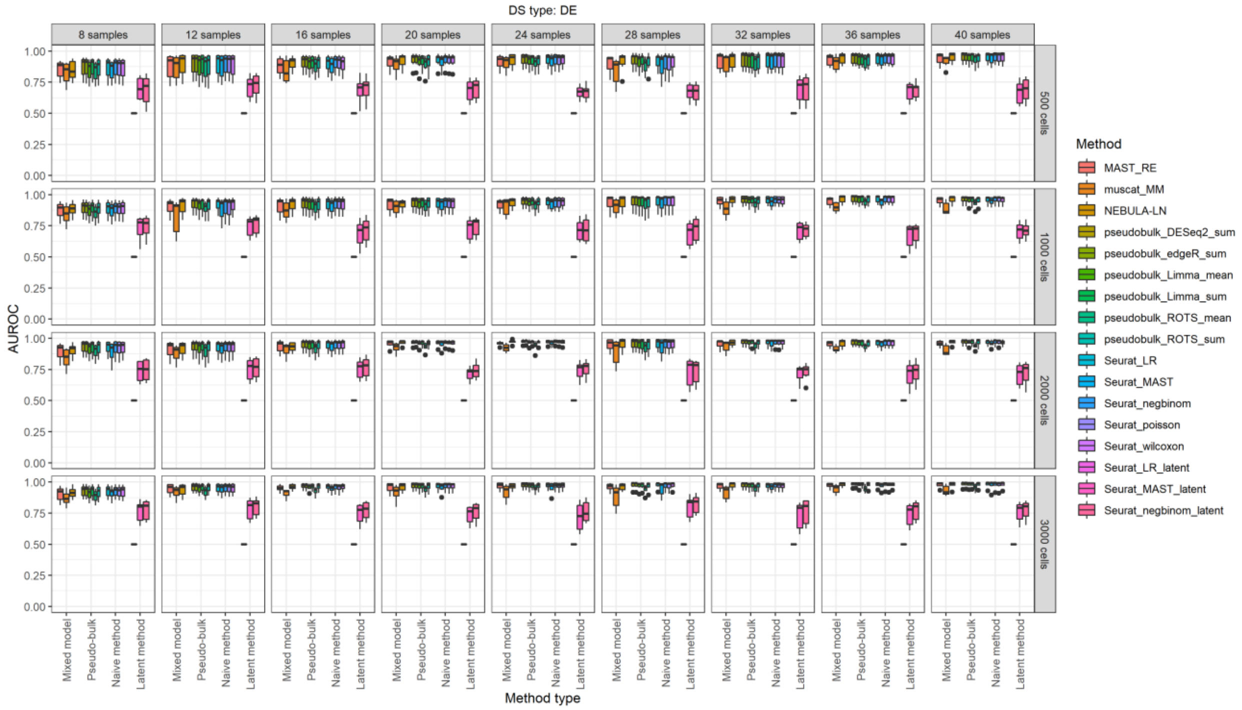


**Supplementary Figure 25. Area Under Receiver Operating Characteristic (AUROC) values for the reference-based negative binomial generative simulation (muscat).** The results are grouped in columns by the number of samples in the comparison and in the rows by the number of cells per subject. In this simulation, each sample includes three clusters. These results are for the differential state (DS) type, which includes changes in mean expression (DE). See more details on how the data was simulated in **Section 2.2.1** of the manuscript.

### Supplementary Figure 26

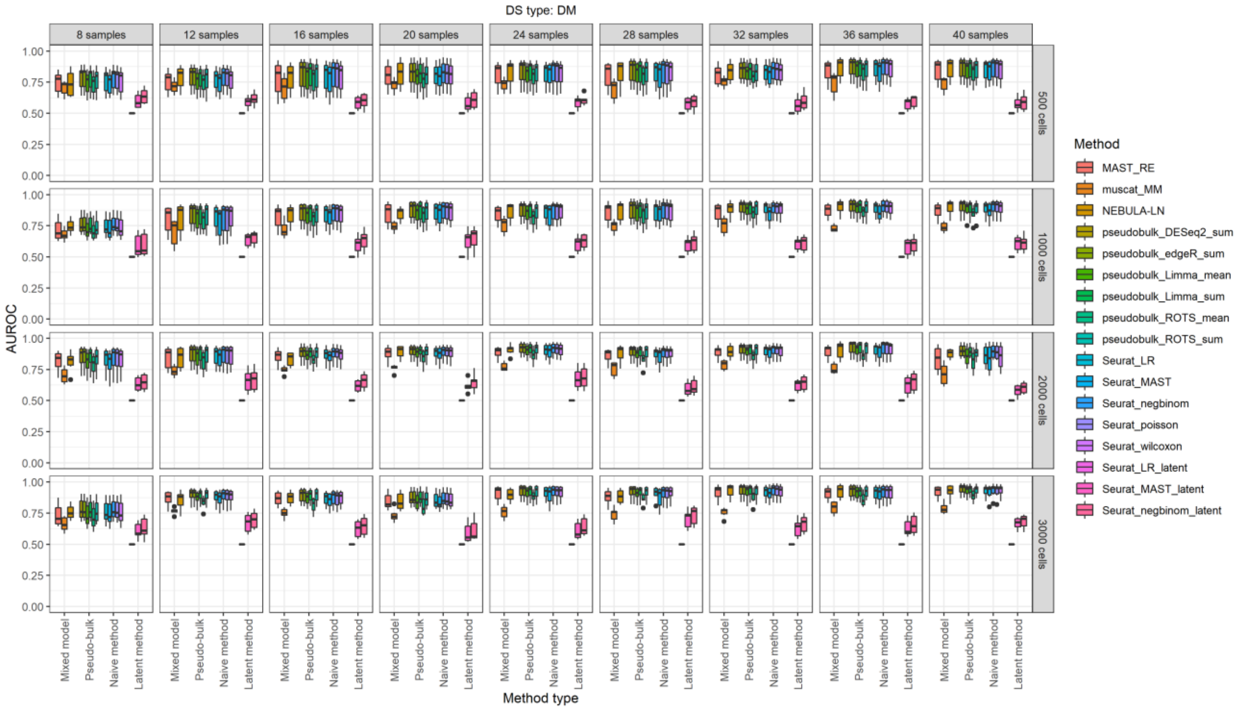


**Supplementary Figure 26. Area Under Receiver Operating Characteristic (AUROC) values for the reference-based negative binomial generative simulation (muscat).** The results are grouped in columns by the number of samples in the comparison and in the rows by the number of cells per subject. In this simulation, each sample includes three clusters. These results are for the differential state (DS) type, which includes changes in modality (DM).

### Supplementary Figure 27

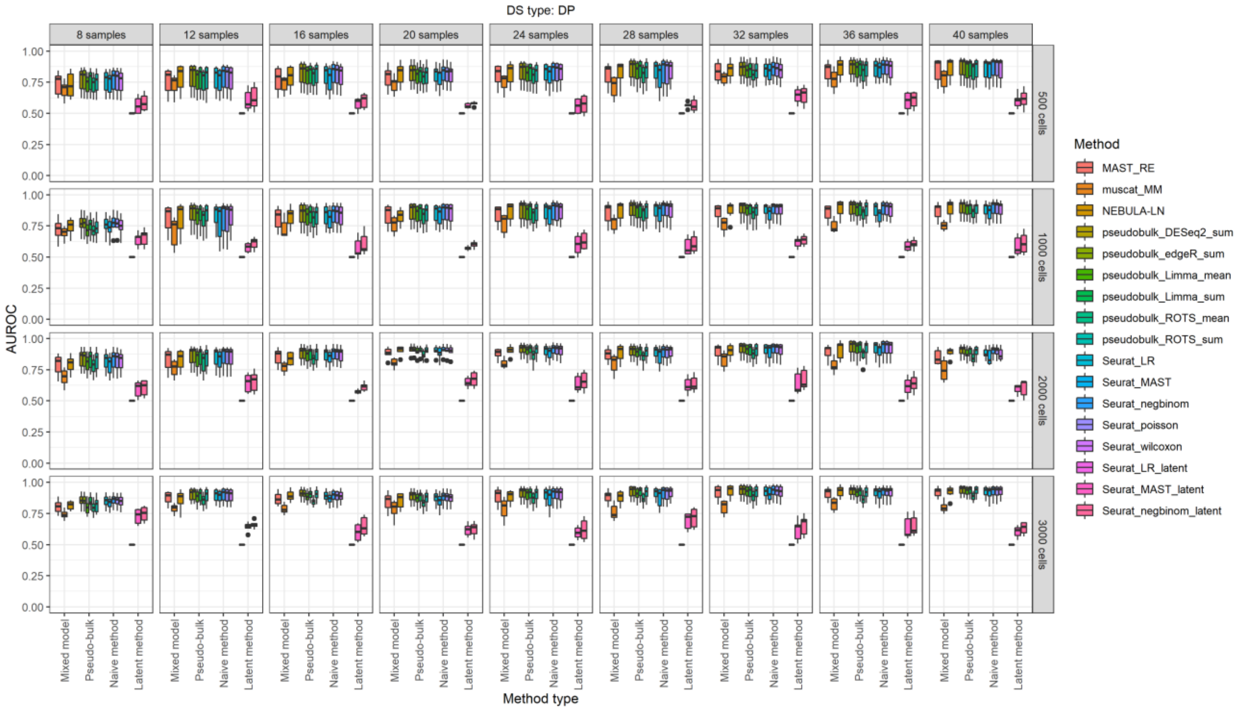


**Supplementary Figure 27 Area Under Receiver Operating Characteristic (AUROC) values for the reference-based negative binomial generative simulation (muscat).** The results are grouped in columns by the number of samples in the comparison and in the rows by the number of cells per subject. In this simulation, each sample includes three clusters. These results are for the differential state (DS) type, which includes changes in the proportions of low and high expression-state components (DP).

### Supplementary Figure 28

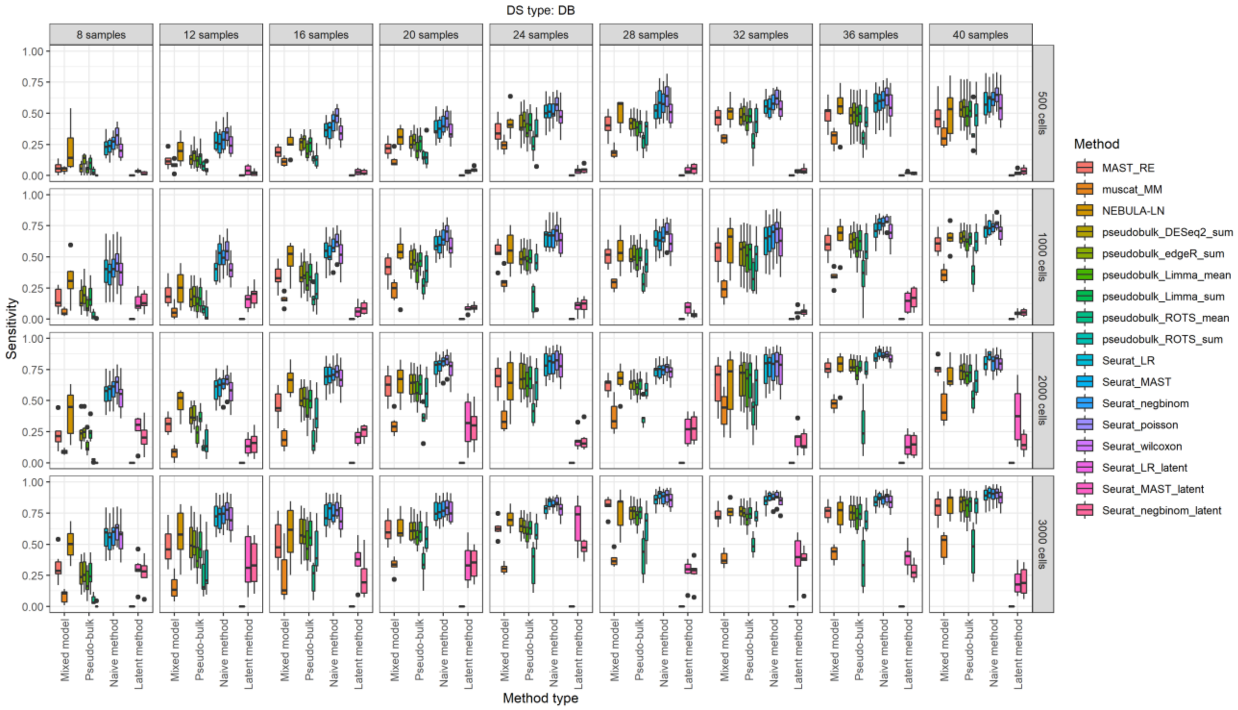


**Supplementary Figure 28. Sensitivity values for the reference-based negative binomial generative simulation (muscat).** The results are grouped in columns by the number of samples in the comparison and in the rows by the number of cells per subject. In this simulation, each sample includes three clusters. These results are for the differential state (DS) type, which includes changes in both modality and proportions (DB). FDR=0.05 was used as the cutoff to define the positives and negatives for calculating sensitivity, specificity and precision.

### Supplementary Figure 29

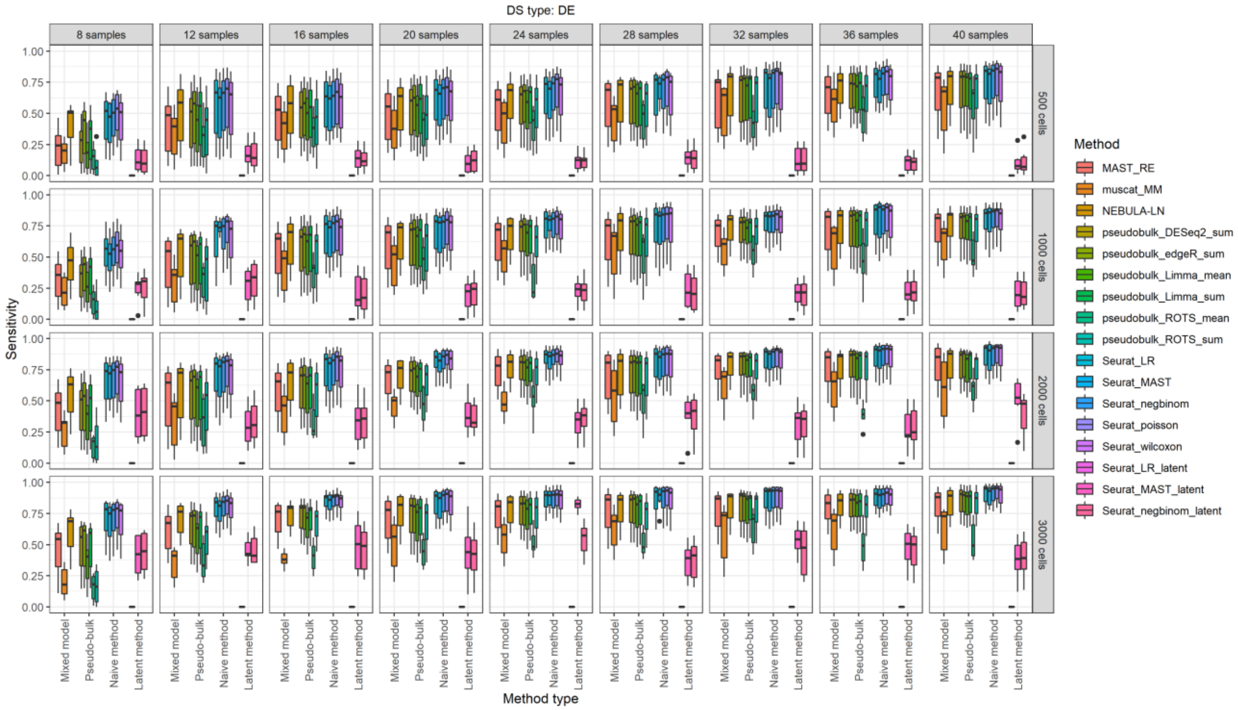


**Supplementary Figure 29. Sensitivity values for the reference-based negative binomial generative simulation (muscat).** The results are grouped in columns by the number of samples in the comparison and in the rows by the number of cells per subject. In this simulation, each sample includes three clusters. These results are for the differential state (DS) type, which includes changes in mean expression (DE). FDR=0.05 was used as the cutoff to define the positives and negatives for calculating sensitivity, specificity and precision.

### Supplementary Figure 30

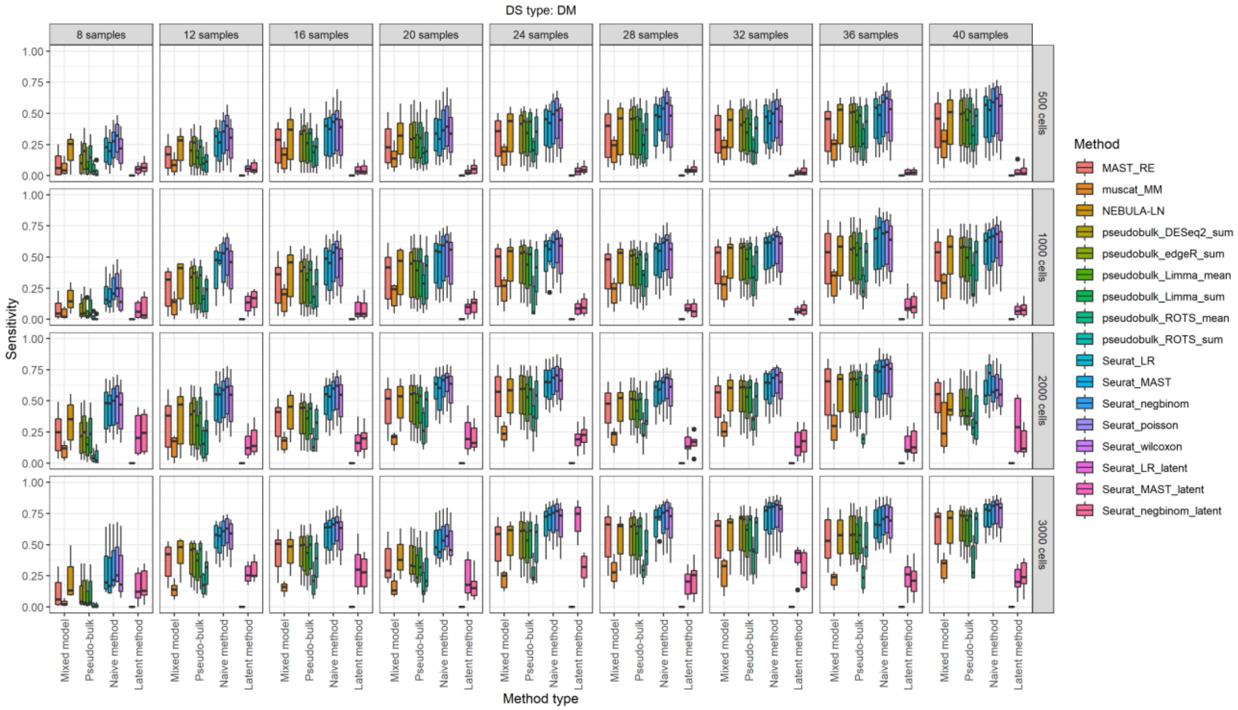


**Supplementary Figure 30. Sensitivity values for the reference-based negative binomial generative simulation (muscat).** The results are grouped in columns by the number of samples in the comparison and in the rows by the number of cells per subject. In this simulation, each sample includes three clusters. These results are for the differential state (DS) type, which includes changes in modality (DM). FDR=0.05 was used as the cutoff to define the positives and negatives for calculating sensitivity, specificity and precision.

### Supplementary Figure 31

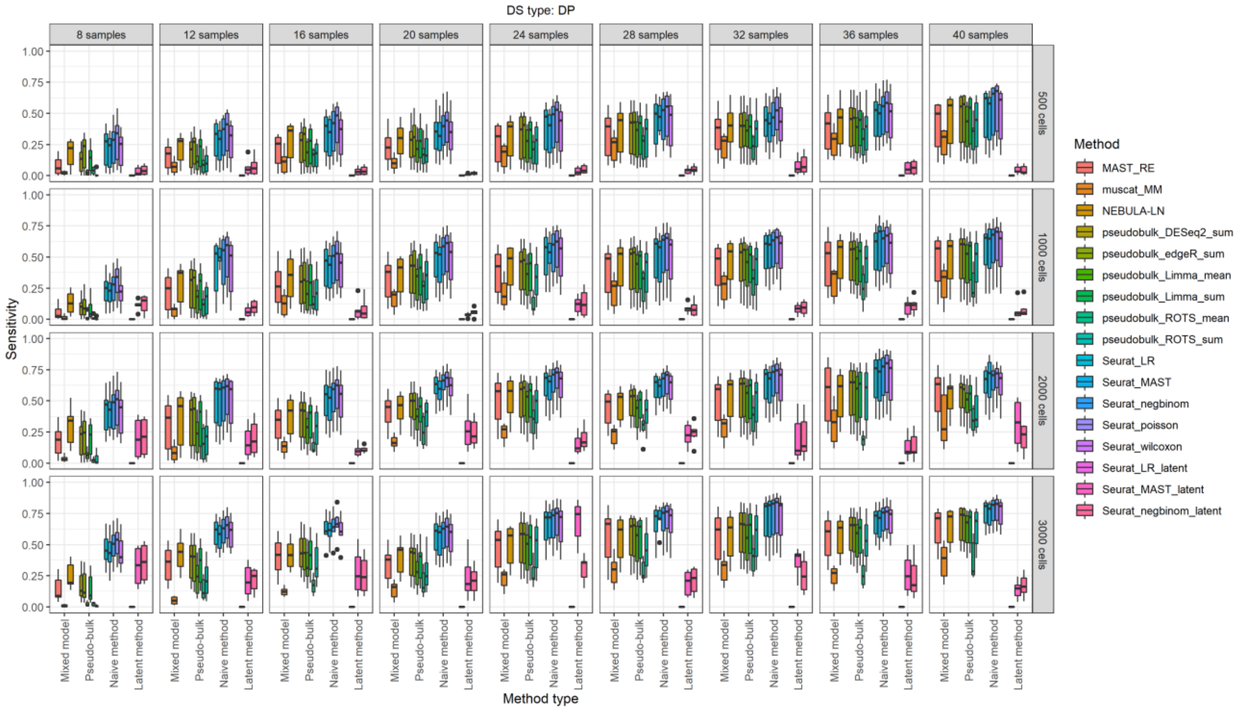


**Supplementary Figure 31. Sensitivity values for the reference-based negative binomial generative simulation (muscat).** The results are grouped in columns by the number of samples in the comparison and in the rows by the number of cells per subject. In this simulation, each sample includes three clusters. These results are for the differential state (DS) type, which includes changes in the proportions of low and high expression-state components (DP). FDR=0.05 was used as the cutoff to define the positives and negatives for calculating sensitivity, specificity and precision.

### Supplementary Figure 32

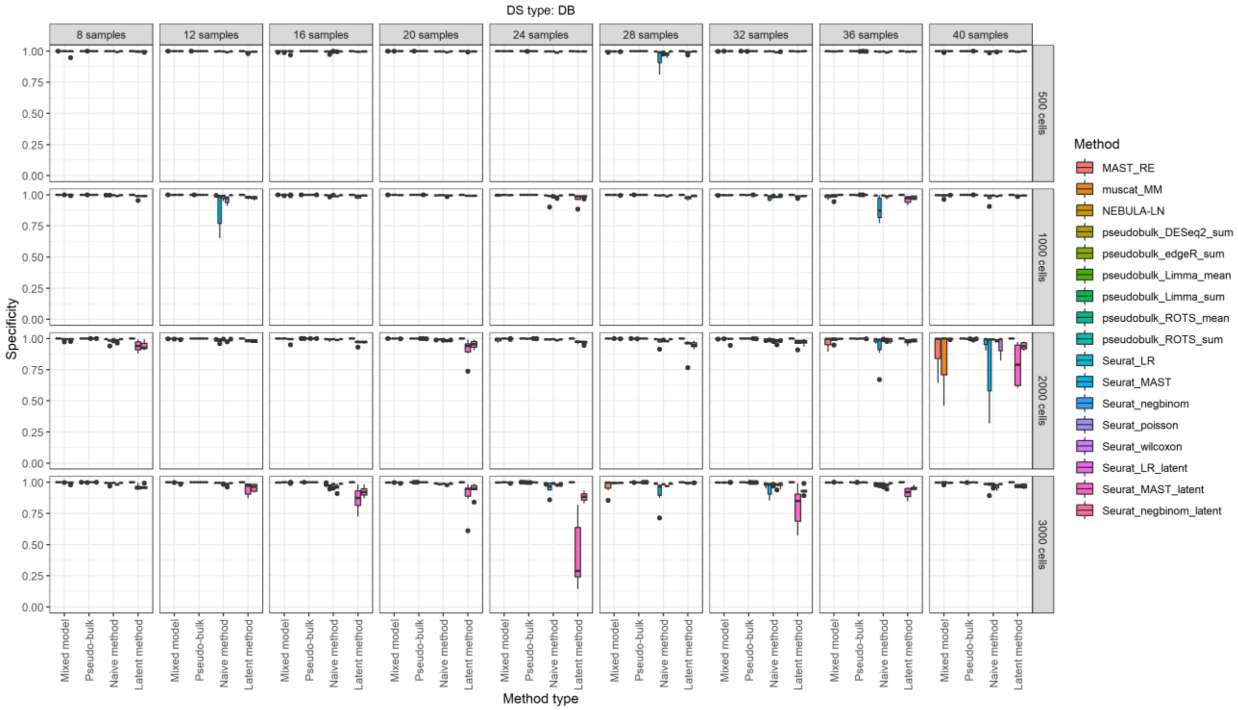


**Supplementary Figure 32. Specificity values for the reference-based negative binomial generative simulation (muscat).** The results are grouped in columns by the number of samples in the comparison and in the rows by the number of cells per subject. In this simulation, each sample includes three clusters. These results are for the differential state (DS) type, which includes changes in both modality and proportions (DB). FDR=0.05 was used as the cutoff to define the positives and negatives for calculating sensitivity, specificity and precision.

### Supplementary Figure 33

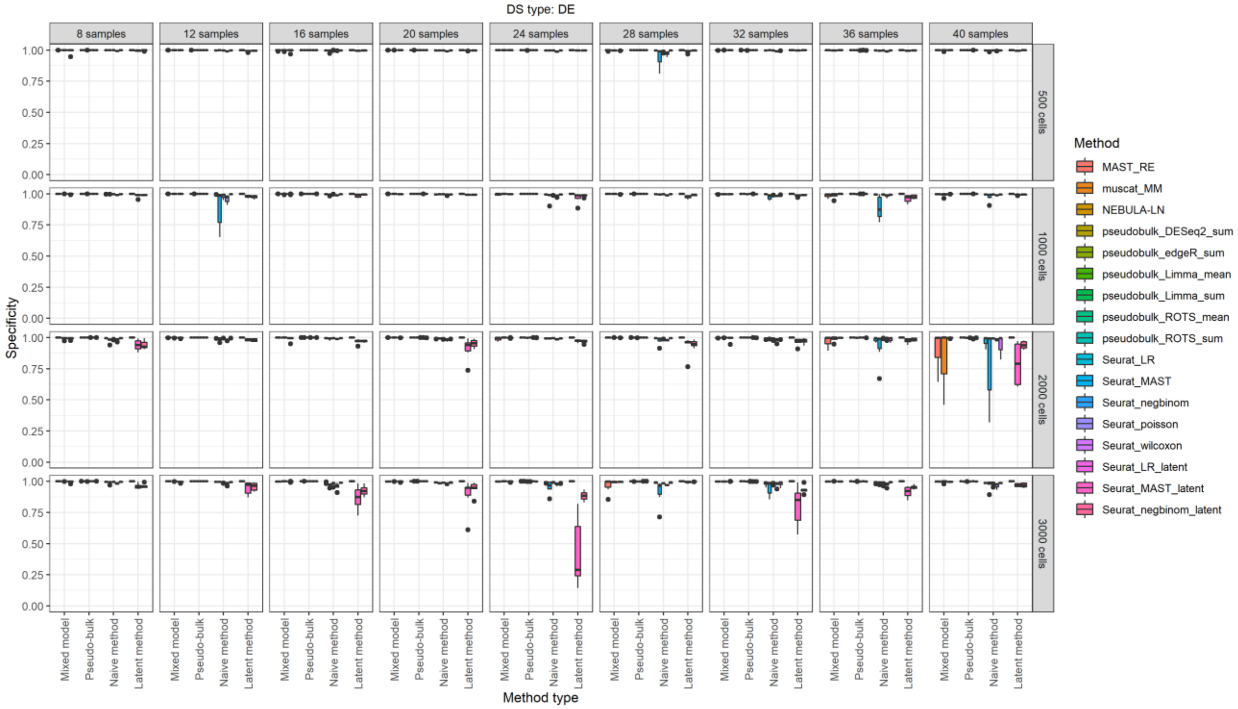


**Supplementary Figure 33. Specificity values for the reference-based negative binomial generative simulation (muscat).** The results are grouped in columns by the number of samples in the comparison and in the rows by the number of cells per subject. In this simulation, each sample includes three clusters. These results are for the differential state (DS) type, which includes changes in mean expression (DE). FDR=0.05 was used as the cutoff to define the positives and negatives for calculating sensitivity, specificity and precision.

### Supplementary Figure 34

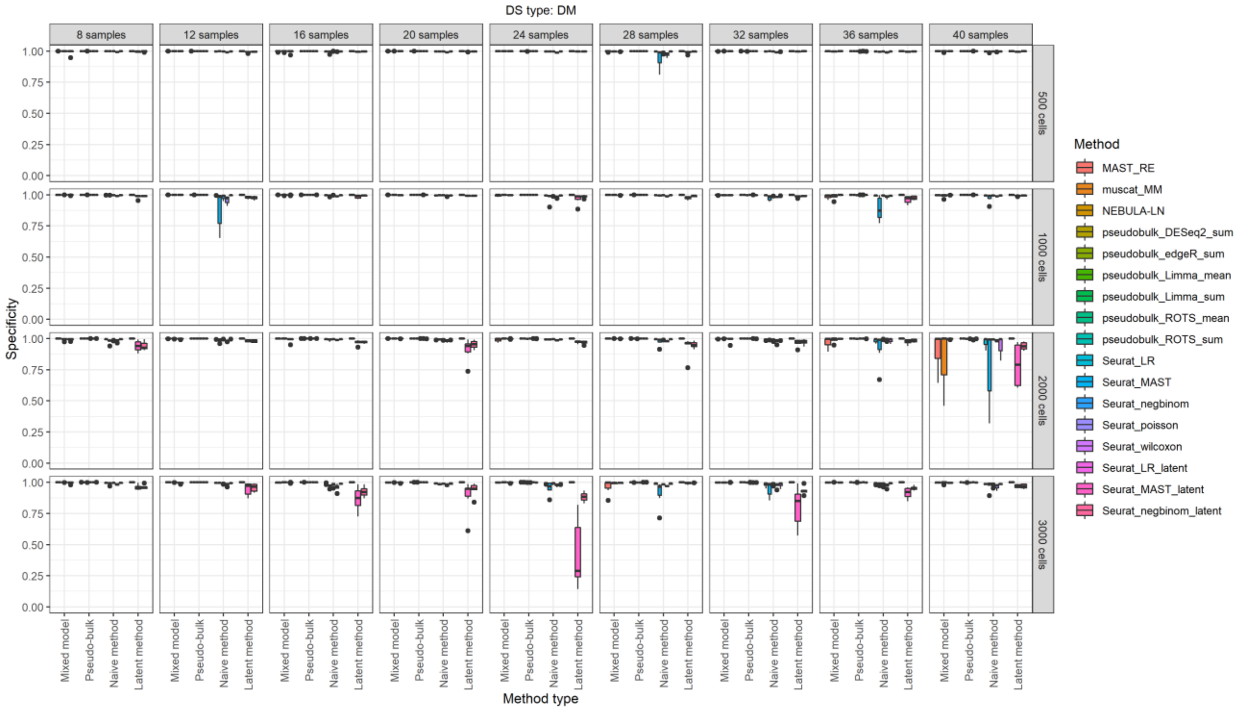


**Supplementary Figure 34. Specificity values for the reference-based negative binomial generative simulation (muscat).** The results are grouped in columns by the number of samples in the comparison and in the rows by the number of cells per subject. In this simulation, each sample includes three clusters. These results are for the differential state (DS) type, which includes changes in modality (DM). FDR=0.05 was used as the cutoff to define the positives and negatives for calculating sensitivity, specificity and precision.

### Supplementary Figure 35

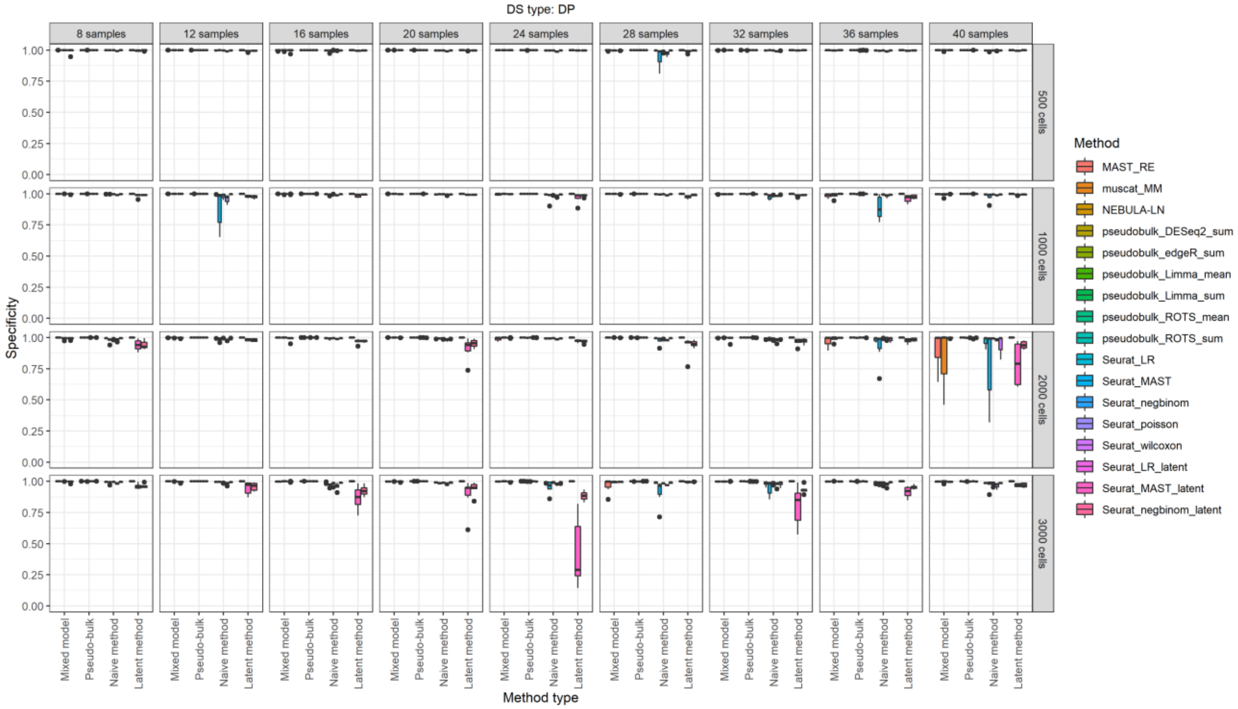


**Supplementary Figure 35. Specificity values for the reference-based negative binomial generative simulation (muscat).** The results are grouped in columns by the number of samples in the comparison and in the rows by the number of cells per subject. In this simulation, each sample includes three clusters. These results are for the differential state (DS) type, which includes changes in the proportions of low and high expression-state components (DP). FDR=0.05 was used as the cutoff to define the positives and negatives for calculating sensitivity, specificity and precision.

### Supplementary Figure 36

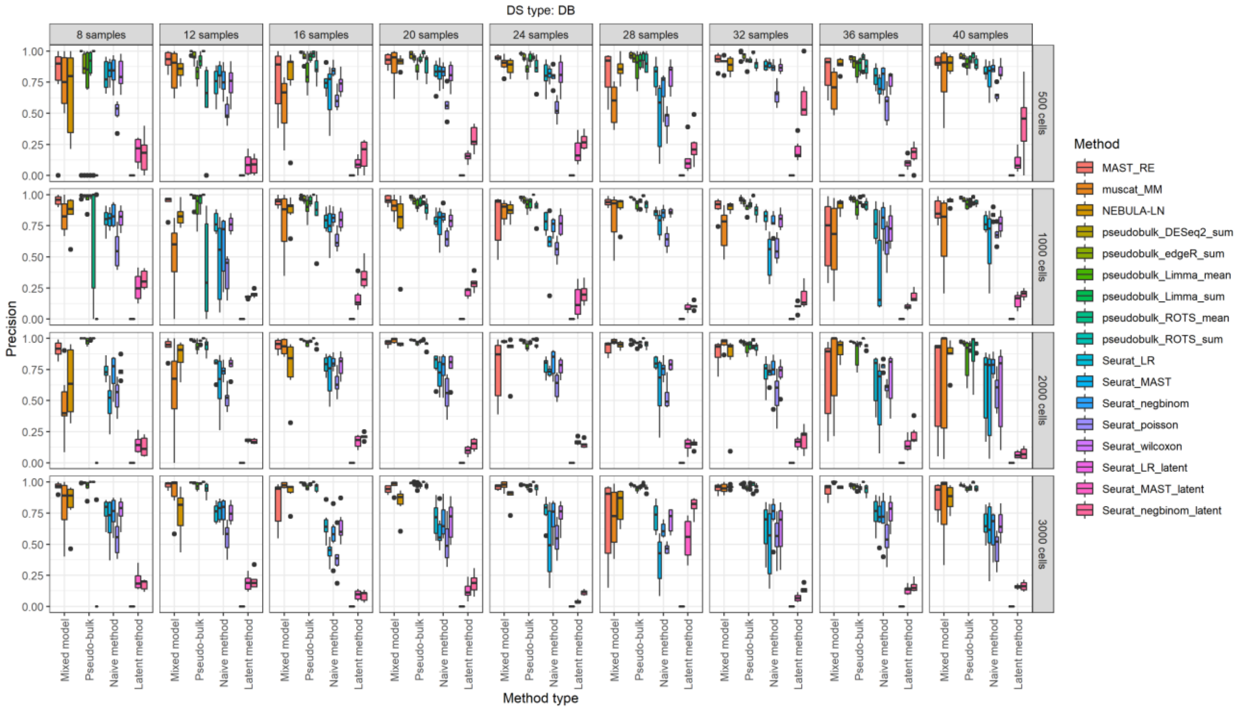


**Supplementary Figure 36. Precision values for the reference-based negative binomial generative simulation (muscat).** The results are grouped in columns by the number of samples in the comparison and in the rows by the number of cells per subject. In this simulation, each sample includes three clusters. These results are for the differential state (DS) type, which includes changes in both modality and proportions (DB). FDR=0.05 was used as the cutoff to define the positives and negatives for calculating sensitivity, specificity and precision.

### Supplementary Figure 37

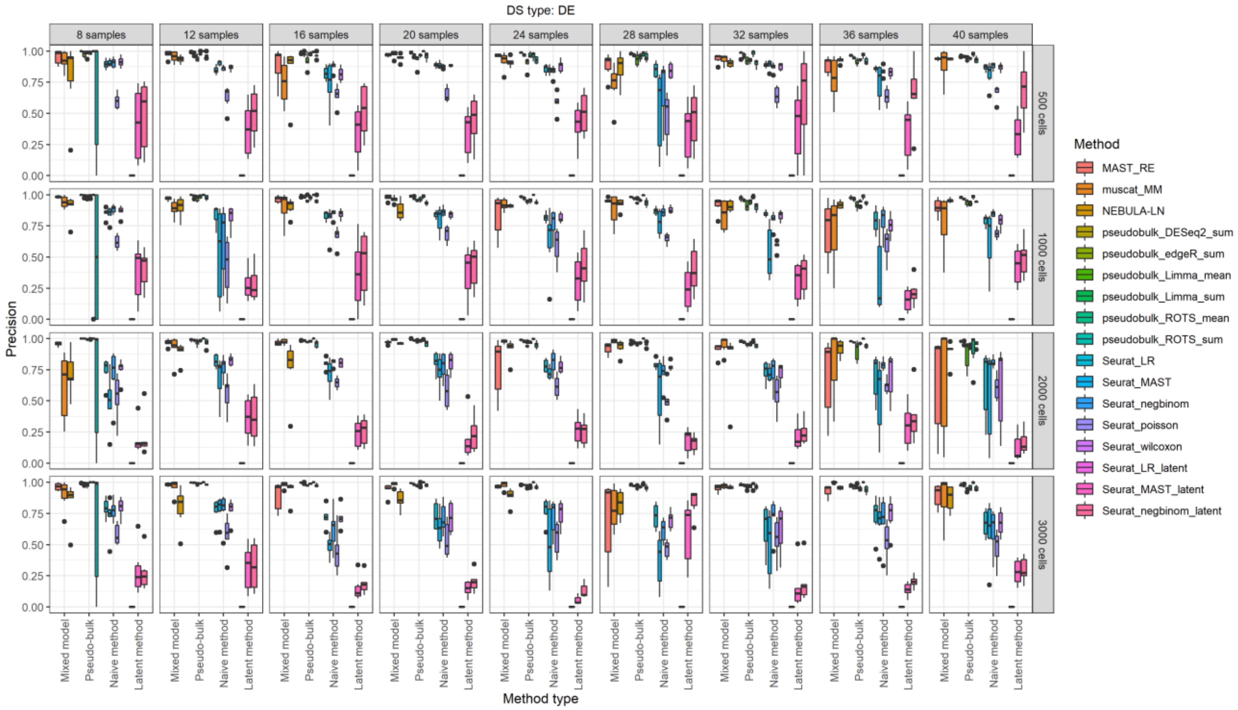


**Supplementary Figure 37. Precision values for the reference-based negative binomial generative simulation (muscat).** The results are grouped in columns by the number of samples in the comparison and in the rows by the number of cells per subject. In this simulation, each sample includes three clusters. These results are for the differential state (DS) type, which includes changes in mean expression (DE). FDR=0.05 was used as the cutoff to define the positives and negatives for calculating sensitivity, specificity and precision. See more details on how the data was simulated in **Section 2.2.1** of the manuscript.

### Supplementary Figure 38

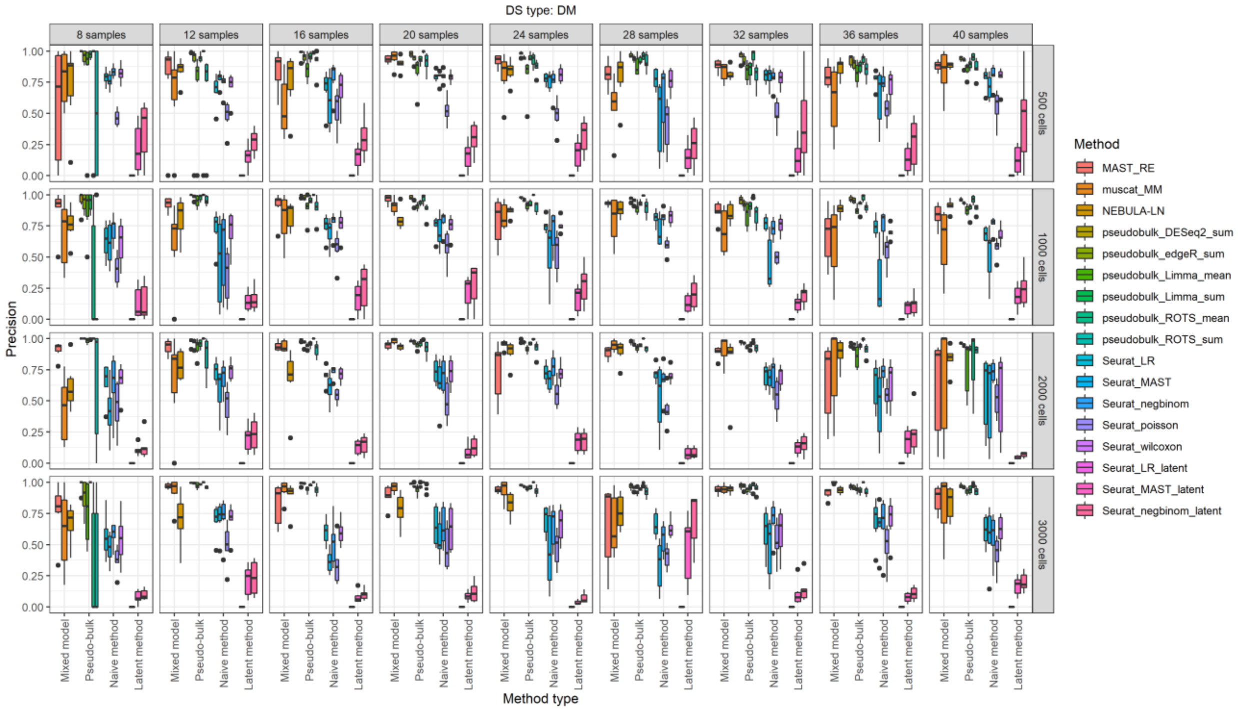


**Supplementary Figure 38. Precision values for the reference-based negative binomial generative simulation (muscat).** The results are grouped in columns by the number of samples in the comparison and in the rows by the number of cells per subject. In this simulation, each sample includes three clusters. These results are for the differential state (DS) type, which includes changes in modality (DM). FDR=0.05 was used as the cutoff to define the positives and negatives for calculating sensitivity, specificity and precision.

### Supplementary Figure 39

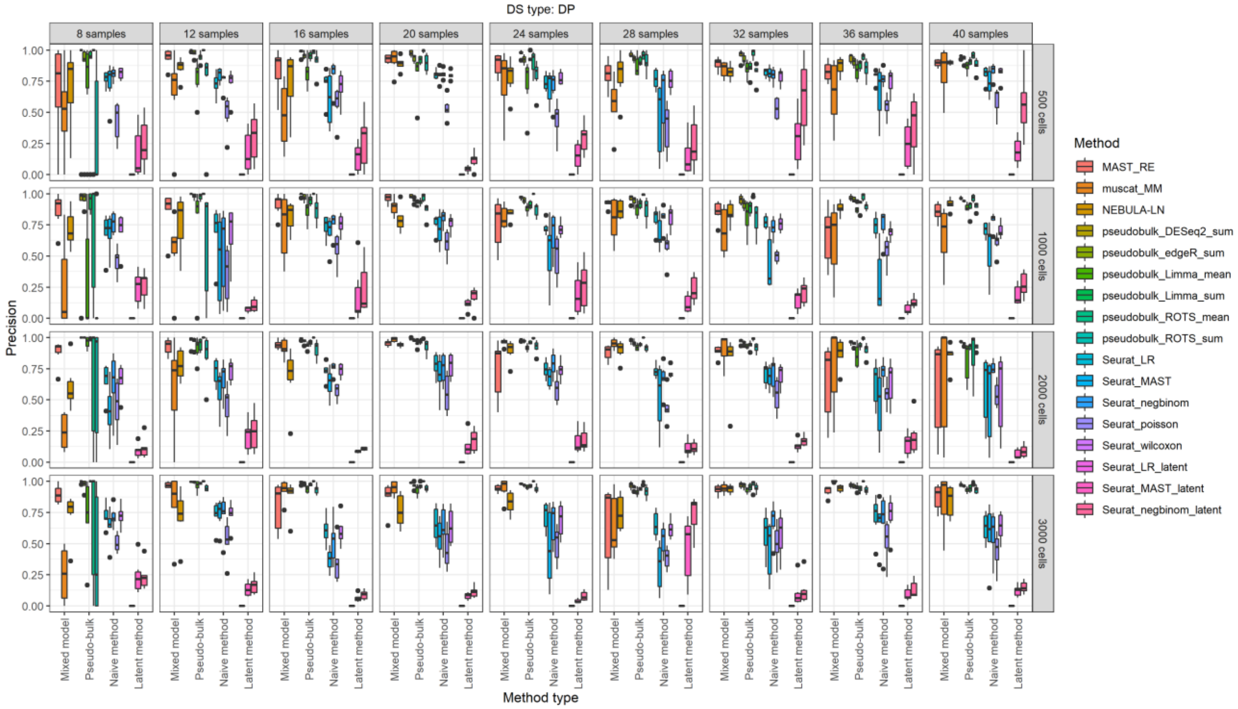


**Supplementary Figure 39. Precision values for the reference-based negative binomial generative simulation (muscat).** The results are grouped in columns by the number of samples in the comparison and in the rows by the number of cells per subject. In this simulation, each sample includes three clusters. These results are for the differential state (DS) type, which includes changes in the proportions of low and high expression-state components (DP). FDR=0.05 was used as the cutoff to define the positives and negatives for calculating sensitivity, specificity and precision. See more details on how the data was simulated in **Section 2.2.1** of the manuscript.

### Supplementary Figure 40

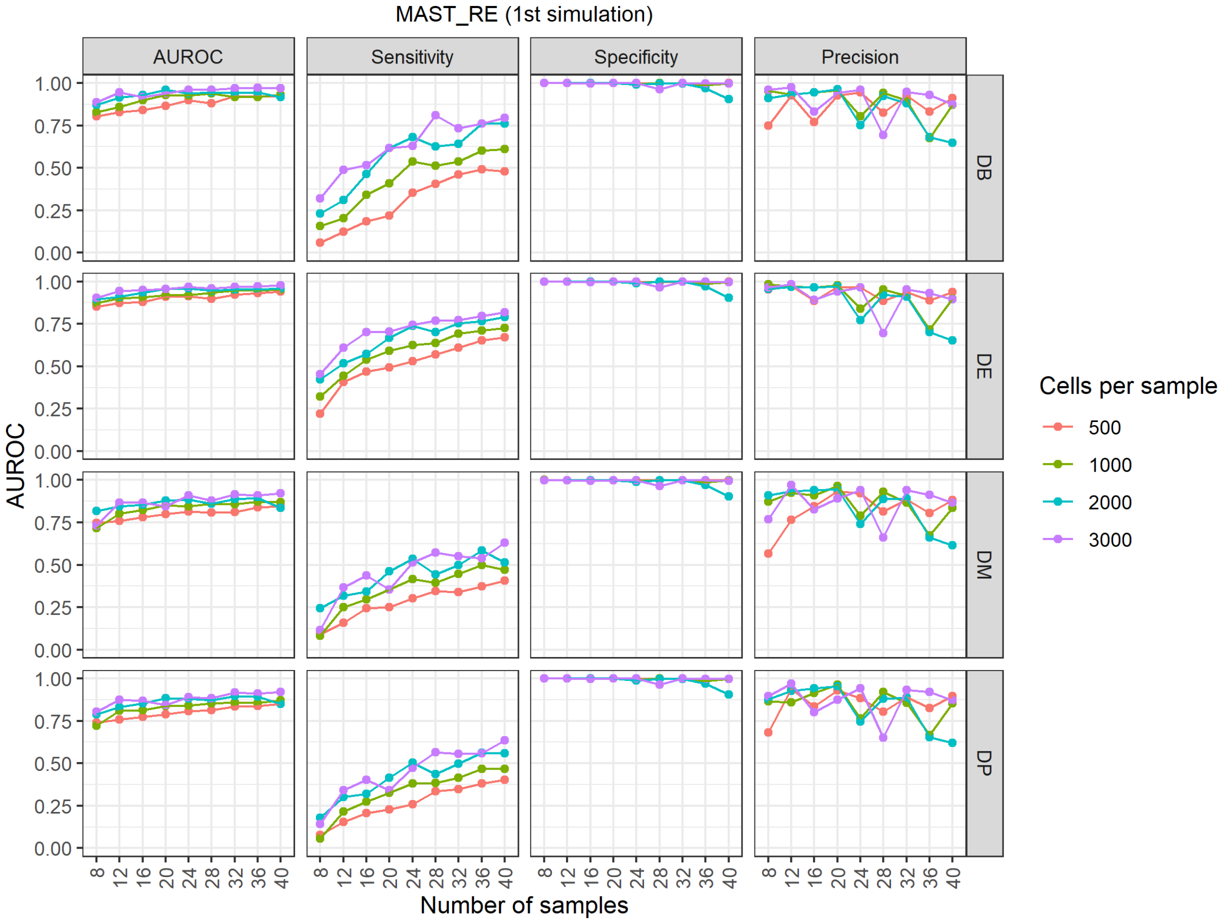


**Supplementary Figure 40. Results of the cell-sample-extended reference-based negative binomial generative simulation for MAST_RE.**

.

### Supplementary Figure 41

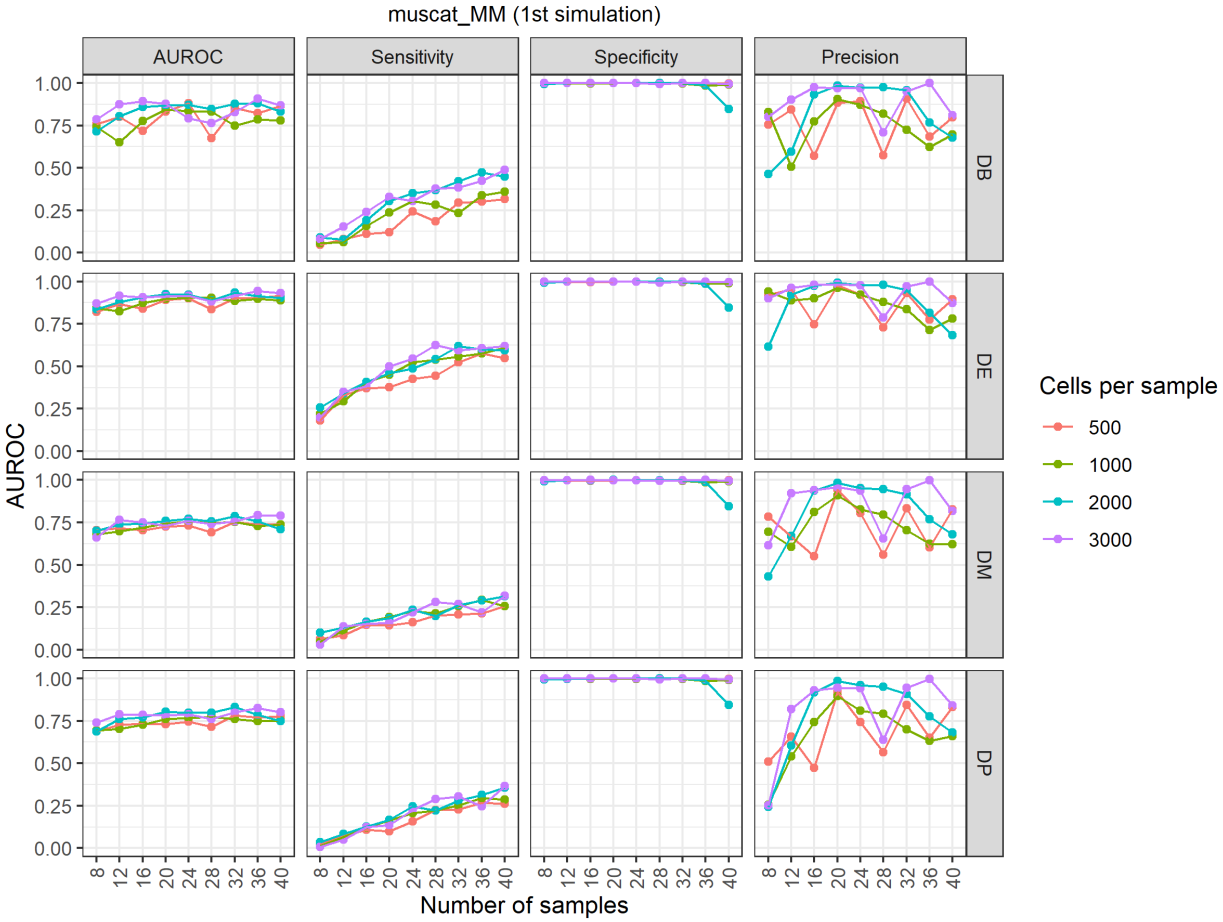


**Supplementary Figure 41. Results of the cell-sample-extended reference-based negative binomial generative simulation for muscat_MM.**

### Supplementary Figure 42

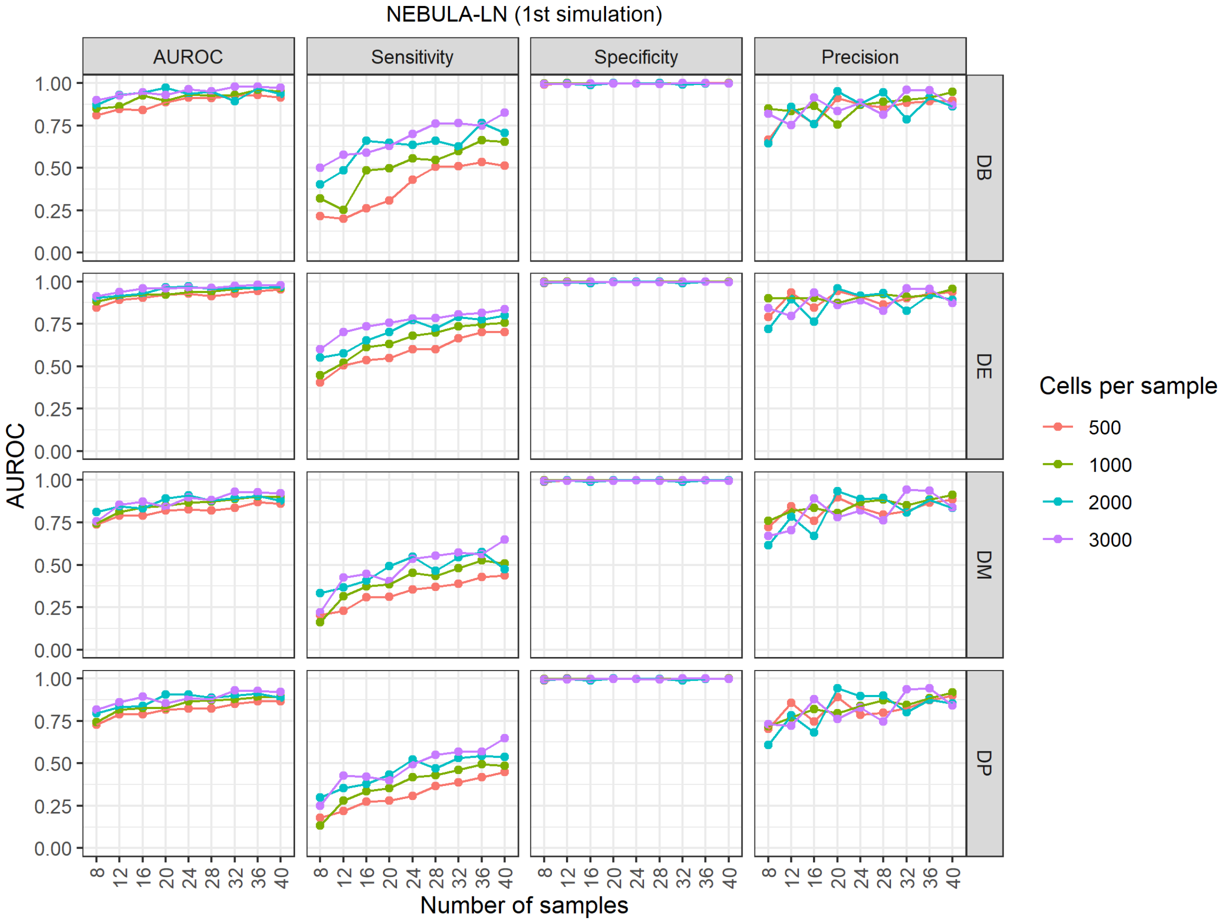


**Supplementary Figure 42. Results of the cell-sample-extended reference-based negative binomial generative simulation for NEBULA-LN.**

### Supplementary Figure 43

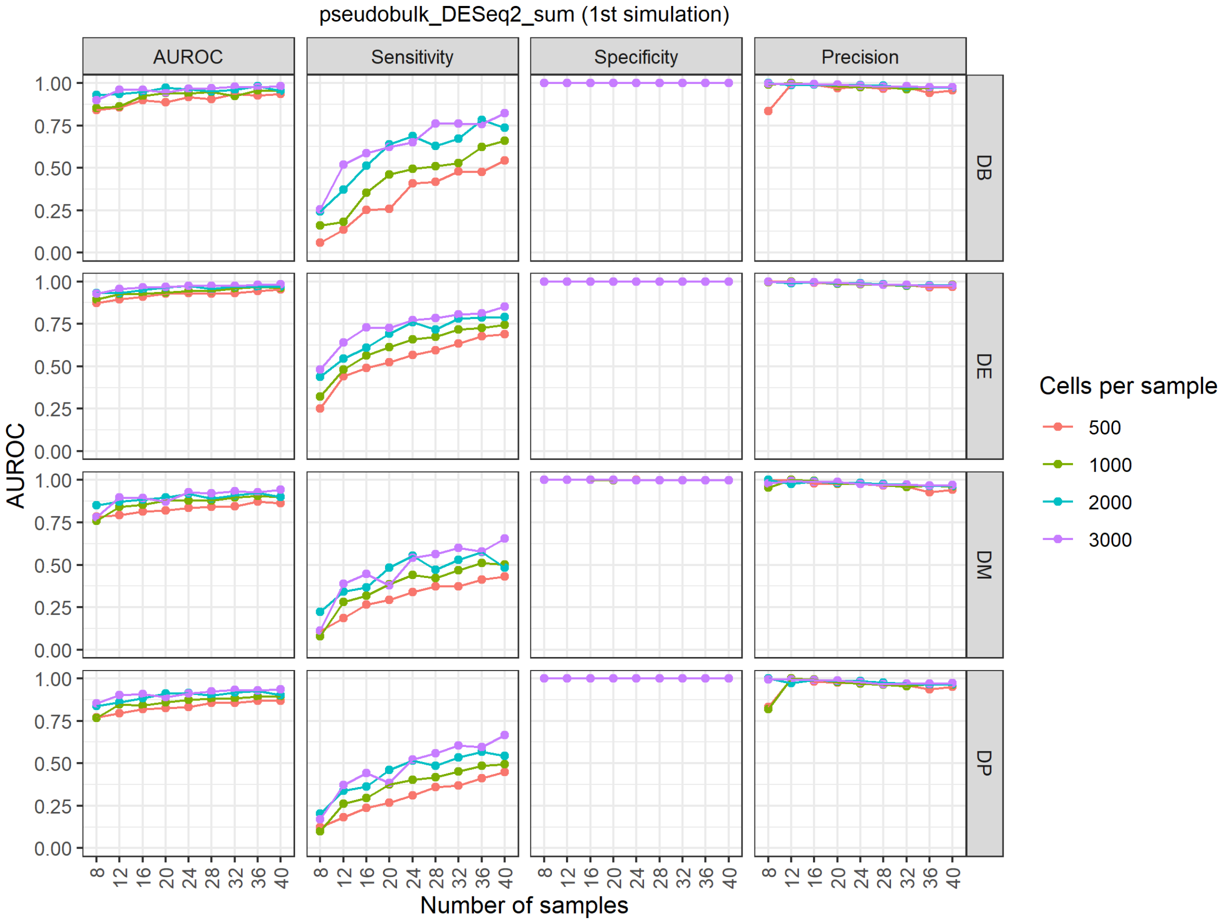


**Supplementary Figure 43. Results of the cell-sample-extended reference-based negative binomial generative simulation for pseudobulk_DESeq2_sum.**

### Supplementary Figure 44

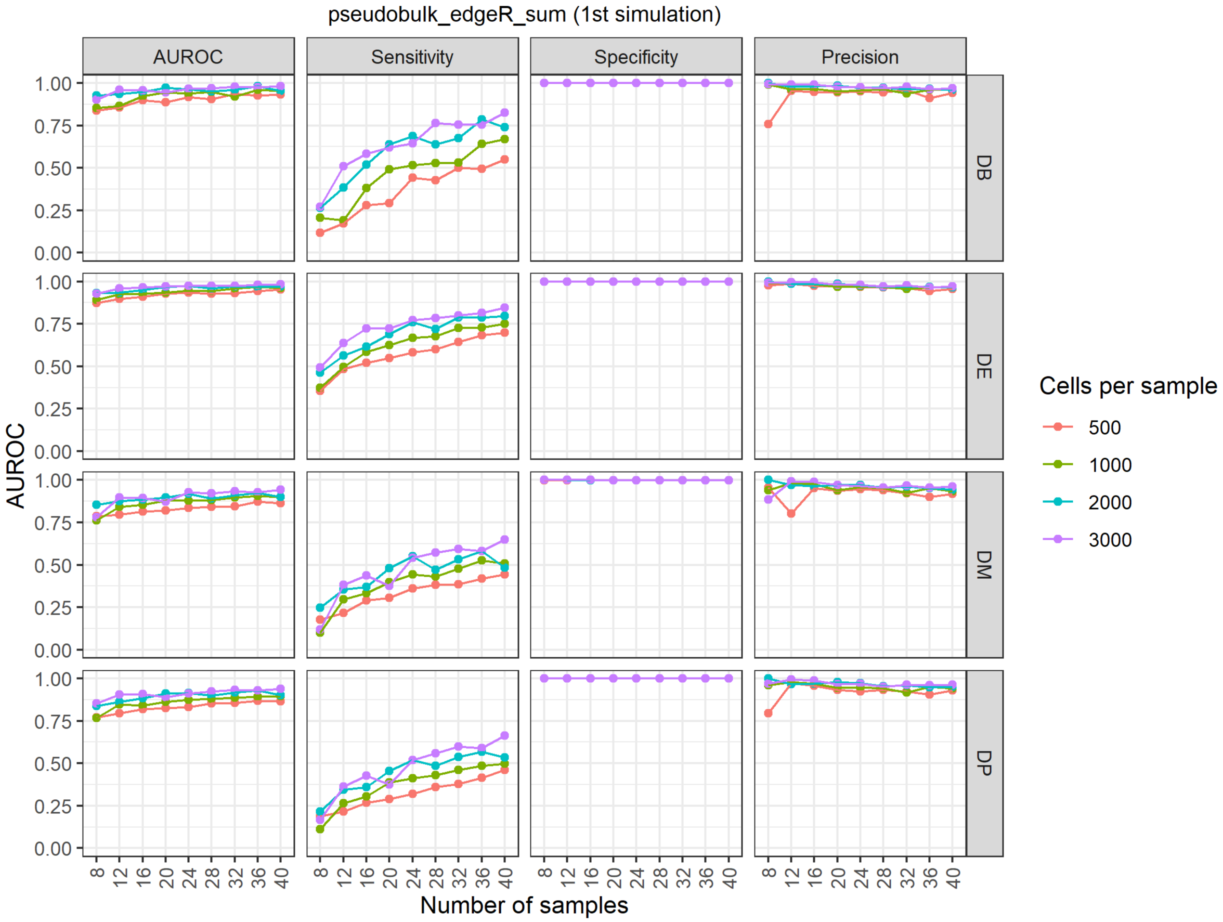


**Supplementary Figure 44. Results of the cell-sample-extended reference-based negative binomial generative simulation for pseudobulk_edgeR_sum.**

### Supplementary Figure 45

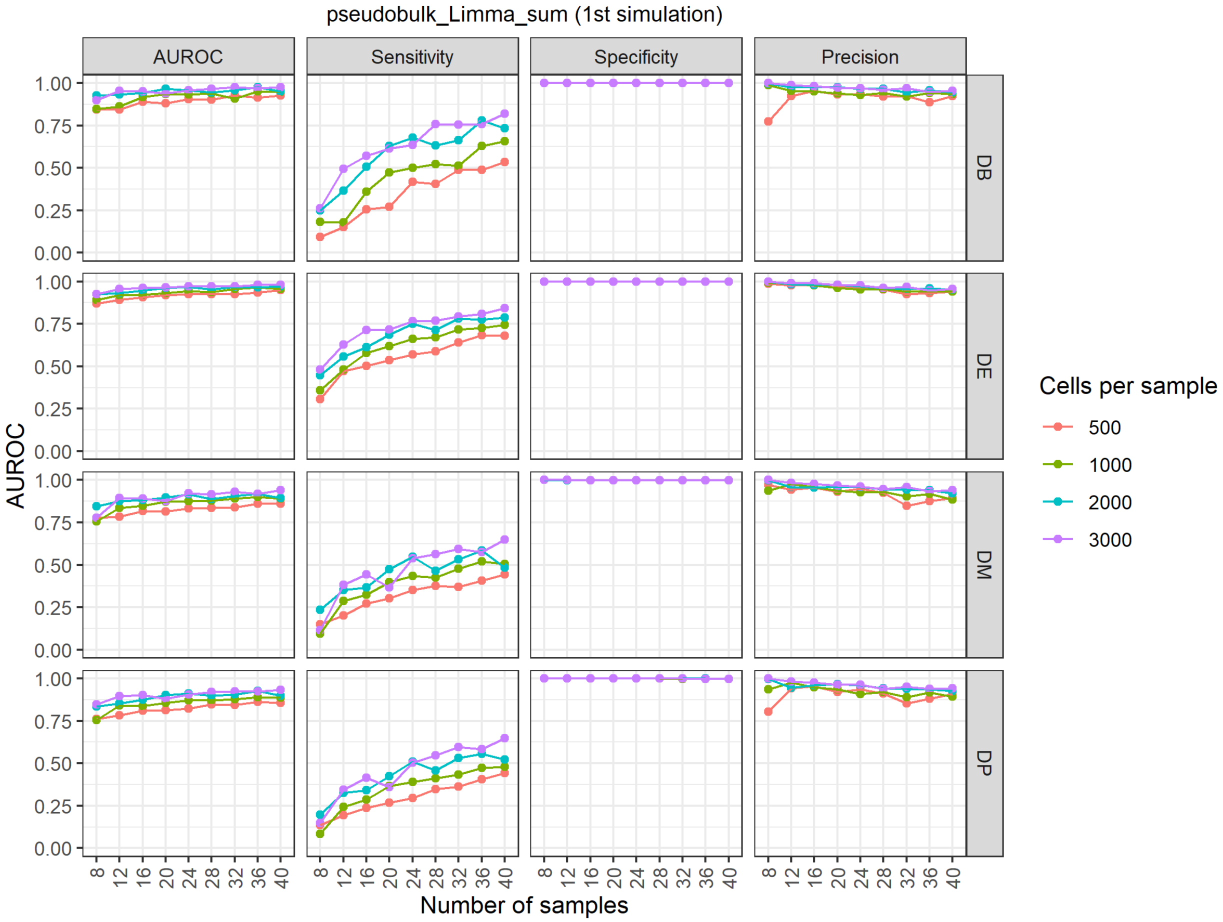


**Supplementary Figure 45. Results of the cell-sample-extended reference-based negative binomial generative simulation for pseudobulk_Limma_sum.**

### Supplementary Figure 46

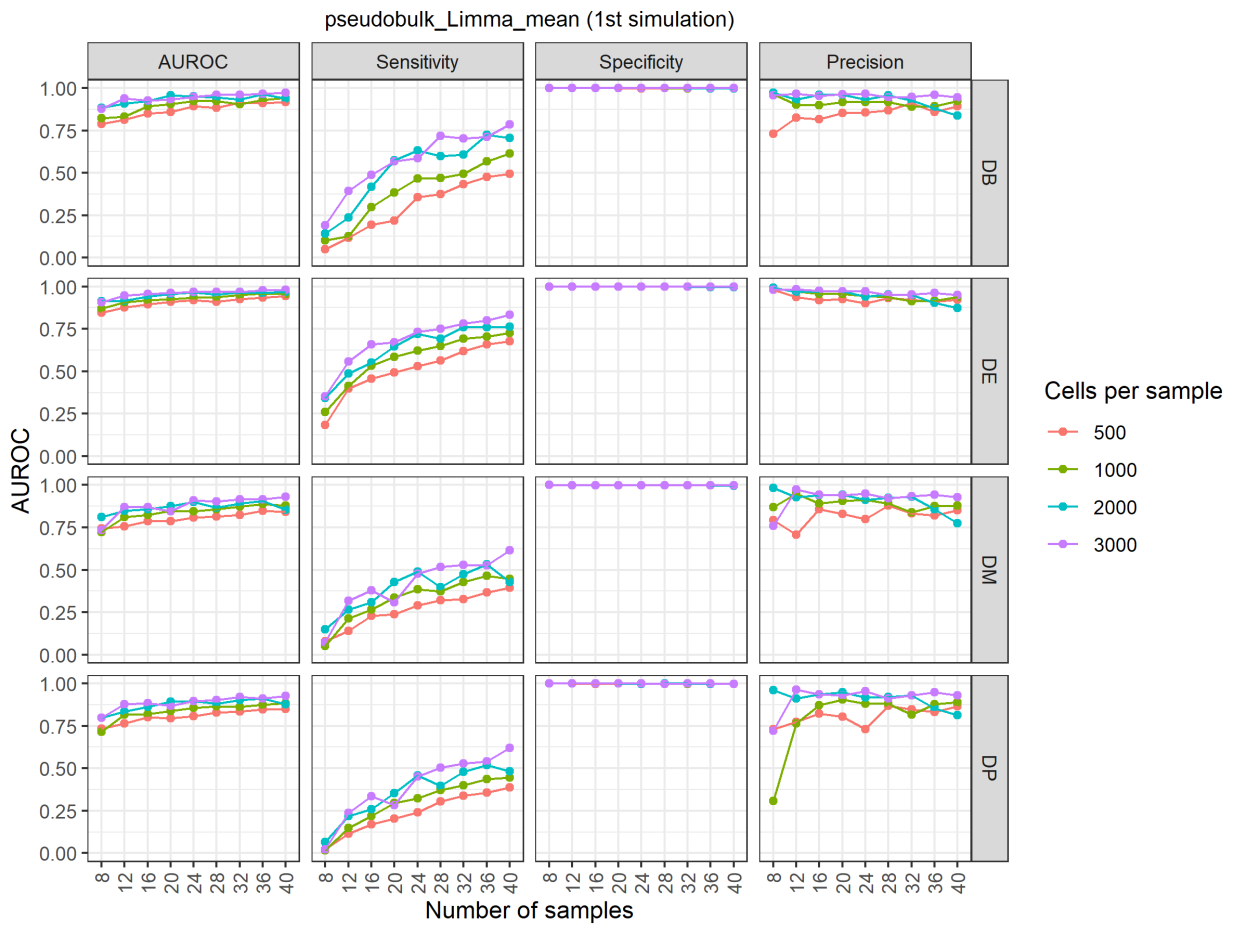


**Supplementary Figure 46. Results of the cell-sample-extended reference-based negative binomial generative simulation for pseudobulk_Limma_mean.**

### Supplementary Figure 47

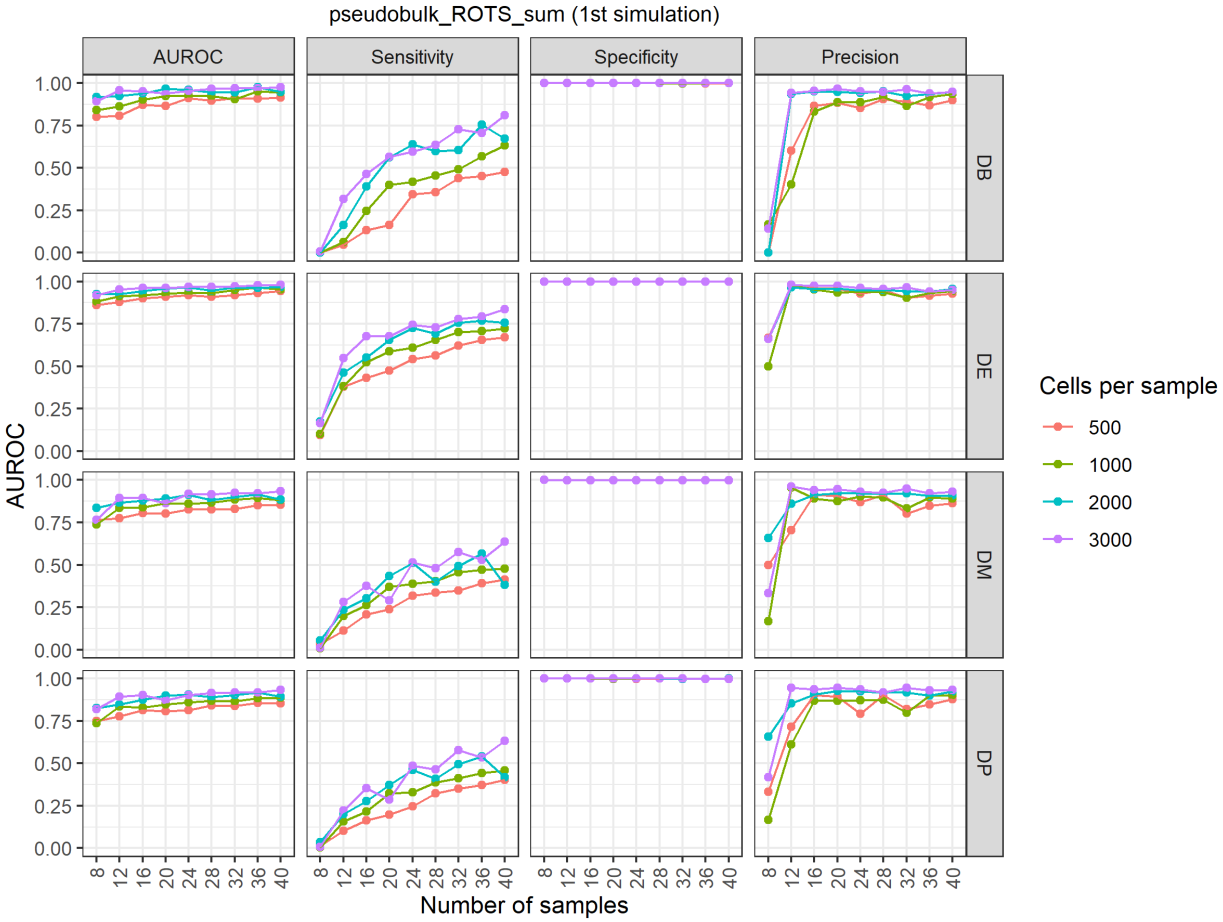


**Supplementary Figure 47. Results of the cell-sample-extended reference-based negative binomial generative simulation for pseudobulk_ROTS_sum.**

### Supplementary Figure 48

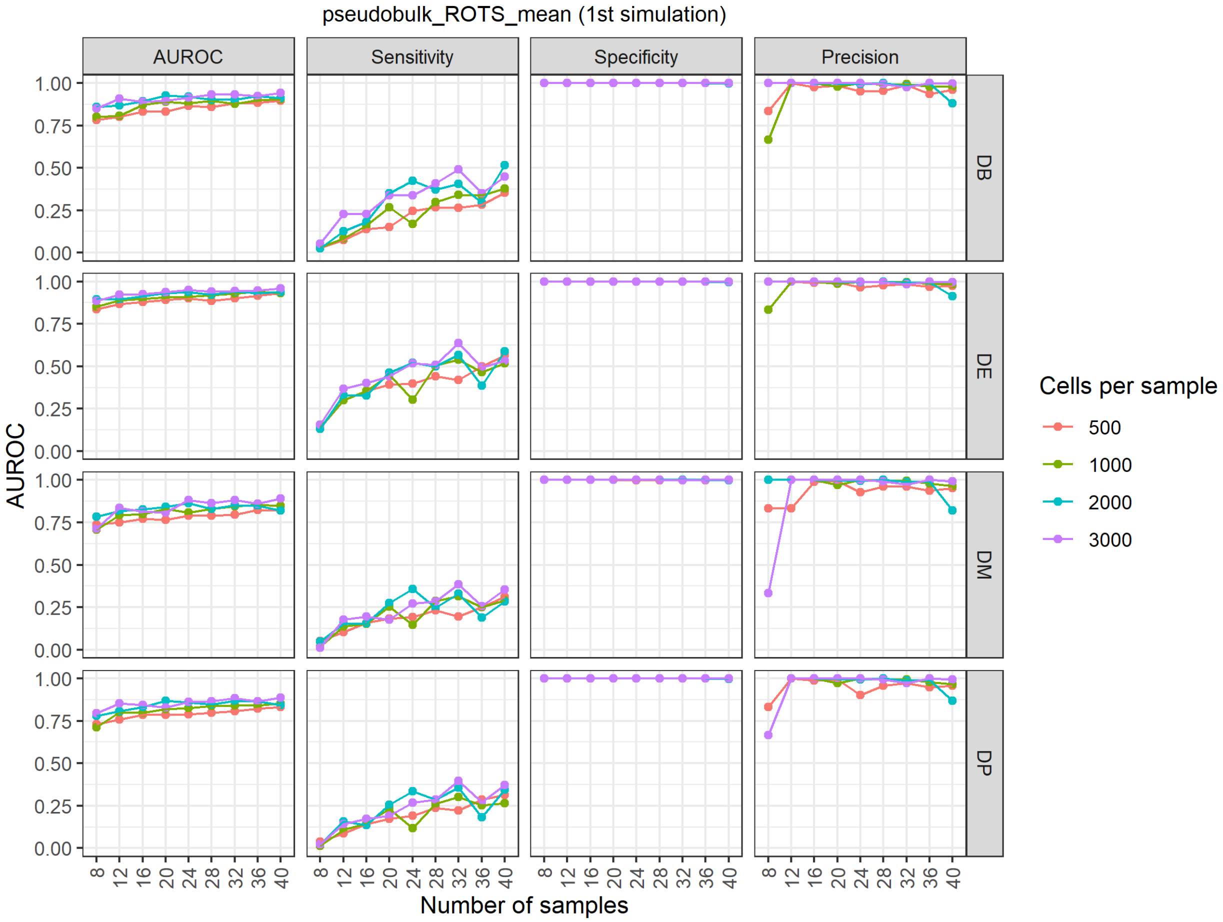


**Supplementary Figure 48. Results of the cell-sample-extended reference-based negative binomial generative simulation for pseudobulk_ROTS_mean.**

### Supplementary Figure 49

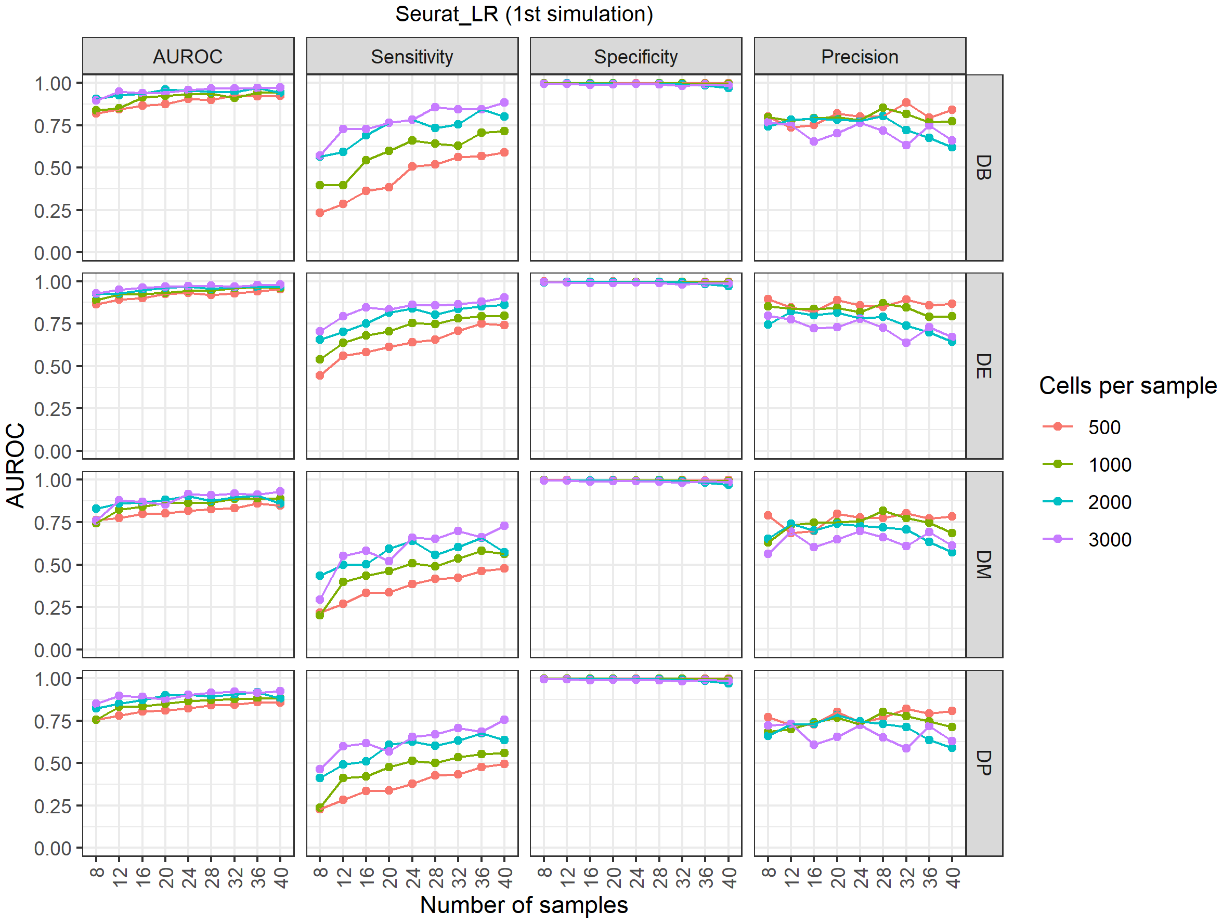


**Supplementary Figure 49. Results of the cell-sample-extended reference-based negative binomial generative simulation for Seurat_LR.**

### Supplementary Figure 50

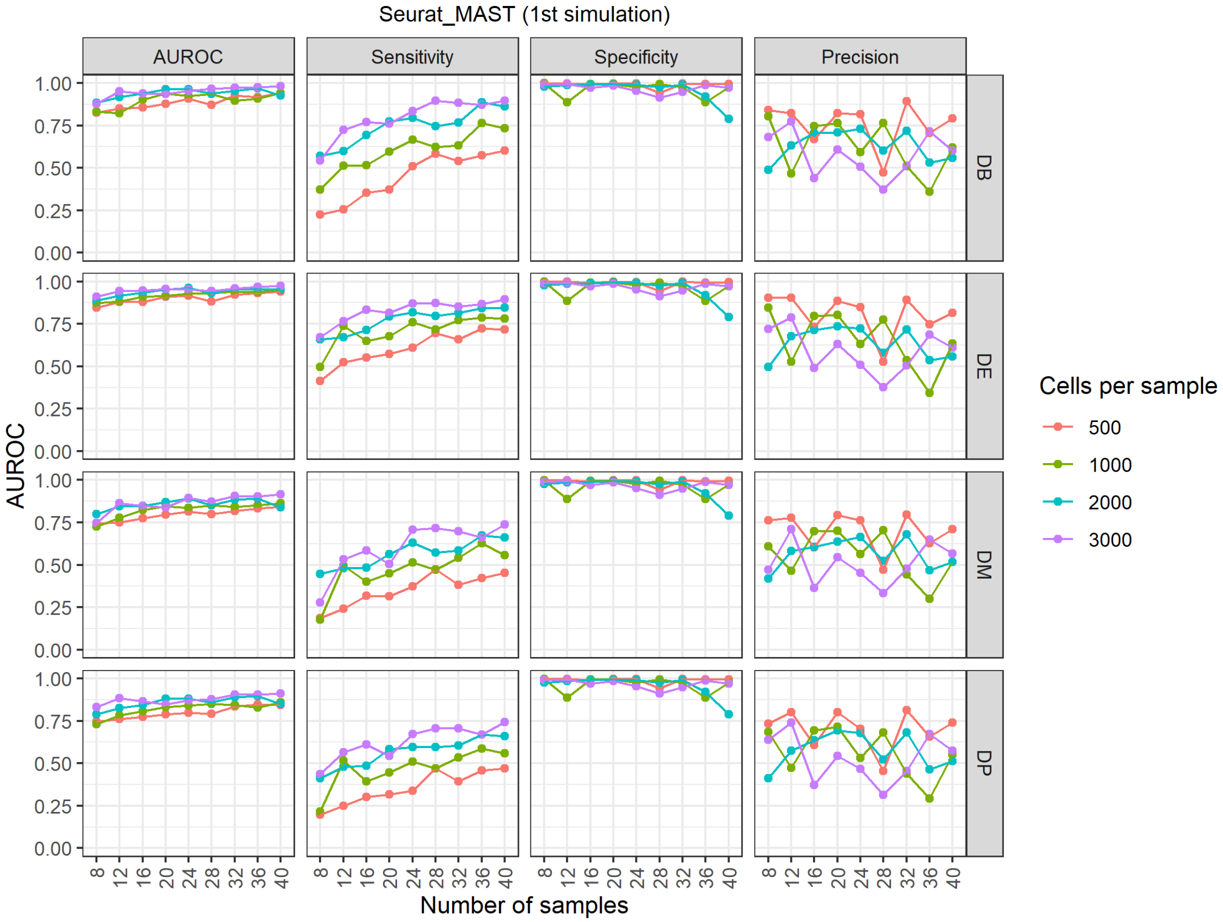


**Supplementary Figure 50. Results of the cell-sample-extended reference-based negative binomial generative simulation for Seurat_MAST.**

### Supplementary Figure 51

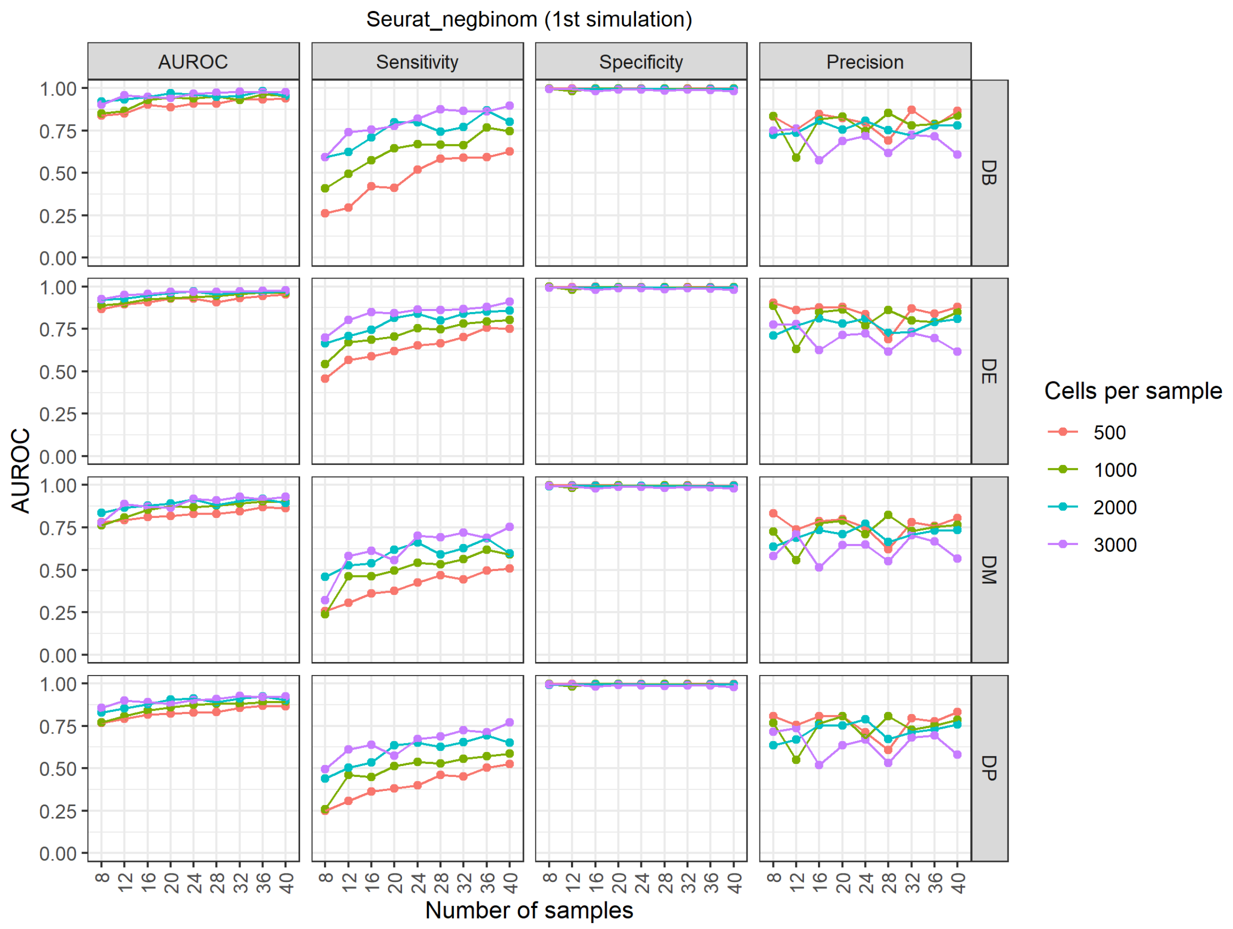


**Supplementary Figure 51. Results of the cell-sample-extended reference-based negative binomial generative simulation for Seurat_negbinom.**

### Supplementary Figure 52

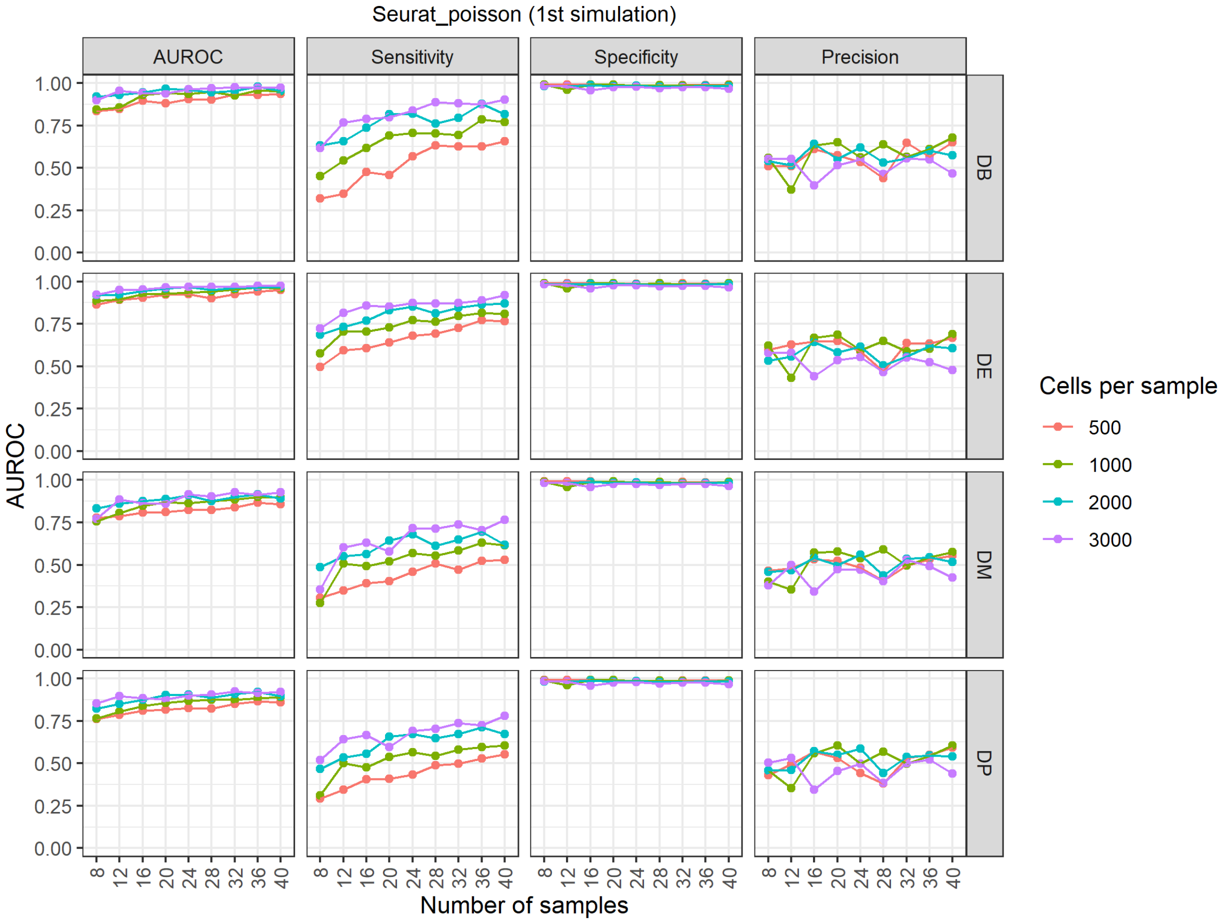


**Supplementary Figure 52. Results of the cell-sample-extended reference-based negative binomial generative simulation for Seurat_poisson.**

### Supplementary Figure 53

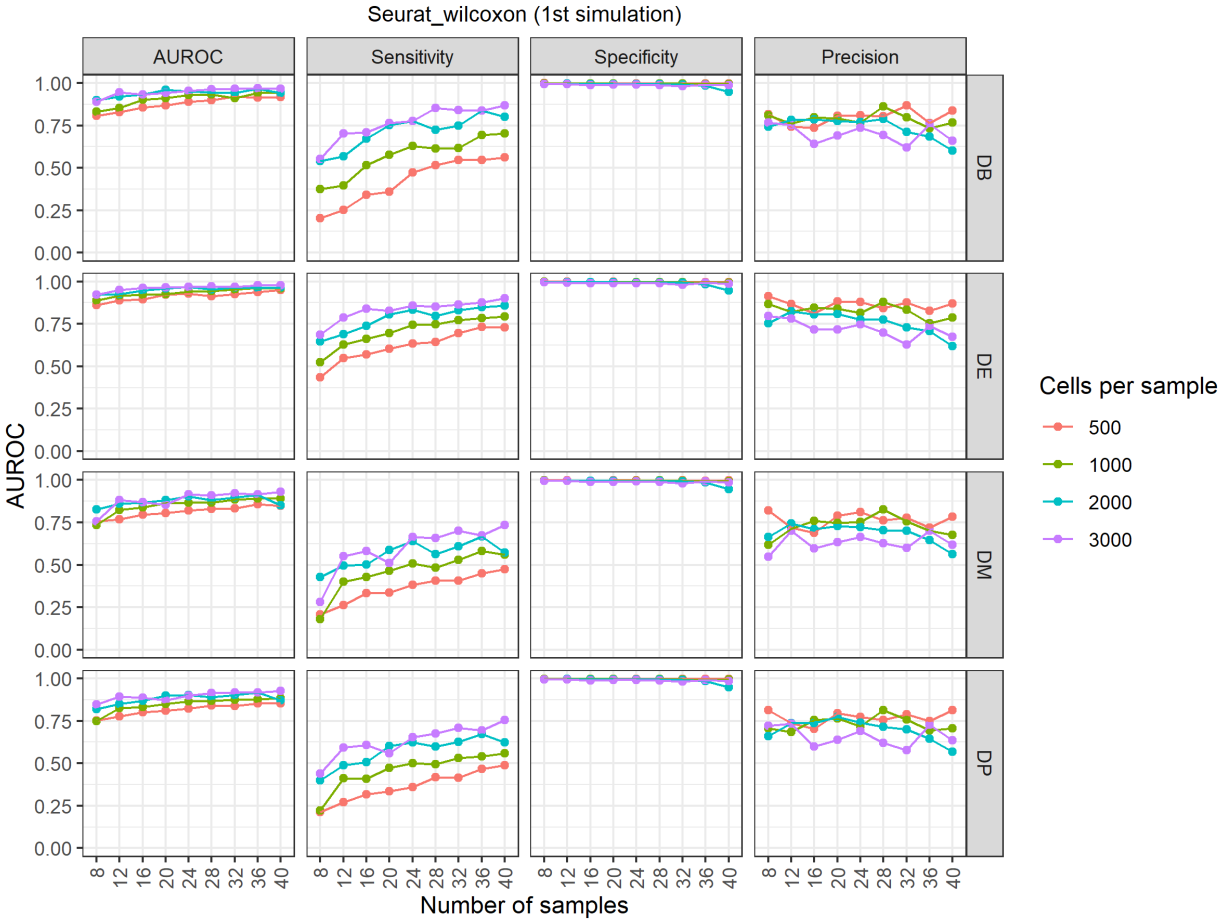


**Supplementary Figure 53. Results of the cell-sample-extended reference-based negative binomial generative simulation for Seurat_wilcoxon.**

### Supplementary Figure 54

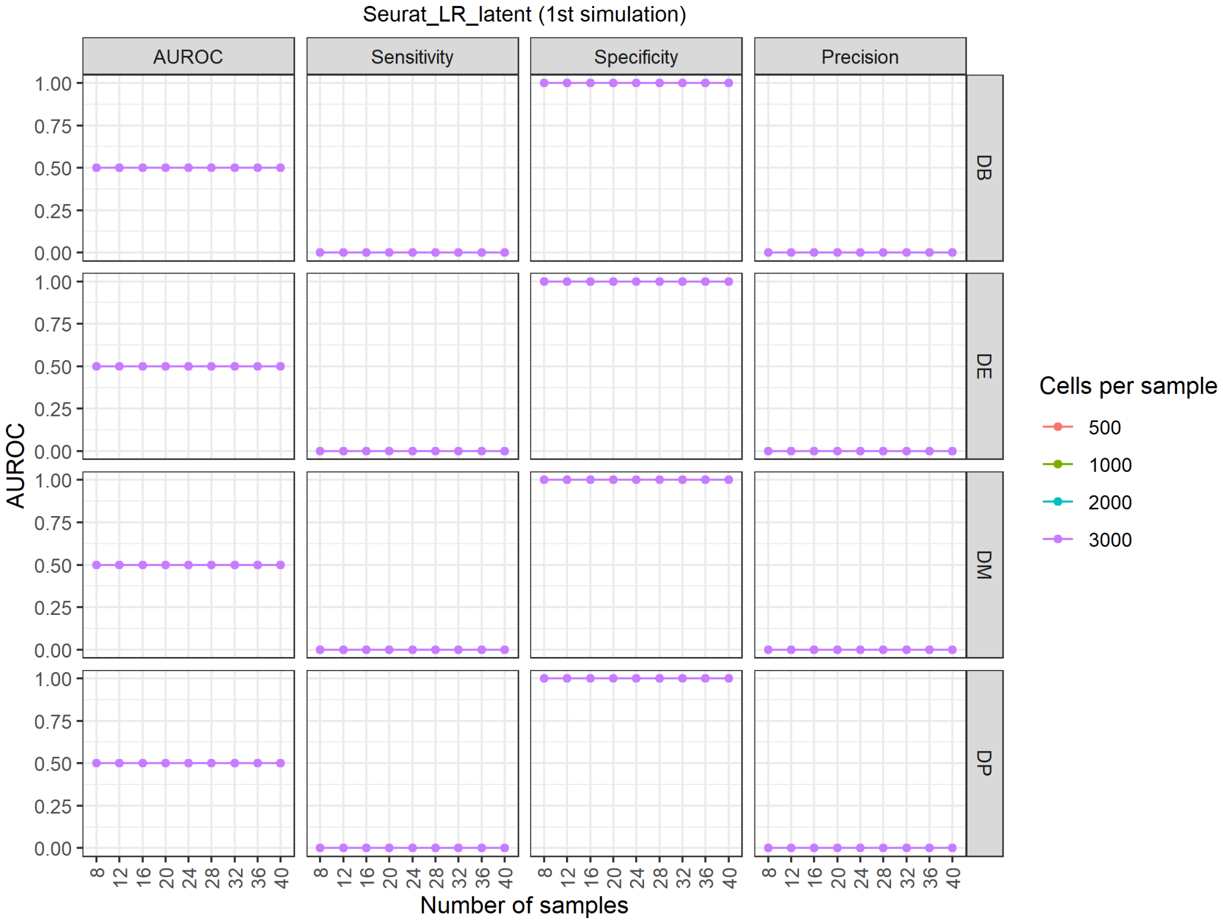


**Supplementary Figure 54. Results of the cell-sample-extended reference-based negative binomial generative simulation for Seurat_LR_latent.**

### Supplementary Figure 55

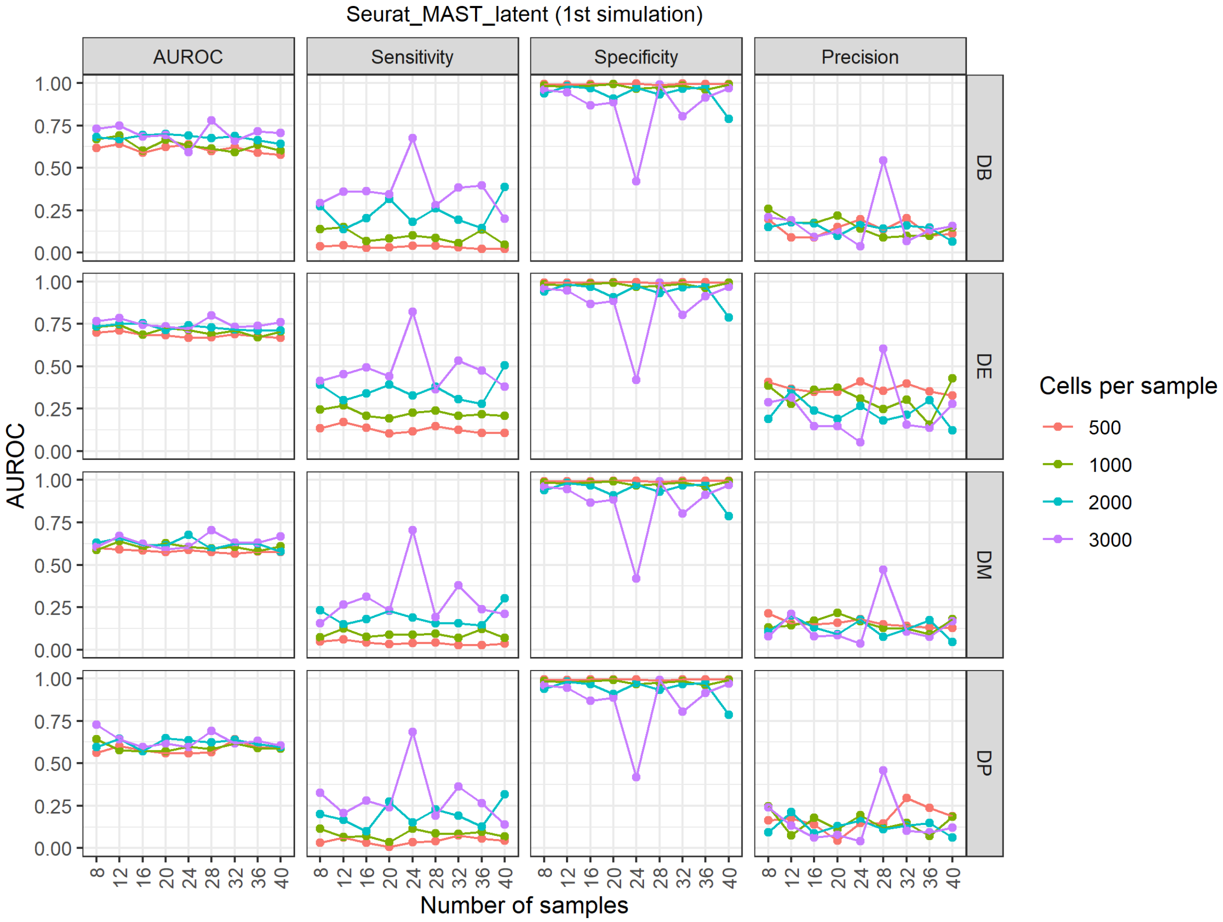


**Supplementary Figure 55. Results of the cell-sample-extended reference-based negative binomial generative simulation for Seurat_MAST_latent.**

### Supplementary Figure 56

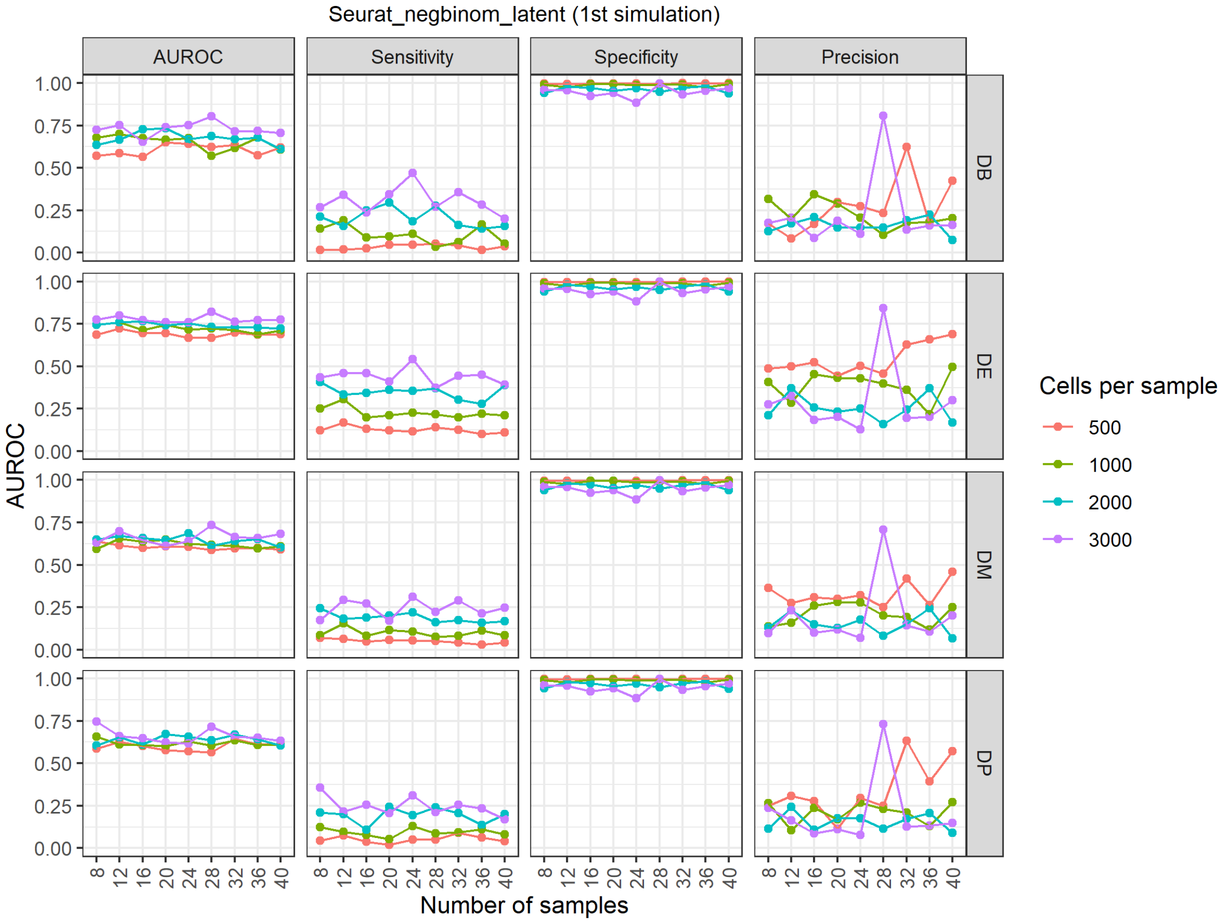


**Supplementary Figure 56. Results of the cell-sample-extended reference-based negative binomial generative simulation for Seurat_negbinom_latent.**

### Supplementary Figure 57

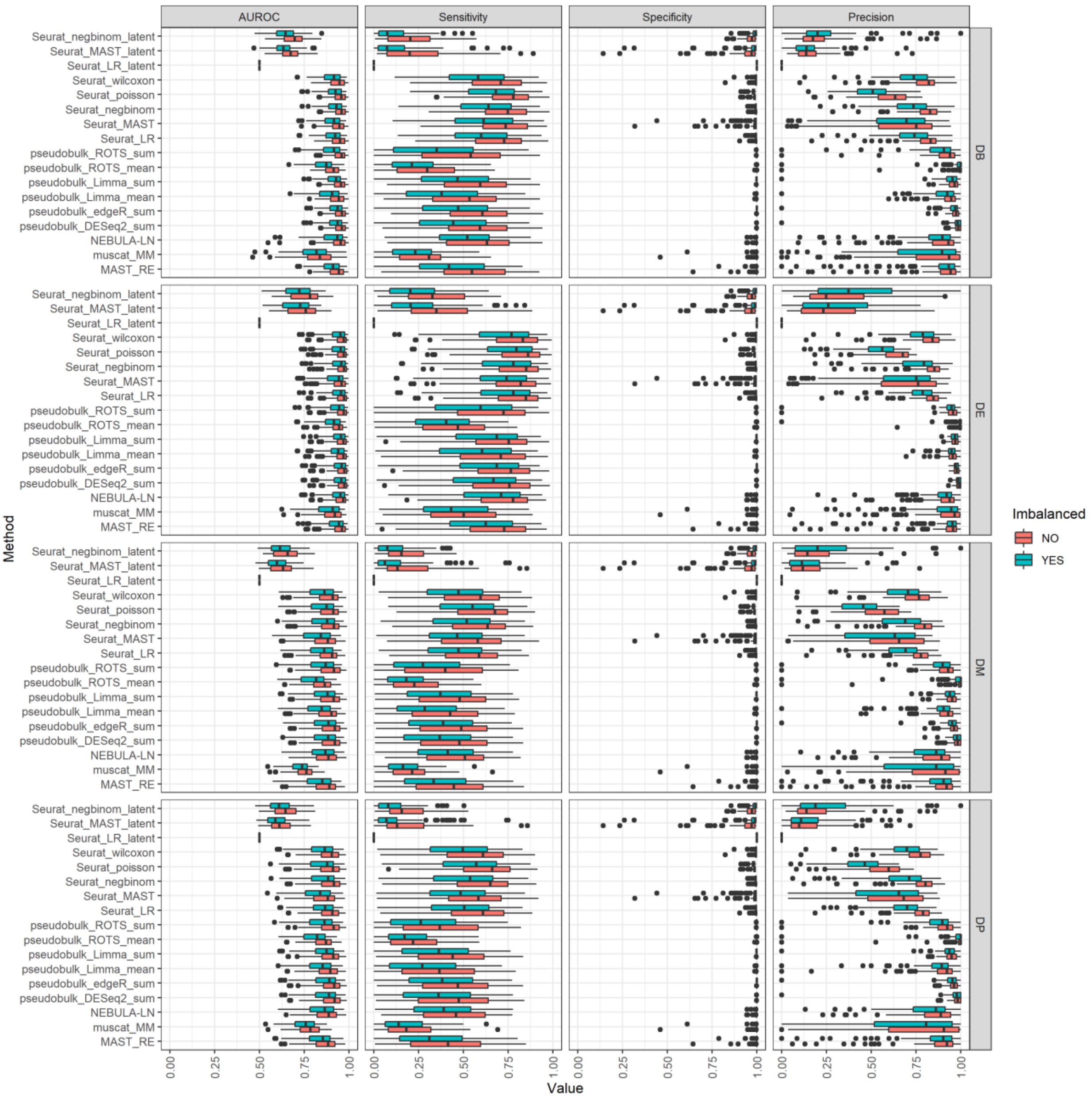


**Supplementary Figure 57. Results of the cell-sample-extended reference-based negative binomial generative simulation grouped by whether the data sets were downsampled to generate an imbalance distribution of cells across the samples.**
